# Design principles of ROS dynamic networks relevant to precision therapies for age-related diseases

**DOI:** 10.1101/647776

**Authors:** Alexey Kolodkin, Raju Prasad Sharma, Anna Maria Colangelo, Andrew Ignatenko, Francesca Martorana, Danyel Jennen, Jacco J. Briede, Nathan Brady, Matteo Barberis, Thierry D.G.A. Mondeel, Michele Papa, Vikas Kumar, Bernhard Peters, Alexander Skupin, Lilia Alberghina, Rudi Balling, Hans V. Westerhoff

## Abstract

The eminently complex regulatory network protecting the cell against oxidative stress, surfaces in several disease maps, including that of Parkinson’s disease (PD). How this molecular networking achieves its various functionalities and how processes operating at the seconds-minutes time scale cause a disease at a time scale of multiple decennia is enigmatic.

By computational analysis, we here disentangle the reactive oxygen species (ROS) regulatory network into a hierarchy of subnetworks that each correspond to a different functionality. The detailed dynamic model of ROS management obtained integrates these functionalities and fits *in vitro* data sets from two different laboratories.

The model shows effective ROS-management for a century, followed by a sudden system’s collapse due to the loss of p62 protein. PD related conditions such as lack of DJ-1 protein or increased α-synuclein accelerated the system’s collapse. Various *in-silico* interventions (e.g. addition of antioxidants or caffeine) slowed down the collapse of the system *in silico*, suggesting the model may help discover new medicinal and nutritional therapies.

## Introduction

Reactive Oxygen Species (ROS) are produced at various intracellular locations (1-3) inclusive of complex I of the Electron Transport Chain (ETC) in the inner mitochondrial membrane (4). ROS play three main roles: (i) “killing” in the immune response (Hall et al, 2013), (ii) signaling in cell differentiation and proliferation (Chaudhari et al, 2014; Juntilla et al, 2010; Kanda et al, 2011; Kinder et al, 2010; Le Belle et al, 2011; Lewandowski et al, 2010; Limoli et al, 2004; Martorana et al, 2018), and (iii) damage intracellular components including Complex I itself, sometimes leading to cell death (Kell, 2010). Around 0.1-2% of ETC electrons escape to form ROS. This percentage increases with increasing damage in Complex I (Li et al, 2013a). Excessive ROS is removed enzymatically, e.g. by superoxide dismutase (Afonso et al, 2007), or scavenged by various antioxidants (Oyewole & Birch-Machin, 2015). Should these processes fail, the removal of damaged mitochondria, called mitoptosis as it averts cell death (Allen et al, 2013; Cui et al, 2011; Fedorowicz et al, 2014; Ivatt & Whitworth, 2014; Wang et al, 2011) may be initiated (Brady et al, 2004; Kane et al, 2014; Koyano et al, 2014), or damaged mitochondria may be recycled into undamaged mitochondria via mitochondria-derived vesicles (MDVs). All these processes appear to be coordinated by a ROS-induced signalling network, with rather complicated cross-talk mechanisms (Joselin et al, 2012; Williamson et al, 2012; Xue et al, 2014). The most relevant involve p62 and its role in mitophagy, the Keap1/Nrf2 axis and NFκB for their role in modulating antioxidant responses, and the stress sensor DJ1 ((Ariga et al, 2013; Wang et al, 2013)). The dynamic response of this network to perturbations is determined by nonlinear interactions producing the functionality that is absent from its components. This means that the functionality emerges from interactions. Disease may thereby correspond to failure of the network to produce the functionality, e.g. both the robustness to the external perturbations and the failure thereof can be caused by various combinations of component failures (Alberghina & Westerhoff, 2005; Barabasi, 2012). Indeed, mistuning of ROS management and decreased ATP energy production has been implicated in aging (Grimm and Eckert, 2017) and in many diseases, including diabetes (Santiago & Potashkin, 2013; Soleimanpour et al, 2014), cancer (Matsuda et al, 2015) and neurodegenerative diseases (Joselin et al, 2012; Varcin et al, 2012; Williamson et al, 2012): The ROS management network contributes a *disease module* (del Sol et al, 2010; Loscalzo & Barabasi, 2011) to disease maps such as that of Parkinson’s disease (PD) (Fujita et al, 2014).

Affecting 1-3% of the population over 65 years old, PD is characterized by symptomatic motor dysfunction and alteration of the mood / reward system due to lack of dopamine secretion by dopaminergic neurons (Kruger et al, 2015). PD has been associated with diverse genetic and environmental factors. On the list of recessive PD-related mutations (Nalls et al, 2014), mutations in α-synuclein (Tran et al, 2014), and in Park 7 (DJ-1) (Wang et al, 2013) are prominent. With respect to environment and nutrition, PD risk correlates positively with pesticide exposure (Pfeiffer et al, 2012), and negatively with coffee consumption (Bjorklund & Cenci, 2010; Palacios et al, 2012; Postuma et al, 2012; van der Mark et al, 2014). On the anatomical level, the disease may be attributed to processes inside the dopaminergic neurons in the substantia nigra, to disruptions of communication between neuronal and glial cells, to inflammation, or even to the pathogenic spread of unfolded α-synuclein or tau proteins between cells (Lee et al, 2011; Walsh & Selkoe, 2016).

Network functionality may be reconstructed by translating network information into mathematical equations (Boogerd et al, 2005; Boogerd, 2007; Kolodkin et al, 2011) and using a mathematical model to identify ‘design principles’, i.e. network patterns responsible for functionalities. The robustness of these design principles to perturbations associated with disease may then be calculated (He et al, 2016; Kolodkin et al, 2010). The present paper implements this approach, with the additional aim of understanding the complexity of ROS management in the context of Parkinson’s disease (PD).

The Results section (and the corresponding Supplemental Materials) is divided into three chapters. In the first chapter we discuss the risks, risk-management principles, and a remaining liability in core models of increasing complexity, meanwhile identifying 5 design principles in the ROS network. In the second chapter we detail the dynamic model, validate it partially and re-examine how the design principles function in the complex integral network. The final chapter of the Results section uses the integral model to assess the time warp of aging, as well as possibilities for personalized therapies of Parkinson’s disease.

We will conclude that we now have a partly validated dynamic model that is able (i) to address aging and Parkinson’s disease as results of networked molecular processes, to (ii) solve the corresponding time warp, and (iii) to enable analyses towards individualized medicinal and nutritional therapies.

## Results

### Risk, management and a remaining liability

#### The risk: The positive feedback loop formed by impaired mitochondria and ROS constitutes a time bomb inherent in aerobic metabolism

The core of the ROS network (Fig. 1A, SM.A, Fig. SM.A.1Model A, so-called “no design”) consists of healthy mitochondria, damaged mitochondria producing ROS, and ROS damaging the healthy mitochondria. The dynamic version 1A of this (model 1A.cps) produced an explosion of both ROS and damaged mitochondria (SM.A, Fig. SM.A.1Model A). The positive feedback loop causing this effect, involving mitochondrial damage and ROS production, is unavoidable in cells using oxidative phosphorylation metabolism due to the proximity of FeS centres and molecular oxygen. To protect against this explosion, cells have implemented various mechanisms, as described in the following five design principles (Fig 1 and Fig 2A).

**Figure 1.**
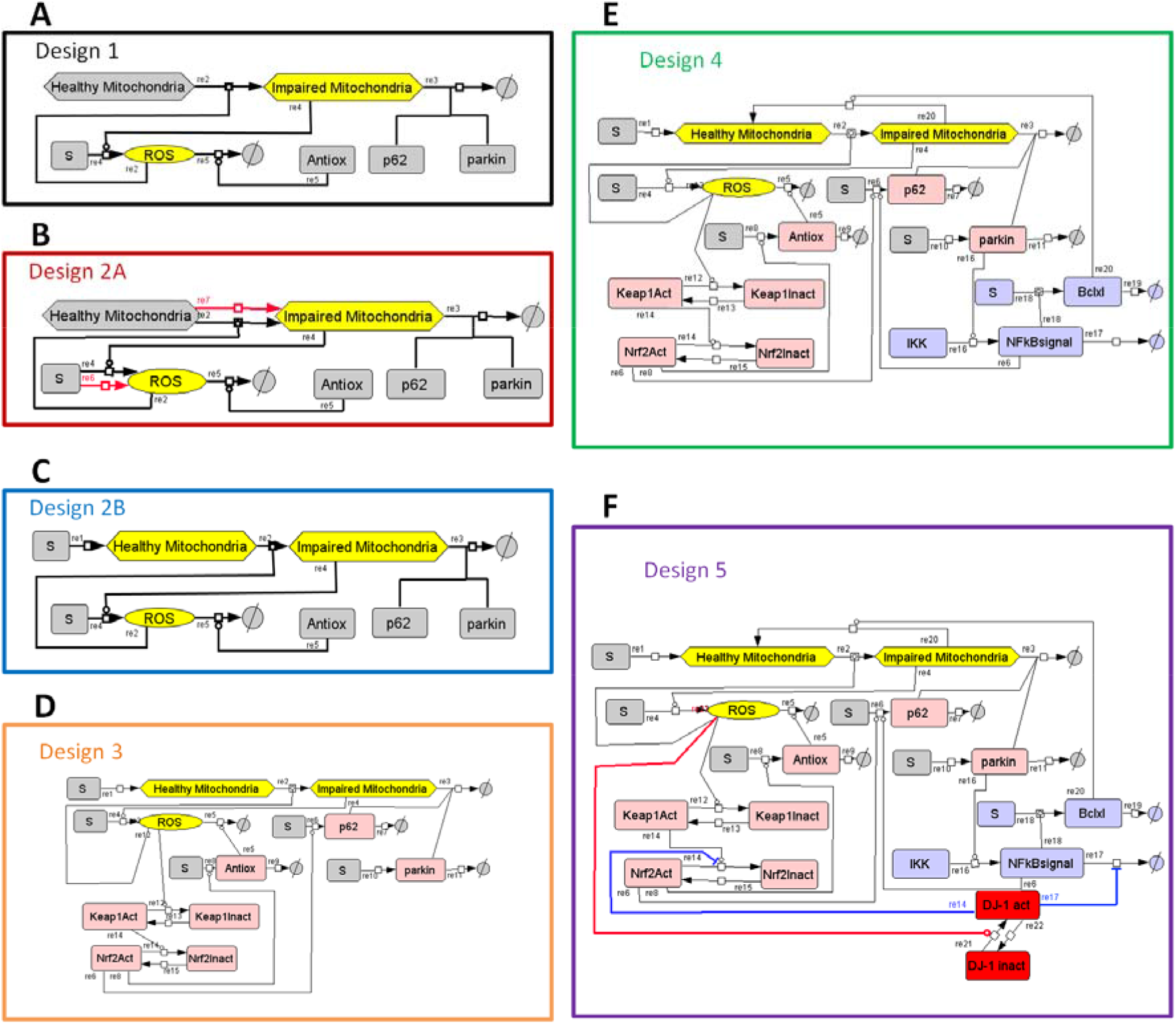
Network diagrams describing 5 principles of ROS management. Design 1 is comprised in Design 2 and in Design 3, Design 3 in Design 4, and Design 4 in Design 5, thereby forming a hierarchy of networks and corresponding models. (A) Design 1: Steady but not yet robust. Healthy Mitochondria present at a fixed concentration are damaged by ROS and thereby converted into Impaired Mitochondria by reaction 2 (re2). Impaired Mitochondria produce ROS (re4). Impaired Mitochondria and ROS hereby constitute a positive feedback loop as described in Fig S1-16. The node called Antiox comprises the total pool of all antioxidant response elements, i.e. both metabolites (e.g. glutathione) and enzymes (e.g. superoxide dismutase) catalyzing ROS removal (re5). p62 and parkin are required for mitophagy (removal of Impaired Mitochondria, re3) and are removed together with impaired mitochondria in the process (re3). (B) Design 2A: Steady but not yet robust. Reaction (re7) of basal ROS-independent mitochondrial aging (model B2A1), or reaction (re6) of basal ROS generation independent of the presence of Damaged Mitochondria (model B2A2), or both reactions (re7 and re6) (model B2A3) were added to Design 1. (C) Design 2B. Robust but not concentration (substances at variable concentrations are shown in non-grey boxes) and a reaction where they are synthesized at a constant rate (re1) has been added to Design 1. (D) Design 3. Homeostatic yet potentially oscillatory. Three negative feedback loops have been added to Design 2B (rose species on the diagram). The new variables Keap1 and Nrf2, in both active an inactive form, involve the Antiox, p62 and parkin species, which were hereto made variables, i.e. subject to synthesis and degradation. ROS oxidize and thereby inactivate Keap1 (re12), which is slowly re-reduced (re13). Active Keap1 shifts the Nrf2 balance towards inactive Nrf2 (re14). When active, Nrf2 activates the synthesis of both p62 (re6) and Antiox (re8), which causes breakdown of ROS (re5). Thus, ROS activates Nrf2, Antiox and p62, and inactivates Keap1 and itself: this forms a dual ROS-regulating negative feedback loop. Removal of damaged mitochondria causes a reduction in parkin levels, which reduces the removal rate of impaired mitochondria, which constitutes another negative feedback loop. (E) Design 4. Dynamically robust yet fragile vis-à-vis repeated challenges. Mitochondrial repair via the NF-κB signaling system (violet species on the diagram) has been added to Design 3: Parkin activates NFkB signaling via IKK (re16). NFκB signal activates the synthesis of Bclxl (re18) and p62 (re6). Bclxl activates the protection of mitochondria by salvaging Impaired Mitochondria (re20). (D) Design 5. Robust against repeated dynamic challenges. The DJ1 module (red species on the diagram) has been added to Design 4 as a ROS-sensor. DJ1 is a protein that may be present in two conformations: active (oxidized) and non-active (reduced). DJ1 activation is catalyzed by ROS (re21). When active, DJ1 inhibits removal of NFκB signal (re17) in sub-design 5.1, inhibits inactivation of Nrf2 (re14) in sub-design 5.2, or regulates both NFκB and Nrf2 signalling pathways in sub-design 5.3. Sub-design 5.3 is the complete version of Design 5, where, upon an increase of ROS concentration, DJ1 simultaneously activates both the antioxidant response and mitophagy (via Nrf2 and p62) and mitochondrial repair (via NFκB and BclXl).

**Figure 2.**
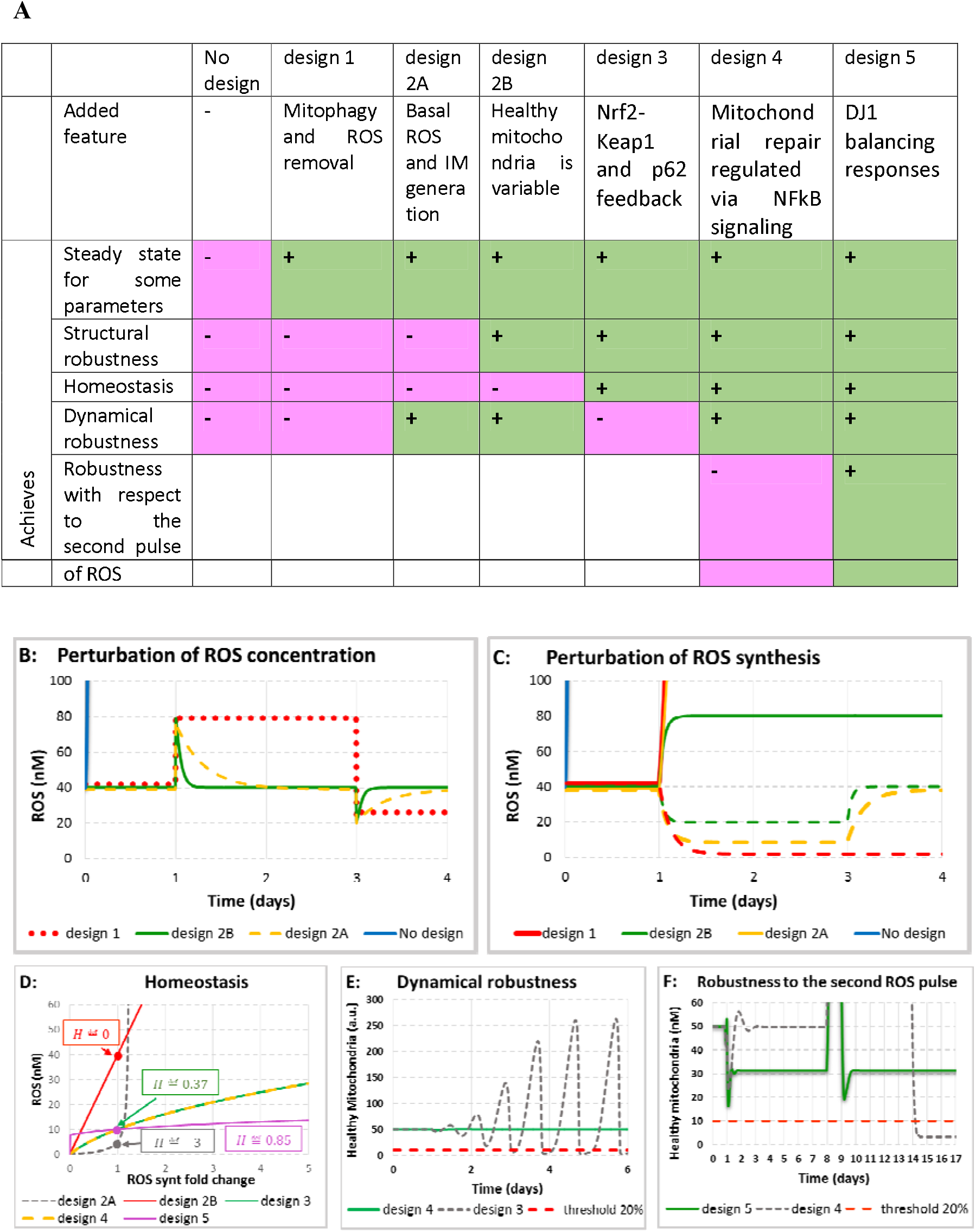
Emergent properties in 5 designs. **(A) Table summarizing added features and new emergent properties gained at every design.** **(B) The response to perturbations of ROS concentration in Design 1 (model B1), Design 2A (model B2A) and Design 2B (model B2B).** ROS concentration (in nM) is shown for the case of “no design” (solid blue line), which increases rapidly from 0 at time zero to over 100 nM within 1h. For numerical reasons, then the simulations was stopped, Design 1 (dotted red line), Design 2A (dashed yellow line) and Design 2B (solid green line). Both ROS injection (doubling of the initial ROS concentration at day 1), or a decrease in ROS level (the ROS concentration was decreased to 20 nM at day 3) was applied. All models are described in detail in Supplemental Materials SM.B1, SM.B2A and SM.B2B. **(C) The response to perturbations of ROS generation in Design 1 (model B1), Design 2A (model B2A) and Design 2B (model B2B).** ROS concentration (in nM) is shown for the case of “no design” (solid blue line), Design 1 (red lines), Design 2A (yellow lines) and Design 2B (green lines). The ROS generation rate constant was either increased 2-fold on day 1 (responses are shown in solid lines: solid red line for design 1; solid yellow line for design 2A; solid green line for design 2B) or first decreased 2-fold on day 1 and then returned back to the initial value on day 3 (responses are shown in dashed lines: dashed red line for design 1; dashed yellow line for design 2A; dashed green line for design 2B). All models are described in details in Supplemental Materials SM.B1, SM.B2A and SM.B2B. **(D) Emergence of homeostasis in Design 3 (model B3).** The steady state concentration of ROS (in nM) is shown against the fold change of the rate constant of ROS generation for Designs 2A (dashed grey line), Design 2B (solid red line), Design 3 (solid green line), Design 4 (dashed yellow line) and Design 5 (purple line). Homeostasis coefficients (H) for each design (shown in boxes) were computed at the point where ROS synthesis fold change was equal to 1. Model is described in Supplemental Materials SM.B3. **(E) Emergence of dynamic robustness in Design 4 (model B4).** The concentration of Healthy Mitochondria (in a. u.) is shown for Designs 3 (dashed grey line) and Design 4 (solid green line). The initial ROS concentration was perturbed (transient increase from 10 nM to 11 nM) at day 1. Dashed red line shows a hypothetical viability that corresponds to the threshold line dissecting 20% (10 a.u.) of the initial concentration of healthy mitochondria (50 a.u.). Model is described in Supplemental Materials SM.B4. **(F) Emergence of dynamic robustness vis-à-vis with respect to the second pulse of ROS in Design 5 (model B5).** The concentration of Healthy Mitochondria (in a. u.) is shown for Designs 4 (dashed grey line) and Design 5 (solid green line). The ROS generation rate constant was increased 15-fold on day 1, but, 3h before the increase of ROS generation, the NFκB signalling was increased 15-fold and the system reached a new steady state. On day 8 the ROS generation rate constant was decreased 15-fold causing the growth of healthy mitochondria. At the time point when the concentration of healthy mitochondria was near its peak value, the ROS generation rate constant was increased 15-fold for the second time. Model is described in Supplemental Materials SM.B5.

#### Its management: Five design principles controlling the ROS-network

##### Antioxidant response and mitoptosis (Design 1): steady but not yet robust

ROS activates antioxidant responses as well as mitoptosis (i.e. mitophagy specifically of damaged mitochondria). Inclusion of these two processes (Design principle 1) enabled the ROS network model to attain a steady state (Fig. 1A-B, Design 1; Fig. 2A, 2B and 2C; Fig.SM.B1.1Model B1).

However, this steady state was found only for a precise balance between the rate constant at which ROS was produced and the rate constant at which ROS was removed. Increasing the rate constant of ROS production caused an indefinite increase in ROS concentration (Fig. 2C; Fig. SM.B1.1A). When decreasing the ROS generation rate constant the concentrations of both ROS and damaged mitochondria dropped to 0 (Fig. 2C; Fig. SM.B1.1B). If, after the drop of ROS and the concomitant drop in damaged mitochondria, the ROS generation was increased back to the initial level, the concentrations of both ROS and damaged mitochondria at 0, the “perfect” state. Moreover, when exposed to a sudden injection of ROS, the system maintained the new ROS concentration rather than returning homeostatically to the pre-existing steady state (Fig. 2B; Fig. SM.B1.1C). These results demonstrated that Design 1 was both structurally and dynamically unstable.

##### Non-autocatalytic ROS and damage generation (Design 2A): steady but not yet robust

The assumption that ROS are only produced by damaged mitochondria is not realistic; there are other ROS producing processes, such as the Fenton reaction in the cytoplasm. Thus, we added a reaction of basal ROS generation (Design 2A, Model B2.A1; Fig 1B). Then, when ROS generation by damaged mitochondria was low because few mitochondria were left, ROS concentration was also low but not equal to 0, as it was still produced (at a lower rate) by other reactions that are independent of damaged mitochondria, as described in details in the Model B2.A1 at SM.B2.A.

When we decreased the total ROS generation in the new model, the concentration of ROS and of damaged mitochondria also decreased and reached a new non-zero steady state. When we returned ROS generation back to the initial level, the concentration of all species came back to the initial steady state (Fig 2C; Fig. SM.B2A.1A). The system also became stable to perturbations in the initial concentration of ROS. When we first increased and then decreased the initial concentration of ROS, both the concentration of ROS and the level of healthy mitochondria went back to the initial steady state (Fig 2B; SM.B2.A, Fig. SM.B2A.1B).

However, the stability of this system is limited. When the total ROS generation flux was increased 2 fold, the concentrations of both ROS and of damaged mitochondria abruptly increased (Fig 2C; Fig. SM.B2.A.1C).

An alternative way by which the system became partially stable emerged either when we took into account that mitochondria may get damaged also in the absence of ROS (Fig 1B, Design 2A; Model B2.A2), or when both the ROS production was possible in the absence of damaged mitochondria, and mitochondria can be damaged in the absence of ROS (Fig 1B Design 2A; Model B2.A3). Models B2.A2 (Fig SM.B2A.1D-1F) and B2.A3 (Fig SM.B2A.1G-1I) show a behaviour similar to that of model B2.A1 (SM.B2A).

One conclusion relevant for our further modelling approach is that zero concentrations of ROS and damaged mitochondria could be observed only for the case where we took into account neither basal ROS generation, nor basal mitochondrial damage (Model B1). Either of these two processes forbids the “perfect state” with zero concentration of ROS and damaged mitochondria (Fig 2C). In fact, we do not need to consider both basal ROS production and basal mitochondrial damage. On the one hand, basal production of ROS guarantees that there will be always a certain degree of mitochondrial damage, and, on the other hand, basal mitochondrial damage guarantees that there will be always a certain degree of ROS production. Thus, all models based on Design 2A would explode in terms of ROS and damaged mitochondria. Design 2A is insufficient for a robust behaviour.

##### Variable levels of healthy mitochondria (Design 2B): Robust but not yet homeostatic

In previous models (Design 1 and Design 2A) we considered the pool of healthy mitochondria to be unlimited. In reality, the cell has a limited capacity to synthesise new mitochondria and thereby to maintain the level of healthy mitochondria constant, hence independent of ROS damage. The sustained damage to mitochondria consequent to increased ROS should work to decrease the pool of healthy mitochondria. Therefore, we next assumed that the pool of healthy mitochondria is variable rather than constant. We also added a reaction of mitochondrial synthesis (Fig 1C, Design 2B) at a constant flux, in order to maintain the possibility that the system reaches a steady state.

The new model, which included Design principle 2B, exhibited a similar response to the perturbation of ROS concentration (Fig 2B; Fig SM.B2B.1C-D, Model B2B), but did no longer explode in terms of ROS levels when we increased ROS generation. When we increased ROS synthesis twice, ROS concentration increased twice as well and reached a new steady state (Fig 2C; Fig. SM.B2B.1E), while the concentration of healthy mitochondria was decreased around 2 fold (Fig. SM.B2B.1F). This effect makes Design 2B (as detailed in Fig SM.B2B) principally different from all previously discussed designs.

It is remarkable that doubling the rate constant for ROS synthesis caused the doubling of ROS concentration (Fig. 2C; Fig. SM.B2B.1E). Perhaps, paradoxically this did not affect the level of damaged mitochondria, but it did halve the level of healthy mitochondria (Fig SM.B2B.1F): it seems that variation in the level of healthy mitochondria is responsible for the regulation. The latter variation caused a temporary increase in the rate of mitophagy (Fig SM.B2B.1I), thus causing a strong decrease in the total mitochondrial concentration. This decrease of healthy mitochondria caused a decrease in the source for damaged mitochondria, explaining why the concentration of damaged mitochondria returned to initial level.

In Design 2B, we did not consider basal ROS synthesis or basal mitochondrial aging. Constant flux of synthesis of healthy mitochondria also helped to prevent the drop of concentrations of damaged mitochondria and ROS to 0. When the ROS level was decreased, the level of healthy mitochondria was increased, which then secondarily again increased the rate of formation of damaged mitochondria. The latter phenomenon prevented the network from reaching the condition with zero concentration of damaged mitochondria and zero concentration of ROS (Fig 2C; Fig SM.B2B.1A-B, Model B2B).

Summarizing, comparative analysis of Design 1, 2A and 2B (Fig 2C) revealed a Design principle 2: while an addition of basal ROS generation or basal mitochondrial damage or both (Design 2A) provides a dynamical, but not a structural robustness, the dynamic reduction of the abundance of mitochondria (Design 2B) ensures that both dynamical and structural robustness are achieved. Paradoxically, the dynamical reduction of the total mitochondrial pool may be essential to protect the cell.

In Fig 2D we plotted the steady state concentration of ROS versus the fold change of the rate constant of ROS generation for Designs 2A and 2B. While in Design 2A a small increase of ROS generation resulted in much higher increase of the ROS concentration, in Design 2B the 2-fold increase of ROS generation resulted in a doubling of the ROS concentration.

We computed the corresponding concentration control coefficient that shows how the increase of ROS generation affected the ROS concentration: 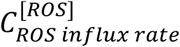. We then also quantified the homeostatic adaptation, which we propose to quantify as: 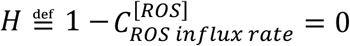. In Design 2A the concentration control coefficient was higher than 1 signifying H<0, i.e. negative (‘anti-‘)homeostasis, and in Design 2B the concentration control coefficient was equal to 1 signifying H=0, i.e. absence of homeostasis (Fig 2D).

##### The Keap1-Nrf2 module enables homeostasis through negative feedback (Design 3): Homeostatic yet potentially oscillatory

Design 1, 2A and 2B are based on a fixed rate constant of ROS consumption, which is determined by a fixed concentration of “Antioxidant Response” factors and a fixed first order rate constant of mitophagy (fixed concentrations of p62 and Parkin). We next added synthesis and degradation of p62, as well as synthesis and removal of “Antioxidant response”. Synthesis and degradation of “Antioxidant response”, p62 and Parkin were balanced in such a way that steady state concentrations of all species were identical to their concentrations in the previous Model B2B.

Then we increased the complexity by incorporating the Keap1-Nrf2 system, which modulates both mitophagy (by changing the p62 concentration), and an antioxidant response (by changing the concentration of antioxidant response species). When active, Keap1 binds to Nrf2, immobilises Nrf2 in the cytoplasm, and marks (through ubiquitination) Nrf2 for degradation. Nrf2 is a transcription factor that actively shuttles between nucleus and cytoplasm (Xue et al, 2014) and regulates the expression of p62 and genes responsible for an antioxidant response. Keap1 works as ROS sensor that regulates Nrf2 degradation and intracellular localisation. ROS oxidizes cysteine residues in the Keap1 molecule; Keap1 changes its conformation and becomes inactive (Williamson et al, 2012). Thus, the higher is the concentration of ROS, the less active is Keap1 and, consequently, the higher is the concentration of Nrf2 in the nucleus, where it induces the expression of p62 (activation of mitophagy) and the antioxidant response. Antioxidant response removes ROS and increased mitophagy removes damaged mitochondria, which produce ROS. Thus, negative feedback is produced: the higher is the ROS concentration, the higher is the removal of ROS. (Fig 1D, Design 3; SM.B3).

Addition of the Keap1-Nrf2 module hereby adds a new emergent property to the system: upon an increase of ROS generation, the ROS concentration first increased but then decreased again, presumably due to the negative feed-back loop which activated the antioxidant response and mitophagy and enabled homeostatic dynamic adaptation (Fig SM.B3.1A). Due to the combination of the positive (ROS inducing ROS via mitochondrial damage) and the negative (ROS reducing ROS via Nrf2-Keap1 signalling) feedback loops, the system exhibited transient oscillations of ROS (Fig SM.B3.1A), healthy mitochondria (Fig SM.B3.1A) and activated Nrf2 (Fig SM.B3.3) during the transition from one steady state to another in a certain range of the ROS generation rate constant.

The dynamic response of Design 2A showed an exponential increase of ROS concentration upon the increase of the ROS generation rate constant. Design 2B showed a linear increase of ROS concentration. In Design 3, the curve showing the variation of ROS steady state concentration with the variation of ROS generation rate was less than linear, and progressively less steep (Fig. 2D). The corresponding control coefficient was 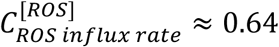. The reduction of this control coefficient to below 1, signifies homeostatic adaptation, which was equal to 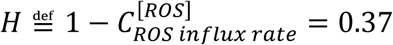. This number corresponds to the tangent to the curve on which the ROS concentration is plotted against the change of ROS generation, at the point where ROS generation was not changed yet (fold change of ROS generation was equal to 1). At the zone where ROS generation is substantially *de*creased, the curve was steeper, thus homeostatic adaptation was lower, and in the zone where ROS generation was high, the homeostatic adaptation was higher as well.

This can be summarised in the Design principle 3: The Nrf2-Keap1 feedback provides homeostatic adaptation. Thus, xenobiotics (like coffee, which besides caffeine contains a complex mixture of various chemical substances) interfering with the Keap1-Nrf2 system may affect the response to oxidative stress by affecting homeostasis.

However, the negative feedback loop of Design 3 introduces a problem. Designs 2A and 2B were robust against the perturbations of ROS concentration. Upon injection of ROS, the concentration of ROS, as well as all other species returned to their initial steady state level (Fig 2B). This is not anymore the case for Design 3. When ROS concentration is increased, e.g. on 10%, the network started oscillating (Fig 2E; Fig SM.B3.2). During oscillations, the concentration of healthy mitochondria swept below a viability threshold that we took corresponding to 20% of the initial concentration of healthy mitochondria (Fig 2E; Fig SM.B3.2). We imagine that a reduction of ATP formation capacity by 80% should be lethal for the cells.

##### Mitochondrial repair via NFκB signaling (Design 4): robustness against the ROS injections yet fragile vis-à-vis repeated challenges

As shown above, mitophagy of damaged mitochondria averts excessive ROS generation thus preventing the cell from ROS-induced damage that might lead to apoptosis (Brady et al, 2004). Mitophagy, however, leads to loss of mitochondria, and mitochondria play a dominant role in the free energy transduction. Loss of ATP synthesis flux is expected to lead to a drop in ATP/ADP ratio and thereby compromise ATP requiring processes. Should the maintenance metabolism be compromised, the loss could lead to necrosis. There is a physiological mechanism where, instead of being degraded in mitophagy, damaged mitochondria are repaired (Dhandapani et al, 2005; Lingappan, 2018; Tamatani et al, 2000). We incorporated this mechanism into the next design (Design 4; Fig. 1E; SM.B4) where NFκB-signalling activates mitochondrial recovery via activation of the expression of Bclxl (Janssen-Heininger et al, 2000).

This Design 4 (Model B4) became robust against the injection of ROS (Fig 2E; Fig SM.B4.1). Immediately upon the increase in ROS concentration, the latter returned back to initial value (Fig SM.B4.1A). Healthy mitochondria followed these perturbations and did not sweep below a hypothetical viability that dissects 20% of initial concentration of healthy mitochondria (Fig SM.B4.1B). This can be identified as Design principle 4: addition of mitochondrial recovery via NFκB signalling provides a dynamic robustness to the homeostatic design.

NFκB signalling also helps to resist high rates of ROS generation. When ROS generation was increased 15 fold and NFκB signalling was not activated, the concentration of healthy mitochondria dropped below a hypothetical viability line ((Fig SM.B4.2B). However, healthy mitochondria were saved if NFκB signalling was activated 15-fold by 6h (Fig SM.B4.2C), or even 3h (Fig SM.B4.2D) prior the increase of ROS generation. When NFκB was increased, the damaged mitochondria started to recover into healthy mitochondria. The concentration of damaged mitochondria decreased, thus the ROS concentration decreased as well and the concentration of healthy mitochondria started growing. The increase of ROS synthesis stopped the increase of the concentration of healthy mitochondria. ROS concentration then increased again and healthy mitochondria dropped again. But, overall, the concentration of healthy mitochondria did not cross a viability line.

At this stage, we could conclude that activation of NFκB signalling is advantageous for mitochondrial functionalities. The plot of steady state concentration of Healthy Mitochondria versus ROS showed that for higher NFκB signalling, and for the same level of ROS production, a higher level of Healthy Mitochondria could be maintained (Fig SM.B4.2E).

However there is also a potential danger of the accumulation of healthy mitochondria in Design 4 (Fig SM.B4.3): when healthy mitochondria are accumulated due to high NFκB activity and ROS generation is suddenly increased, accumulated healthy mitochondria could serve as a substrate for the production of damaged mitochondria. Because the concentration of substrate is high, there is also a high rate of mitochondrial damage and ROS generation. This triggers a positive feed-back loop: ROS damage mitochondria and damaged mitochondria produce more ROS. When damaged mitochondria accumulate faster than they can be neutralized by mitophagy, sharp peaks of damaged mitochondria and of ROS are observed. A vicious cycle fed by the high initial concentration of healthy mitochondria then leads to a collapse of the whole system. To avoid this catastrophe, the rate of mitochondrial recovery should adapt to ROS concentrations proportionately.

##### DJ-1 as a ROS sensor that coordinates Nrf2 and NFκB signalling (Design 5): Robustness against dynamic repeated challenges

DJ-1 protein is a ROS sensor that coordinates NFκB signalling with the level of ROS. When oxidised by ROS, the conformation of DJ-1 is changed. When DJ-1 becomes active it activates the rate of mitochondrial recovery via NFκB in a ROS dependent manner. In parallel, DJ-1 amplifies ROS-induced Nrf2-Keap1 signalling, thus activating both mitophagy and antioxidant response. To examine the role of DJ-1, we added the DJ-1 module to Design 4 thereby obtaining Design 5 (Fig. 1F).

We studied 3 versions of Design 5 (SM.B5): (i) Design 5.1 (Model B5.1), where DJ1 regulated NFκB signalling only, (ii) Design 5.2 (Model B5.2), where DJ1 regulated Nrf2-Keap1 signalling only, and (iii) Design 5.3 (Model B5.3), where DJ1 regulated both NFκB and Nrf2-Keap1 signalling.

When healthy mitochondria had accumulated due to high NFκB activity and low ROS generation, and ROS generation was suddenly increased at the time point when the concentration of healthy mitochondria was near its peak value, in both Designs 5.1 (Fig SM.B5.1) and 5.2 (Fig SM.B5.2), the concentration of healthy mitochondria transiently swept below a hypothetical viability line (that dissects 20% of the initial concentration of healthy mitochondria), but quickly recovered back to its initial level. This demonstrated the limited robustness of the model with respect to the second pulse of ROS when NFκB or Nrf2-Keap1 signalling pathways are already active. When similar perturbations were applied to Design 5.3 (Fig SM.B5.3), the concentration of healthy mitochondria did not sweep below the viability line revealing that Design 5.3 is robust *vis-à-vis* the second pulse of ROS and that simultaneous regulation of both NFκB and Nrf2 signalling pathways should be advantageous under such dynamic conditions.

To understand whether robustness is due to summation of two regulatory mechanisms, we increased ROS generation in Design 5.3 not only 15, but also 30 fold, i.e. twice higher than the cases when each of two mechanism worked alone. Still the concentration of healthy mitochondria did not sweep below a viability line (Fig SM.B5.4). In the next computational experiment (Design 5 with regulation of both NFκB and Nrf2 signalling), the sensitivity of DJ1 to ROS was reduced 2 fold (Model B5.5). In spite of the reduced DJ1 activity (Fig SM.B5.5A), Design 5.4 exhibited a behaviour similar to Design 5.3, and was still the most advantageous when comparing with Design 4, Design 5.1 and design D5.2 (Fig SM.B5.5B).

The final Design 5 with both NFκB and Nrf2 regulated by DJ1 exhibited very strong homeostasis (Fig 2D; Fig SM.B5.6). The corresponding control coefficient was 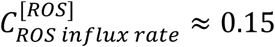 that signifies strong homeostatic adaptation, i.e. 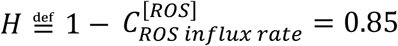.

Overall, we conclude that Design 5 provides robustness against a second injection of ROS and strong homeostasis. This should be due to DJ-1 and synergistic coordination of Keap1-Nrf2 and NFκB signalling.

Since DJ-1 coordinates mitochondrial recovery and amplification of Nrf2 signalling, and helps to bring dynamic homeostasis close to perfect adaptation, mutations in DJ-1 might lead to PD in cases where the network is challenged by large perturbations. When DJ-1 is present, the model seems perfect. However, we will now show that it may still fail when confronted with additional disease-related biological detail, such as ROS-dependent polymerisation of α-synuclein.

#### Remaining liability: Limited capacity to deal with α-synuclein polymerisation

In the previous designs, ROS activated p62 expression required for mitophagy. This process involved two negative feedback loops: (i) via Keap1-Nrf2 signalling, and (ii) via DJ1 signalling. However, ROS might also reduce mitophagy via α-synuclein polymerisation. Increased ROS concentration induces α-synuclein polymerisation. α-synuclein aggregates sequester p62, and lower p62 concentration is then responsible for the decline in mitophagy.

We added the α-synuclein module to Design 5 (Model B5) thereby obtaining Model C. (described in details in Fig SB.C.1). When α-synuclein was absent, Model C was at a steady state. With the addition of a constant source of α-synuclein at low concentration, oxidation of α-synuclein by ROS caused the formation of α-synuclein aggregates, which sequestered p62 and reduced mitophagy. Due to the reduction of mitophagy, this mechanism caused the increase of the concentration of healthy mitochondria in silico (Fig SM.C.2A). This hypothetical scenario could be attractive because it favours the increase of ATP production at the cost of somewhat elevated concentrations of ROS and damaged mitochondria.

When the concentration of α-synuclein was increased, the system did not reach a feasible steady state (Fig SM.C.2B). The concentration of damaged mitochondria started to grow constantly and the system exploded, demonstrating the potential danger of ROS-induced α-synuclein polymerisation.

We also checked the robustness of model C to a 10-fold increase of the ROS generation rate constant in the absence or presence of a source of α-synuclein at low concentration. Without α-synuclein, the system exhibited strong homeostasis (Fig SM.C.3). When a constant source of α-synuclein was added at low concentration, and the ROS generation rate constant had not yet been increased, the system attained a new steady state (Fig SM.C.4). However, once the ROS generation rate constant was increased in the new model that was already in a steady state, the steady state was no longer reachable (Fig SB.C.5).

To summarise, our in silico cell may tolerate (and perhaps even profit from) the reduction of mitophagy due to ROS-induced α-synuclein polymerisation, in the case of low ROS concentrations and low concentration of α-synuclein. However, ROS-induced α-synuclein polymerisation decreases the robustness against oxidative stress.

It is remarkable that in our simulations (SM.C) healthy mitochondria were not lost, because there was no mechanism in the model to stop mitochondrial synthesis even when damaged mitochondria and ROS grew to infinity. We expect that in live cells no more mitochondria could be produced with the accumulation of damage. Then the rate at which mitochondria are damaged should ultimately decrease in parallel to the decrease in the level of healthy mitochondria. This hypothesis will be examined in a more detailed model in the following sections.

### The detailed model

#### Detailed ROS-management model and validation by in vitro experiments

Based on literature data, we further increased the resolution of the ROS management model (Fig 3). All features of the five designs we identified and the α-synuclein module (SM.C) were maintained, but nucleus and cytoplasm compartmentalization, an mRNA layer for several proteins, and several additional species and interactions were added (Fig 3). In particular, in the new ATP module, reductive equivalents (RE) present at fixed levels in NADH+H^+^ provided electrons to the electron transport chain for the reduction of molecular oxygen either to water or to incompletely reduced oxygen (ROS). The former process consisted of a part that was coupled to the phosphorylation of ADP, as well as a part that was uncoupled of that phosphorylation. As we did not measure oxygen consumption and the oxygen levels were modelled as fixed, the latter process was considered irrelevant and is not reported on.

**Figure 3.**
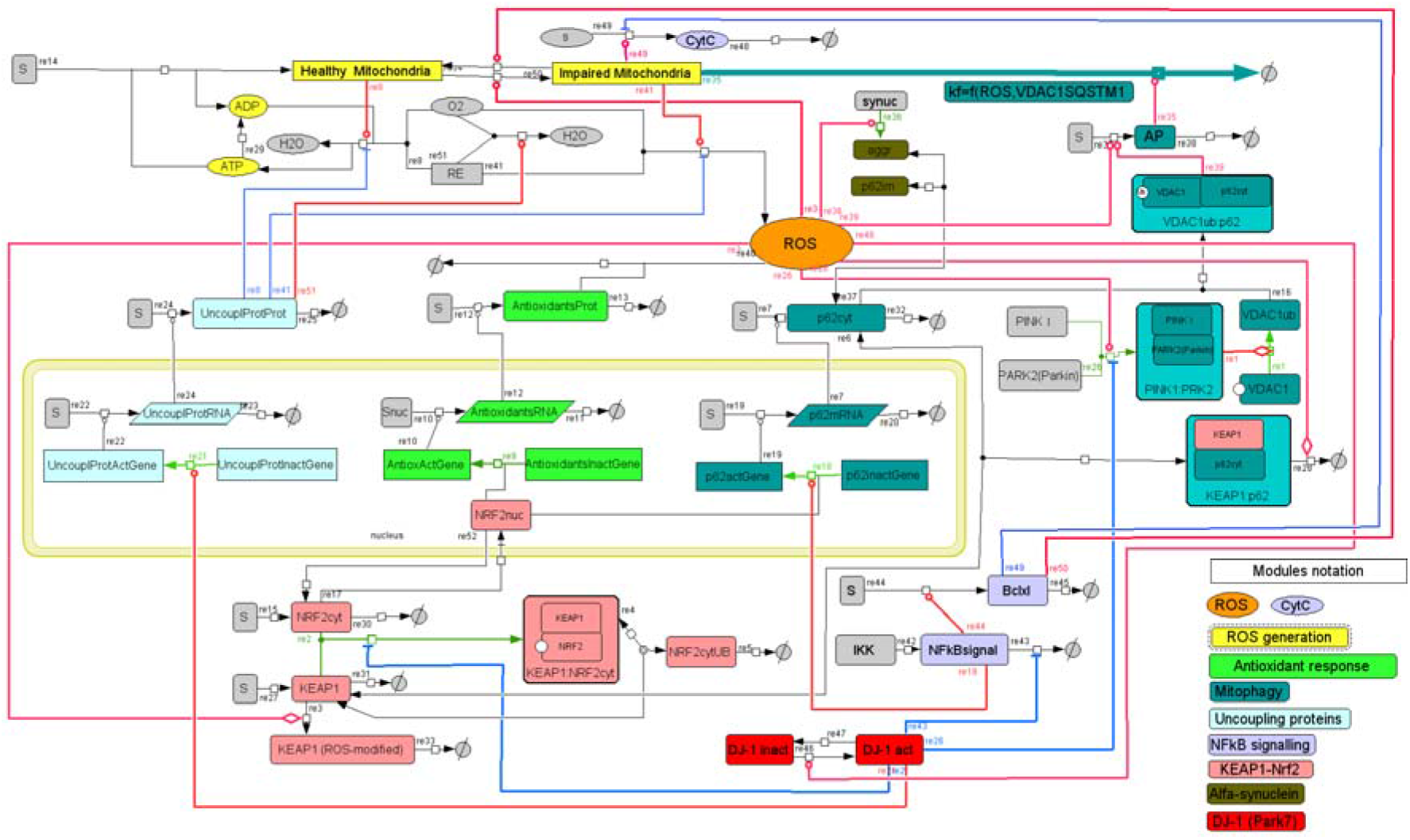
Network diagram of the detailed model (‘Model D’) of ROS management. The resolution of model C (Fig SM.C.1) was increased, i.e., the nucleus and cytoplasm compartments, an mRNA layer for several proteins, and several additional species and interactions were added as follows. **ATP module.** A constant source of reductive equivalents (RE) in the form of reduced NAD (i.e. NADH+H^+^), and a constant source of O_2_ drive three reactions: (i) in the reaction catalyzed by healthy mitochondria (re8), reduction of O_2_ to H_2_O is coupled to phosphorylation of ADP into ATP; (ii) in the reaction catalyzed by impaired mitochondria (re41), reductive equivalents are only used as a source of incompletely reduced oxygen species (ROS); (iii) in the reaction activated by uncoupling proteins (re51), O_2_ is reduced to H_2_O without phosphorylation of ADP into ATP. ATP is dephosphorylated back to ADP in reaction re29, which represents the overall net reaction of all ATP consuming reactions. **Uncoupling proteins**: The expression of uncoupling proteins is activated by active DJ1 in the ‘bipartite’ irreversible reaction (re21). Genes coding uncoupling proteins are present in active (transcribed) and inactive (silent) forms with the conserved moiety. The higher is the concentration of DJ1, the higher is the transcription of uncoupling proteins (re22). In turn, the higher is the concentration of uncoupling proteins mRNA, the higher is the rate of its translation into uncoupling proteins (re24). Uncoupling proteins inhibit the production of both ATP (re8) and ROS (re41) and activate the uncoupled respiration (re51). Since the concentrations of all its substrates (O_2_ and reductive equivalents) and products (H_2_O) are fixed, the reaction of respiration was omitted in the COPASI version of the models. A strategy similar to the modeling of ‘uncoupling proteins’ expression was used for the modelling of **Antioxidant response** and p62, for which active (transcribed) and inactive (silent) forms of genes were considered too. Activation of transcription would mean an activation of the ‘bipartite’ irreversible transition of a silent gene into its transcribed gene (re9 and re18). Binding of active (localized in the nucleus) fraction of NRF2 transcription factor which shuttles between the nucleus and cytoplasm (re17 and re52) facilitates the transition of a silent gene to its actively transcribed counterpart. When the gene is active, it catalyzes the transcription – (production of mRNA in re10 and re20). When the concentration of Mrna increases, the translation is activated (the production of proteins in re7 and re12). More detailed **mitophagy mechanism**: Pink1 binds to parkin in a ‘bipartite’ irreversible reaction (re26) and facilitates ubiquitination of VDAC1 in another ‘bipartite’ irreversible reaction (re1). The latter interacts with p62 (re16) to form a complex that, like ROS, facilitates the formation of apoptotic machinery (re39). This apoptotic machinery (called AP) catalyzes the reaction re35 in which impaired mitochondria are degraded. **Cyt C and NFκB**. Impaired mitochondria release Cyt C (re49). Apart from activating mitochondrial recovery (re50), Bclxl inhibits Cyt C release. When Cyt C exceeds a threshold, it induces cell death (not shown on diagram). **Keap1 module**. Keap1 transiently binds and then ubiquitinates and thereby facilitates degradation of both Nrf2 (reaction chain re2, re4, and re5) and p62 (re6 and re28; p62 is not released back; it is not a catalytic factor but a co-substrate). In these diagrams SBGN notation was used, i.e. –o for stimulation, -| for inhibition and – for co-reaction. <-> refers to reaction, which can be reversible [black double headed arrows], →irreversible [black arrows], ‘bipartite’ irreversible [green arrows] (this is specified in the Copasi files). ‘Bipartite’ irreversible refers to any case where a process has a forward and a reverse reaction that are not each other’s microscopic reversal, the one being affected by a third agent whilst the other is not. Reaction numbers are positioned at the origins of reaction arrows. The species with constant concentration (e.g., a constant source of substrates or a constant sink of products) are shown in grey color.

Our detailed mitophagy mechanism includes Pink1 (Morais et al, 2014; Rakovic et al, 2013) which activates Parkin E3 (Muller-Rischart et al, 2013; Sha et al, 2010) and facilitates the ubiquitination of VDAC1. The latter interacts with p62 and facilitates the formation of the apoptotic machinery. Besides activating Nrf2 and NFκB, DJ1 activates the expression of uncoupling proteins, which reduce mitochondrial transmembrane potential and inhibit the production of both ATP and ROS. This process was here modelled by the uncoupling proteins inhibiting ATP production, stimulating uncoupled respiration and stimulating ROS production. More details of NFκB signalling have been incorporated (Arena et al, 2013; Kodama et al, 2011). In addition, we have taken into account that impaired mitochondria release Cyt C. In addition to activating mitochondrial recovery, Bclxl inhibits Cyt C release. When Cyt C exceeds a certain threshold, it induces cell death. In the new model, Keap1 ubiquitinates and thus facilitates the degradation of both Nrf2 and p62. For p62, antioxidant response, and uncoupling proteins, active (transcribed) and inactive (silent) forms of the corresponding genes were considered, giving rise to the corresponding mRNAs and proteins (Fig 3). The resulting comprehensive model was called ‘Model D’.

Model D was then fitted to two independent data sets from *in vitro* experiments performed in different labs on different cell types (SM.D). First, we fitted the detailed model to the fold change in the relative concentrations of ROS (Fig 4A) and mRNAs (Fig 4B) upon an addition of menadione (100 μM) to HepG2 cells, starting at 1h steady state incubation with menadione. Menadione does not induce ROS immediately. A time delay of 35 min was therefore taken into account in the model simulation. The effect of menadione was modelled by assuming that menadione is transported into the cell and degraded, with a higher rate of the degradation inside the cell than in the extracellular media. In the cell, menadione induces ROS generation (re41 in Fig 3). This corresponds to the menadione-induced inhibition of mitochondrial complex I, enhancing electron leakage towards ROS generation.

**Figure 4.**
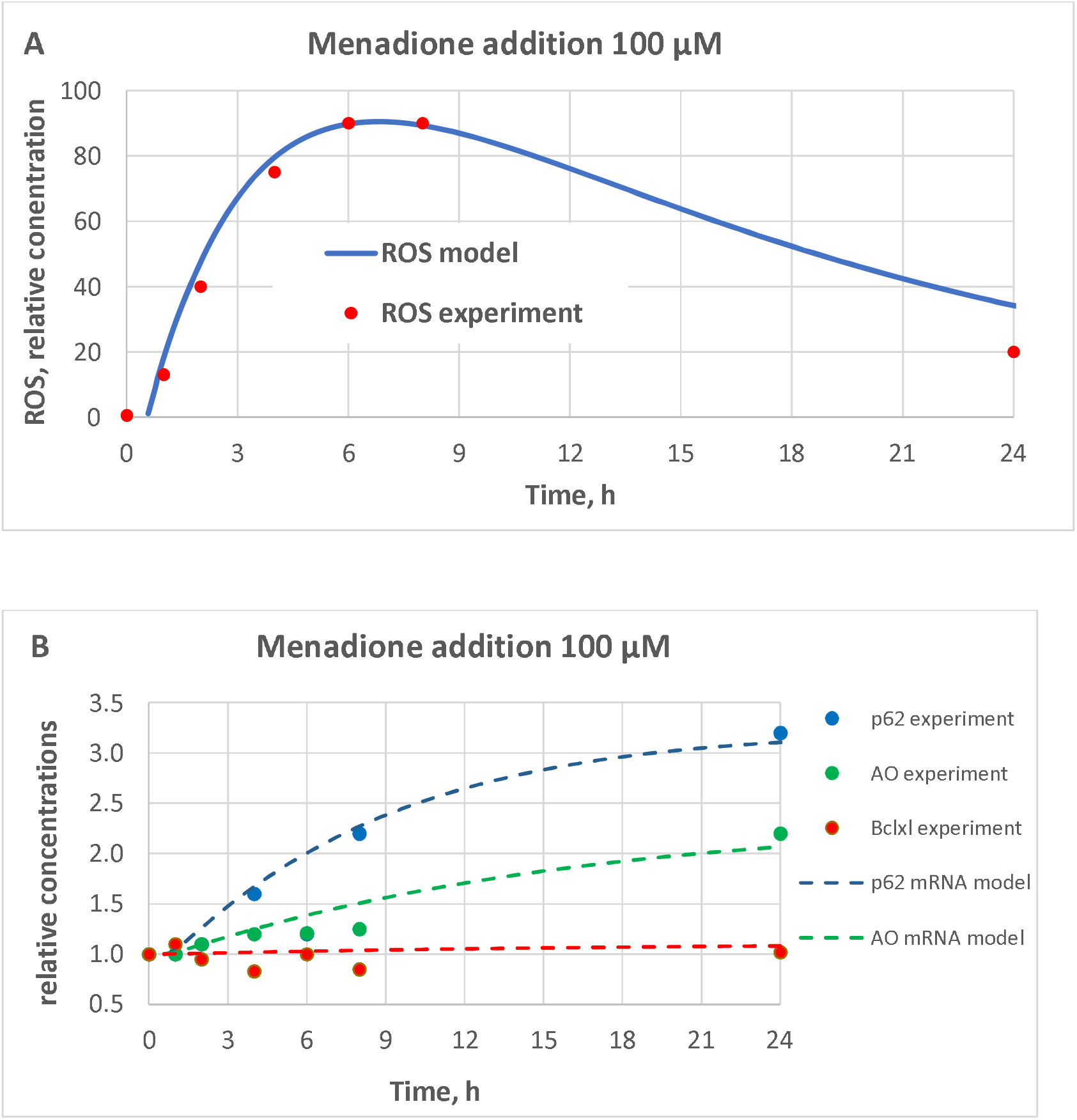
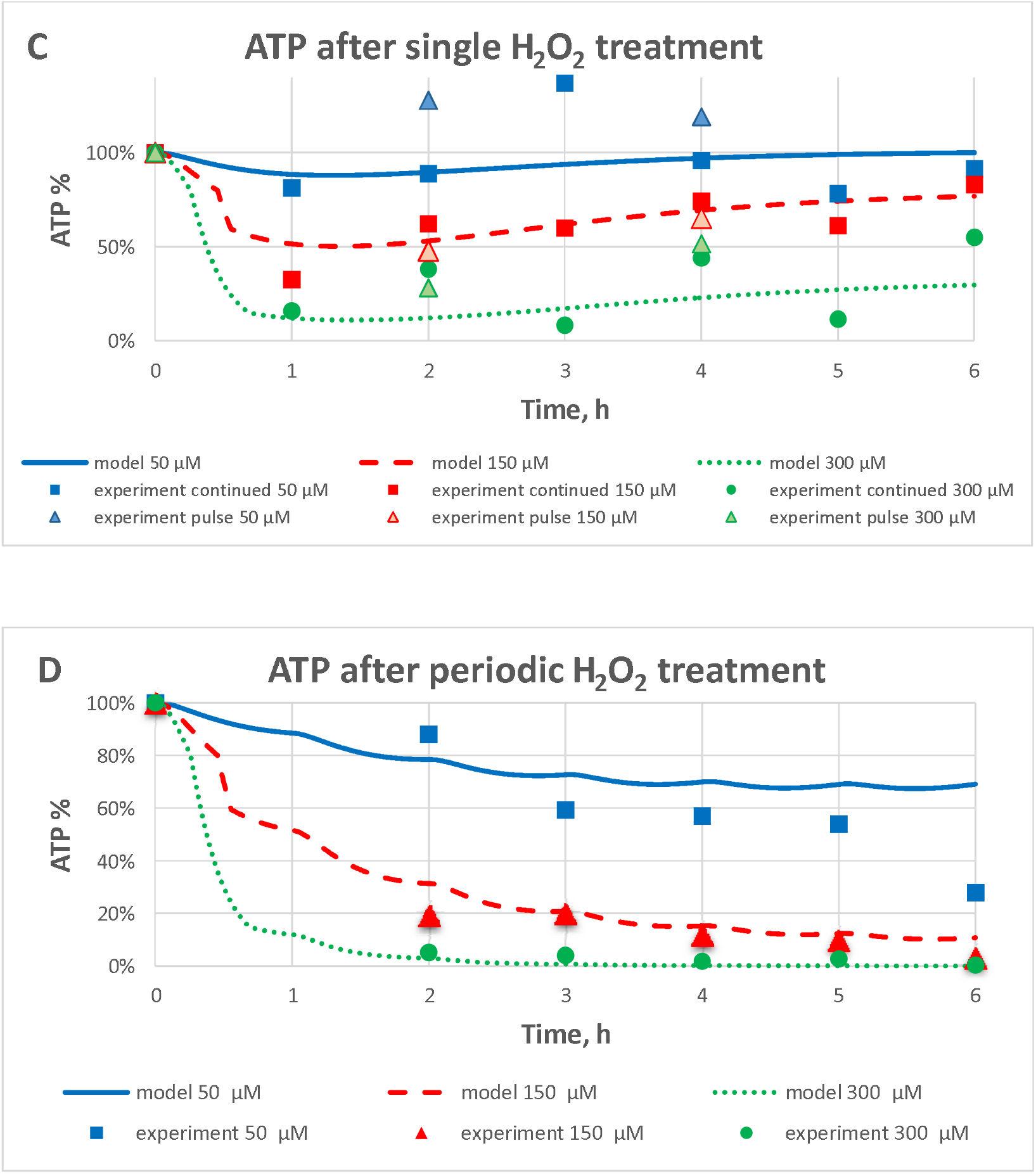
Validation of model D in terms of response to menadione and hydrogen peroxide. **A-B.** Fold change in relative concentrations of (A) ROS and (B) mRNAs upon an addition at time zero of menadione (100 μM) in model D to HepG2 cells, starting from 1 at the steady state before the addition of menadione. A time delay of 35 min was taken into account in the modelling. **C-D.** Changes in relative concentrations of ATP upon addition at time zero of H_2_O_2_ (50 μM, 150 μM, or 300 μM) to PC12 cells. (A) Time course of fold change concentrations of ROS. ROS predicted by the model (detailed model D) is shown by the blue line. Experimentally measured data are shown as red dots. The first point resides at a fold change of 1. (B) Time course of fold change in relative concentrations of p62 (blue), Antioxidant response using NQO1 as representative gene (green), and Bclxl (red) mRNA. (C) The concentration of ATP (% of its initial value) after one single addition to PC12 cells (at t=0) H_2_O_2_ 50 μM (blue), 150 μM (red), or 300 μM (green). This single treatment was either continued (squared dots) or pulse (triangle dots) treatment. In the pulse treatment, peroxide was washed away after 30 min by replacing the medium. In the continued treatment, no such action was taken. Separate experiments (Fig SM.D.2A) showed that peroxide degrades much faster than the time before washing. Thus, we did not differentiate between pulse and continued treatment during model fitting. (D) The concentration of ATP (% of its initial value) during periodic treatment with different concentrations of peroxide. H_2_O_2_ 50 μM (blue), 150 μM (red), or 300 μM (green) was added once per hour throughout the experiment, starting at t=0 hour. In all cases model D (corresponding to Fig 3) was used to calculate model predictions (see also Materials and Methods and Supplemental Materials SM.D). The perturbations by menadione and H_2_O_2_ were modelled as follows. Both menadione and H_2_O_2_ are transported to the cell in reversible reactions, and degraded both in the extracellular media and in the cell. Their rate of degradation in the cell is higher than in the extracellular media. In the cell, menadione induced ROS generation (re41 in Fig 3), that corresponds to the menadione-induced inhibition of mitochondrial complex I, enhancing electrons leakage for ROS generation. H_2_O_2_ catalyzes the production of other species, so-called ‘damage’, which correspond to ·OH radical formation and the accumulation of cellular damages by ·OH radical (DNA mutation, lipids oxidation etc.). ‘Damage’ is also removed (damage reparation). Similar to menadione, ‘damage’ causes an increase of ROS generation (re41 in Fig 3). When replacing this menadione module (A-B) with an H_2_O_2_ module (C-D), all parameters in the model are kept the same, except for those relating to ROS induction by menadione or H_2_O_2_, or the transport and degradation of menadione and H_2_O_2_. The model is publically available at FAIRDOMHub: Kolodkin A, Prasad Sharma R, Westerhoff HV. Mechanism-based detailed model of ROS management. FAIRDOMHub. 2019; http://doi.org/10.15490/fairdomhub.1.model.571.1. The model can be simulated online for FAIRDOMHub registered users, or the COPASI version of the model can be downloaded.

Then we studied another mechanism of ROS induction, through addition of H_2_O_2_, which was assumed to produce ·OH radicals damaging the electron transport chain. An additional species called ‘damage’ was produced and removed (damage repair). Damage caused the increase of ROS generation in re41 (Fig 3). When replacing the menadione module with an H_2_O_2_ module, all parameters in the model persisted. Although only the parameters of ROS induction by H_2_O_2_ were held adjustable, the model was cable to reproduce accurately the experimentally observed data of the time-dependent ATP change (Fig 4C-D and SM.D.2). In these experiments, PC12 cells were exposed to H_2_O_2_ (50 μM, 150 μM or 300 μM) in either one single treatment (Fig. 4C; Fig SM.D.2A,B,E) where H_2_O_2_ was added only once at t=0, or in a periodic treatment (Fig. 4D; Fig SM.D.2C,D,F) where H_2_O_2_ was added once every hour.

The fact that our comprehensive model D satisfied experimental data from both menadione and H_2_O_2_ titrations should provide some confidence in the model. On the other hand, a model of this complexity will need much more experimental validation and calibration in the future. Nevertheless, we decided to examine whether this somewhat validated model might resolve some important issues around aging and (Parkinson’s) disease.

#### Preconditioning and aging in the detailed ROS model

We exposed the detailed model D to several consecutive stress events (the increase of the ROS generation rate constant) and discovered that prior stress stimuli enabled the system to deal better with a subsequent stress (Fig SM.E.1A). This preconditioning was observed on the time scale of hours and was explained in terms of “memory” of the antioxidant response that was maintained at an elevated level after stress stimuli (Fig SM.E.1B). The system exhibited bi-stability. It was either quickly stabilized, or collapsed immediately upon a strong increase of the ROS generation rate constant (Fig SM.E.2). We were not able to observe any slow deterioration.

However, we should also consider that ROS damage the cell (e.g. accelerate the drizzle of mutations and oxidize lipids in mitochondrial membranes) and that the damaged cell produces more ROS. When incorporating this mechanism into the detailed model D and simulating the model’s behavior at the time scale of years, we observed the emergence of an at first gradual but subsequently sharp decline of the ATP concentration (Fig 5), which might be interpreted as ROS-related aging.

**Figure 5.**
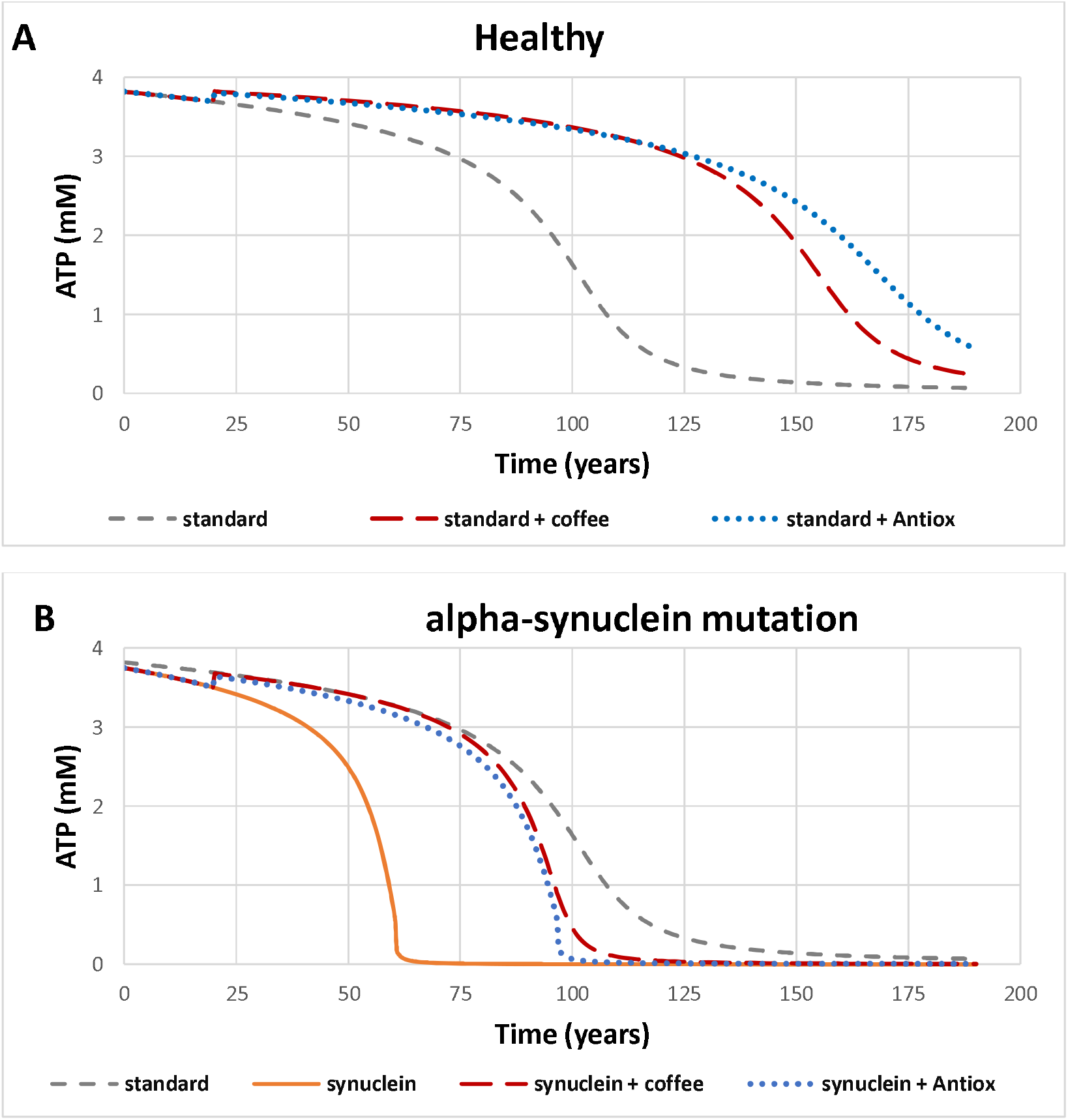
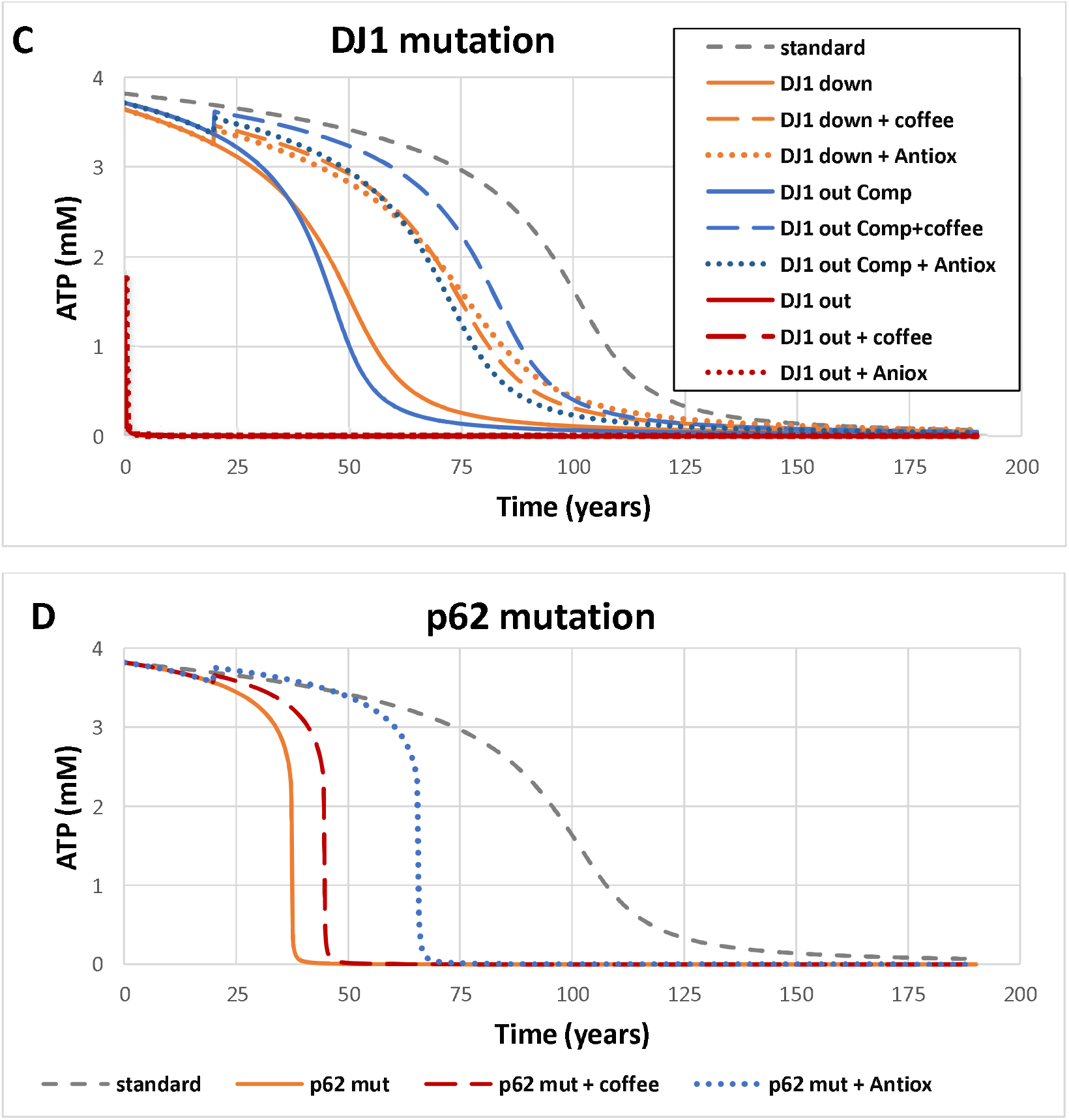
Personalized aging and medicine. The detailed model D (see Fig 3) was used for the simulations. Species representing ROS-induced damage accumulated in the model, which enhanced ROS production by Impaired Mitochondria. The treatment with coffee started at 20 years and was simulated by the 1.5-fold activation of Nrf2 nuclear import. The treatment by antioxidant started at 20 years as well, and was simulated as the 1.5-fold activation of the synthesis of antioxidant proteins. Aging (represented as the decline of ATP concentration thought to represent generalized failure of energetics) was simulated for 4 scenarios (A-D). A) Simulated ATP(t) in a healthy cell without any treatment (dashed grey line), and when treated with antioxidants, starting at 20 years as well, and simulated as the activation of antioxidant proteins synthesis (1.5-fold, dotted blue line), or with caffeine, started at 20 years and simulated by activation of the Nrf2 nuclear import (1.5-fold, dashed red line). B) Aging when α-synuclein source and thereby the rate constant of α-synuclein ‘aggregates’ formation (re36) is increased 2-fold without any treatment (solid orange line), or when being treated (like treatments in A) with either antioxidants (dotted blue line) or coffee (dashed red line), is compared with the standard aging of a healthy cell (dashed grey line). C) Aging in the presence of DJ-1 mutations, without any treatment (solid lines), or when being treated (like treatments in A), with either antioxidants (dotted lines) of coffee (dashed lines) is compared with the standard aging in a healthy cell (dashed grey line). Three mutated sub-versions were modelled: (i) DJ-1 mutation (‘DJ1 down’: orange lines), where DJ-1 activity is decreased 2-fold, (ii) DJ-1 mutation (‘DJ1 out’: red lines), where the concentration of total DJ1 protein is kept at almost 0, and (iii) DJ1 mutation compensated with the increased activity of Keap1-Nrf2 signaling, modelled by a 4-fold increase of the rate constant of Nrf2 production (‘DJ1 out Comp’: blue lines). Dashed lines refer to the corresponding cases treated as in C. D) Aging in the presence of a p62 mutation in which p62 mRNA level are fixed and are not regulated by ROS, without any treatment (solid orange line), and when being treated (like treatments in A) with either antioxidants (dotted blue line) or coffee (dashed red line). All cases are compared with the standard aging in a healthy cell (dashed grey line).

#### The design principles and the detailed integral model

As described in Supplemental Material SM.F we checked how the five design principles identified above with the use of simplified models might continue to work in the detailed model. The first three design principles worked in a straightforward way: without mitophagy (Design principle 1) the system did not reach any steady state (Fig SM.F.1), without limiting the mitochondrial synthesis (Design principle 2) the structural robustness was lost (Fig SM.F.2), without Keap1-Nrf2 signalling (Design principle 3) the homeostasis was substantially decreased (Fig SM.F.3). Homeostasis was not lost completely, because other feed-back mechanisms, such as NFκB and DJ-1 signalling, were still working in the system.

The interpretation of Design principle 4 (Fig SM.F.4) and Design principle 5 (Fig SM.E.5) in the detailed model was less direct at first sight. However, when considering these designs in the context of ROS-related aging, the role of both NFκB (Design principle 4) and DJ-1 (Design principle 5) signalling pathways became obvious. When NFκB (Fig SM.G.1) or DJ-1 (Fig 5C) signalling was compromised, an earlier ROS-induced aging was observed.

The role of α-synuclein in the detailed model (Fig SM.G.1) was slightly different from the role discovered earlier in the simplified model C (SM.C). The negative role of α-synuclein (increasing ROS concentration) persisted. However, the positive role (increasing the concentration of Healthy Mitochondria) was no longer observed. With the use of the detailed model, we also detected a possible role of increased concentration of mis-folded α-synuclein in the development of early ROS-induced aging (Fig 5B).

Since the abovementioned design principles might be compromised due to various mutations, for example mutations leading to reduced NFκB or DJ-1 signalling or to increased formation of α-synuclein aggregates, we used the detailed model to simulate the effect of those mutations in the context of age-related diseases, such as PD.

### Towards model-based design of personalized PD medicine

We asked whether our detailed ROS-management model with its ROS-mediated aging, could reproduce in silico, and thereby help us understand some intriguing observations reported in the literature. These include time spans much longer than the life times of the molecules engaged, the effect of various individual genes, the effects of diet, such as coffee and antioxidants, as well as personalized effects of diet-gene combinations. We looked at these events from the perspective of Parkinson’s disease (PD). First, we confirmed that all components of our ROS model have a place in the PD map (Fujita et al, 2014). Perturbing those components, we were able to simulate several scenarios of PD development, corresponding to several hypothetical patients and related to the design principles identified above. Two PD scenarios were related to p62: either direct mutations affecting p62 functioning (Fig 5D), or sequestration of p62 by misfolded α-synuclein (Fig 5B) (Tran et al, 2014). Other scenarios were related to DJ-1 encoded by Park7 being mutated in some PD cases (Fig 5C) (Wang et al, 2013) and compared with standard aging (Fig. 5A).

In standard aging, the initially high level of ATP was maintained for some 50 years, after which a slow ATP decline followed (Fig 5A). This decline accelerated after the age of 75 years to drop below 50% at around 100 years. At early age, patients without excessive inherent ROS generation but with either a substantial increase of α-synuclein (Fig 5B, orange line), or with a reduced DJ-1 functionality (Fig 5C, orange line) behaved similarly to the standard aging, however in an accelerated way. High concentration of ATP was maintained for only around 50 years. Assuming that ATP levels need to exceed 50% for viability, the virtual patient with the mutation in DJ-1 would not be viable, which may explain why a DJ-1 loss of function mutation has not been observed in the human population (Fig 5C, red line). However when the DJ-1 knockout was compensated with an increased Nrf2 activity (Fig 5C, blue line), the behaviour was similar to that of a DJ-1 knockdown. The effect of the p62 mutation was more severe, the concentration of ATP being maintained for only around 30 years (Fig 5D, orange line).

From the previous chapter, we might expect that an activation of Nrf2-Keap1 signalling (e.g. by caffeine and other coffee components) might protect from oxidative stress. Our simulations confirm that a PD-related collapse due to oxidative stress might be delayed under coffee-based or antioxidant treatment (Fig 5A). Thus, our detailed model reveals a network mechanism through which diet may play a PD protective role.

Our simulations also demonstrate that our virtual patients with increased levels of α-synuclein could be helped equally well by coffee and antioxidants (Fig 5B). Patients with the DJ-1 knockout compensated by Nrf2 signalling were affected more strongly by coffee (Fig 5C). The patient with p62 mutations was sensitive to antioxidants more than coffee (Fig 5D). Thus, inter-individual variations may cause disease variability between individuals as well as variability in therapeutic effects. Taking into account the possibility to identify the group of patients that would respond to therapy, coffee – based PD medicine could be more promising than the population or randomised trial studies where negative correlations between PD and coffee consumption have been observed (Bjorklund & Cenci, 2010; Palacios et al, 2012; Postuma et al, 2012; van der Mark et al, 2014).

## Discussion

Our models demonstrated that both mitochondrial recovery and mitophagy may avert ROS-induced cell death. This is related to several design principles that we identified and explained using our dynamic sub-models: (i) mitophagy enables steady state (Design principle 1), (ii) the dynamically variable concentration of mitochondria ensures that steady state is structurally robust against fluctuations in ROS generation (Design principle 2), and (iii) mitochondrial recovery via NFκB signalling serves robustness against increased ROS production. Paradoxically, a high rate of mitochondrial recovery is not always beneficial, but harms the cell if ROS generation suddenly fluctuates (see Design principle 4).

We have shown that mitochondrial recovery and mitophagy should also be coordinated with the antioxidant response. We have identified roles of Nrf2-Keap1 (Design principle 3) and DJ-1 (Design principle 5) in this coordination. The Nrf2-Keap1 system works as ROS sensor and forms the first line of defence by activating the antioxidant response and mitophagy (Bhatia et al, 2019; Lewis et al, 2015). DJ-1 is an additional ROS-sensor that amplifies the activity of Nrf2-Keap1 signalling and coordinates it with mitochondrial recovery (Ariga et al, 2013). Our models predicted that DJ-1 upregulation increases the cell’s robustness, whereas its downregulation should sensitize the system to oxidative stress. Indeed, some cancers are associated with upregulation of DJ-1 (Li et al, 2013b), while some cases of neurodegeneration are related to DJ-1 mutation or downregulation (Wang et al, 2013). It is remarkable that the same components are oppositely mistuned in opposing diseases: in cancer the cell survives elevated ROS, while in neurodegeneration the cell dies from ROS. Taking into account that DJ-1 is localised mostly in the mitochondria, but Keap1 is localised in the cytoplasm, one could foresee the importance of adding spatial aspects to the complexity of ROS-management. Here we have done this in the simplest way, i.e. by discussing three compartments, i.e. nucleus, cytosol and mitochondrion.

When we increase the resolution of model B5 to obtain the detailed model D, dynamic homeostasis emerges: several consecutive pulses of increased ROS generation (mild oxidative stress), “train” the ROS management system to deal with subsequent larger stresses (Yun & Finkel, 2014). This may explain several paradoxes reported in the literature, for example, those related to the observations that antioxidants may exhibit a hormesis response (Shore et al, 2012) and that, in some cases, antioxidant therapies disappoint clinical experience in cancer treatment (Bjelakovic & Gluud, 2007).

Our modeling results are also compatible with previous reports (Cloutier et al, 2012; Cloutier & Wellstead, 2012) demonstrating how the sequence of oxidative stress events may lead to the development of PD via a vicious cycle. For example, when rats were exposed to three pulses of paraquat (PQ) in order to imitate oxidative stress, the second addition allowed the system almost perfectly to compensate for the stress (adaptation), whilst the third pulse of PQ caused again a larger effect (and ultimately perhaps system collapse) (Finnerty et al, 2013). The authors explained these phenomena by bi-stability (Cloutier et al, 2012; Cloutier & Wellstead, 2012). Our models also contain bi-stability and may allow a similar interpretation, the first pulse of PQ leading to quick adaptation via activation of an antioxidant response and mitophagy that make the system more tolerant to the consecutive mild stress, but with the third pulses of PQ exceeding the threshold of the accumulated ROS-induced damage with the system then collapsing.

In several simulations, we observed that activated Nrf2 oscillated transiently (e.g. model 3). This correlates with literature data (DeNicola et al, 2011) showing that Nrf2 undergoes autonomous frequency-modulated oscillations between cytoplasm and nucleus. Oscillations occurred when cells were stimulated at physiological levels of activators, they decreased in period and amplitude and then evoked a cyto-protective transcriptional response. According to the data shown in (Xue et al, 2014), Nrf2 might be activated in cells without a change in total cellular Nrf2 protein concentration. In our model, an activation of Nrf2 is also related mostly to the shift between nuclear and cytoplasmic localization.

Metabolic reprogramming, due to a shift to nutrients that increase ROS production, change the rate at which mitochondria generate ROS and ATP; nutritional shifts may vary the NADH/ATP/ROS ratio as well as the sensitivity to oxidative stress (Polyzos et al, 2019). Such metabolic reprogramming corresponds to the changes in the operating regime of our blueprint model, for example where we use the NADH redox level as our model’s input and the ATP hydrolysis work load as one of our model’s output. Addition of this extra complexity is yet another potential application of our blueprint model.

Our detailed model is an interesting example where a slow aging process emerges from quick processes in the network. It is an example of multiscale modelling, where the minutes’ time scales of molecules are shown to be relevant for a hundred year process in a multicellular organism. It further shows how apparently minor differences in persons’ genomes (or expressomes) that do not affect an unchallenged function, may become crucial when that functions becomes challenged by oxidative stress. To be able to achieve this, we built our models *ab initio*, i.e. starting from the molecular cell physiology of the response to oxidative stress and increasing the complexity of the network step by step. Adding every new level of complexity in a domino approach, enabled us to identify design principles of ROS management. We feel that this initial simplicity was essential for this identification to succeed. At the same time, our most complex model, which still comprised these design principles, became a blueprint model into which the information from the available disease maps could be projected. Overlaying our blueprint model with the PD map (Fujita et al, 2014), we obtained several PD-related patient-specific models.

Yet, the findings of this paper should be taken with appropriate reservations. The literature knowledge it was based on is still limited and the principles discovered are therefore principles of the types of network that might operate *in vivo*. At the same time, our models only contain a small fraction of all that is known about the networks they deal with. But they are blueprint models and readily allow insertion of more and more precise detail. When examining a particular patient with single nucleotide polymorphisms (SNPs) in a set of relevant genes, such particularization will become important and possible. We entertain the concept that an individual’s genome may be examined for SNPs in pro-and anti-apop-and mitop-totic factors, the implications then becoming predictable by more detailed versions of our models.

Whether the design principles we identified do operate more generally, should now be a matter of experimental validation. In a way this paper merely helped to recognize what important principles are there to be validated in the near future. We did present two lines of model validation, but much more such validation is needed. Our examples merely serve to show how validation will now be possible.

Even before their precise validation, our calculations may serve as case studies connecting data-driven biomedical disease maps with systems biological dynamic models built *ab initio*. They then serve as proofs of concept showing how personalised medicine might benefit from such connections and how fundamental design principles studies may inspire practical biomedical questioning.

In conclusion, we showed how to deal with complex networks and understand their functioning, by principles of function. We implemented this for PD and showed several examples of how Nature and Nurture interactions and personalization may be studied.

## Materials and Methods

### Model building

Model diagrams (e.g. Fig. 1 and Fig. 3) were generated using CellDesigner (v4.4; Systems Biology Institute, http://celldesigner.org/index.html), a graphical front-end for creating process diagrams of biochemical networks in systems Biology Markup Language (Kitano et al, 2005). CellDesigner-generated models were transferred to COPASI (v4.6, build 32) (www.copasi.org), which is another Systems Biology Markup Language-compliant programme, but with a wider variety of analysis options. For each reaction, a kinetic term can be included, detailing the interactions between the species. A number of the parameter values used within the model were fitted to the known biological behaviour of the system, while maintaining these parameters within previously determined biologically realistic bounds.

### Model-stability checks

Two procedures to check model stability were used: (I) We took the final state as the initial state (Copasi enables one to do this with a single click) and then ran the model again and checked that then nothing changed with time. Subsequently, the initial condition of some variables were changed somewhat (e.g. by a factor of 2) and a time calculation was started, again checking that the same final steady state was reached. (II) We ran the steady state mode of Copasi and then activated its stability analysis, checking that none of the eigenvalues were reported to be positive.

### Models parameterisation

All parameters were set in the physiological range, and maintained in the physiological during all model adjustments and simulations. Several parameter values were identified from the literature. Other parameters were initially choosing in the physiological range and adjusted to achieve the biologically meaningful steady state. When validating the model by the experiments with menadione-induced oxidative stress the parameters were adjusted to reproduce the experimental data. When validating the model with the data on the changes of the ATP concentration induced by H_2_O_2_, the only rate constant of (i) H_2_O_2_ transport to the cell, (ii) H_2_O_2_ degradation, and (iii) H_2_O_2_-induced ROS generation were adjusted. All other parameters were persisted as in the experiments with menadione.

### Model sharing (FAIRDOM)

The main model of ROS management (detailed model D) is publically available at FAIRDOMHub: Kolodkin A, Prasad Sharma R, Westerhoff HV. Mechanism-based detailed model of ROS management. FAIRDOMHub. 2019; http://doi.org/10.15490/fairdomhub.1.model.571.1. The model can be simulated online for FAIRDOMHub registered users. The COPASI version of the model can there be downloaded.

### ATP experiments (with H_2_O_2_ perturbations)

#### Cell cultures and treatments

PC12 cells were maintained in Dulbecco’s Modified Eagle medium (DMEM) supplemented with 10% fetal bovine serum, 5% heat-inactivated horse serum, 2 mM L-glutamine, 100 µg/ml streptomycin, 100 U/ml penicillin in a humidified atmosphere of 95% air 5% CO_2_ at 37^°^C, as previously described (Martorana et al, 2018). All cell culture reagents were purchased from EuroClone (Milano, Italy).

#### Cell viability

Cell survival was assessed by the MTT assay as described in (Amara et al, 2015). Briefly, PC12 (5000 cells/well) were plated onto 96-well plates (Euroclone) pre-coated with poly-L-lysine. Tetrazolium salts (0.5 mg/ml) were added to the culture medium during the last 4h of treatment. After incubation, 100 µl of MTT solubilization buffer (100 µl) was added to each well for 1h. The absorbance of samples was measured at a wavelength of 570 nm (700 nm reference wavelength) with a Microplate Reader (BioRad). MTT conversion levels were expressed as a percentage of control.

#### ROS analysis by Flow Cytometry

Determination of intracellular levels of total ROS was carried out by flow cytometry using 2’,7’-dichlorodihydrofluorescein diacetate (DCH2FDA, Thermo Fisher Scientific), as previously described (Amara et al, 2015). PC12 (2×10^5^ cells/well) were plated in 6-well plates (EuroClone) pre-coated with poly-L-lysine (0.1 mg/ml). DCH2FDA was added during the final 30 minutes of treatment. Cells were then washed with PBS, harvested with 0.08% Trypsin and analyzed by FACS (FACScan Becton-Dickinson, San Jose, CA) using the Cell Quest Software (BD Bioscience). Fluorescence was measured on 1×10^4^ cells and data were analyzed by using the Flowing Software 2.5.1 (Turku Centre for Biotechnology, University of Turku, Finland).

#### ATP determination

PC12 cells were plated into 6-well plates (7×10^4^ cells) and exposed to H_2_O_2_ for the indicated times. Cells were then lysed by using lysis buffer and ATP activity was analyzed by using the Adenosine 5’-triphosphate (ATP) Bioluminescent Assay Kit (Sigma-Aldrich) according to manufacturer instructions. The light intensity was measured with a luminometer (Lumat LB9507, Berthold) in a 5s time period and expressed as relative light units/μg of protein.

#### Statistical analysis

All data are presented as the mean ±SEM of the number of independent samples in separate experiments, as indicated in the figure legends. Statistical analysis was performed by using GraphPad Prism 6.0 (GraphPad Software, La Jolla, CA, USA). All quantitative data were analyzed by one-way ANOVA and Dunnett’s multiple comparisons test for multiple treatments or by Student’s t-test for single comparisons (* p≤ 0.05, ** p≤ 0.01, *** p≤ 0.001 versus control (CTR)), as indicated in the figure legends. For morphology analyses, individual images of CTR and treated cells were assembled and the same adjustments were made for brightness, contrast and sharpness using Adobe Photoshop (Adobe Systems, San Jose, CA).

### Menadione experiments

In the experiment HepG2 cells were exposed to menadione and a consequent increased ROS intracellular concentration and via micro-array analysis of whole genome gene expression the fold change in the expression of p62, Bclxl, and NQO1 (the gene representing the antioxidant response) were measured. The experiment lasted 24 hours. For the measurement the following time points were chosen: 0.5, 1, 2, 4, 6, 8 and 24 hours after menadione addition.

#### Cell culture, menadione dose selection and ROS formation

HepG2 cells (ATCC, LGC logistics) were cultured in 6-well plates in the presence of minimal essential medium supplemented with 1% nonessential amino acids, 1% sodium pyruvate, 2% penicillin/streptomycin, and 10% fetal bovine serum (all from Gibco BRL, Breda, The Netherlands). The cells were incubated at 37°C and 5% CO_2_. When cells were 80% confluent, the medium was replaced with medium containing menadione (Sigma-Aldrich, Zwijndrecht, The Netherlands). As a solvent control, just medium was used. A non-cytotoxic concentration of menadione was selected based on a 3-(4,5-Dimethylthiazol-2-yl)-2,5-diphenyltetrazolium bromide (MTT) cytotoxicity assay with 80% viability after 24h exposure. In addition, using ESR spectroscopy, the final concentrations of 100 µM menadione was determined based on maximum oxygen radical formation at a non-cytotoxic dose. Radical formation in HepG2 cells was measured by ESR spectroscopy in combination with the spin trapping technique using 50 mM 5,5-dimethyl-1-pyrolline N-oxide (DMPO, Sigma-Aldrich). Protein oxidation was confirmed via a protein carbonyl assay and oxidative DNA damage via detection of this using the Fpg comet assay (Deferme et al, 2013).

#### Whole genome gene expression via analysis of microarray array data

cDNA was prepared using Affymetrix synthesis and labeling kits as described before (Affymetrix, Santa Clara) (Jennen et al, 2010). cRNA targets were hybridized on high-density oligonucleotide GeneTitan chips (Affymetrix Human Genome U133 Plus PM GeneTitan 24 arrays) according to the Affymetrix Eukaryotic Target Hybridization manual. The GeneTitan arrays were hybridized, washed and stained using the GeneTitan hybridization, wash and stain kit for 3’ IVT Arrays and GeneTitan Operating Software, and scanned by means of an Affymetrix GeneTitan scanner. Normalization quality controls, including scaling factors, average intensities, present calls, background intensities, noise, and raw Q values, were within acceptable limits for all chips.

Data from one hundred and twenty six arrays were obtained, and Robust Multi-array Averages (RMA) were normalized and re-annotated to produce custom CDF files using the array analysis tool (http://arrayanalysis.org/). In addition, 18,909 genes were analyzed for the number of absence calls in the three replicates per treatment. Genes that contained two or more absence calls within the three replicates for all the treatments as well as in controls were omitted from the data.

The data discussed in this publication have been deposited in NCBI’s gene expression omnibus (Edgar et al, 2002) and are accessible through GEO series accession number GSE39291: http://www.ncbi.nlm.nih.gov/geo/query/acc.cgi?acc=GSE39291

The intensities of the filtered data sets were log2 transformed, and subsequently, log ratios of treated versus controls were calculated. Differentially expressed genes (DEGs) for each experimental group were selected using the following criteria: (1) log2 ratio of −0.26 or >0.26 (i.e., absolute fold change of 1.2) for the average of the three replicates within the experimental group, (2) same direction of the log ratio for all replicates, (3) intensity of log2 values >6 for at least 2 out of 3 replicates, and (4) a p value of <0.05 determined using the student’s t-test. No FDR was used since it has been reported that reproducibility of microarray data is higher when criteria such as fold change are used (Guo et al, 2006). Gene expression was confirmed via real time PCR and correlated significantly for all tested genes being SOD1, SOD2, CAT, OGG1, CDKN1A, GPX1 and HMOX1 assay (Deferme et al, 2013).

## Acknowledgements

Alexey Kolodkin acknowledges funding from the Luxembourg BioTech Initiative and LCSB; Alexey Kolodkin and Andrew Ignatenko acknowledge the financial support from an FNR grant for the PhD project ROSIM; Hans V. Westerhoff thanks the EU, the BBSRC, EPSRC and NWO for extensive research support throughout many years, such as in grants BB/F003528/1, BB/C008219/1, BB/F003528/1, BB/G530225/1, BB/I004696/1, BB/I017186/1, BB/I00470X/1, BB/I004688/1, BB/J500422/1, BB/J003883/1, BB/J020060/1, and the EU-FP7 projects SYNPOL, EC-MOAN, NUCSYS, UNI-CELLSYS, ITFoM, BioSiM and EPIPredict. Matteo Barberis acknowledges the Systems Biology Grant of the University of Surrey and the Swammerdam Institute for Life Science Starting Grant of the University of Amsterdam. This work was further supported by grants from the Italian Ministry of University and Research (MIUR) SYSBIO-Italian ROADMAP ESFRI Infrastructures to Lilia Alberghina, Anna Maria Colangelo and Michele Papa; MIUR IVASCOMAR-National Cluster to Anna Maria Colangelo).

We cordially thank Nathan Price and Evangelos Simeonidis from ISB (Seattle), Nilgun Sahin from the VU University Amsterdam, Ewelina Weglarz-Tomczak from the University of Amsterdam and Bernhard Peters from the University of Luxembourg for highly influential discussions and support. We thank Stephan Gebel for his great help with the PD map.

## Author Contributions

AK, RPSh, HVW, RB, AMC, AI, NB, MB, VK, TDGAM, LA contributed to the development of network diagrams, model building, model parametrisation. AK, RPSh, AI and HVW contributed to dynamic model building and simulations. DJ, JB, AMC, FM and LA provided the experimental data and expert support. MP, AMC, LA, NB, AS and RB provided the expertise in ROS and neurodegeneration. AK, HVW, MB, NB and RB contributed to project supervision and to setting the direction of the research. AK and HVW wrote most of the manuscript, with significant help from AMC.

## Conflicts of Interest

The authors declare that there are no competing financial interests in relation to the work described.

## References

Afonso V, Champy R, Mitrovic D, Collin P, Lomri A (2007) Reactive oxygen species and superoxide dismutases: role in joint diseases. Joint, bone, spine: revue du rhumatisme 74: 324–329

Alberghina L, Westerhoff HV (eds) (2005) Systems biology: definitions and perspectives. Berlin: Springer, 408pp

Allen GF, Toth R, James J, Ganley IG (2013) Loss of iron triggers PINK1/Parkin-independent mitophagy. EMBO Rep14: 1127–1135

Amara F, Berbenni M, Fragni M, Leoni G, Viggiani S, Ippolito VM, Larocca M, Rossano R, Alberghina L, Riccio P, Colangelo AM (2015) Neuroprotection by Cocktails of Dietary Antioxidants under Conditions of Nerve Growth Factor Deprivation. Oxidative medicine and cellular longevity 2015: 217258

Arena G, Gelmetti V, Torosantucci L, Vignone D, Lamorte G, De Rosa P, Cilia E, Jonas EA, Valente EM (2013) PINK1 protects against cell death induced by mitochondrial depolarization, by phosphorylating Bcl-xL and impairing its pro-apoptotic cleavage. Cell death and differentiation 20: 920–930

Ariga H, Takahashi-Niki K, Kato I, Maita H, Niki T, Iguchi-Ariga SM (2013) Neuroprotective function of DJ-1 in Parkinson’s disease. Oxidative medicine and cellular longevity 2013: 683920

Barabasi AL (2012) Network science: Luck or reason. Nature 489: 507–508

Bhatia TN, Pant DB, Eckhoff EA, Gongaware RN, Do T, Hutchison DF, Gleixner AM, Leak RK (2019) Astrocytes Do Not Forfeit Their Neuroprotective Roles After Surviving Intense Oxidative Stress. Frontiers in molecular neuroscience 12: 87

Bjelakovic G, Gluud C (2007) Surviving antioxidant supplements. Journal of the National Cancer Institute 99: 742–743

Bjorklund A, Cenci MA. (2010) Recent advances in Parkinson’s disease: basic research. Progress in brain research 183. Elsevier, Amsterdam etc., pp. 1 online resource (x, 320 p.).

Boogerd FC, Bruggeman FJ, Richardson RC, Stephan A, Westerhoff HV (2005) Emergence and its place in nature: A case study of biochemical networks (vol 145, pg 131, 2005). Synthese 145: 501–501

Boogerd FC, Bruggeman, F., Hofmeyr, J.H.S., Westerhoff, H.V (2007) Systems Biology Philosophical Foundations. 342 p

Brady NR, Elmore SP, van Beek JJ, Krab K, Courtoy PJ, Hue L, Westerhoff HV (2004) Coordinated behavior of mitochondria in both space and time: a reactive oxygen species-activated wave of mitochondrial depolarization. Biophysical journal 87: 2022–2034

Chaudhari P, Ye Z, Jang YY (2014) Roles of reactive oxygen species in the fate of stem cells. Antioxidants & redox signaling 20: 1881–1890

Cloutier M, Middleton R, Wellstead P (2012) Feedback motif for the pathogenesis of Parkinson’s disease. IET systems biology 6: 86–93

Cloutier M, Wellstead P (2012) Dynamic modelling of protein and oxidative metabolisms simulates the pathogenesis of Parkinson’s disease. IET systems biology 6: 65–72

Cui T, Fan C, Gu L, Gao H, Liu Q, Zhang T, Qi Z, Zhao C, Zhao H, Cai Q, Yang H (2011) Silencing of PINK1 induces mitophagy via mitochondrial permeability transition in dopaminergic MN9D cells. Brain research 1394: 1–13

Deferme L, Briede JJ, Claessen SM, Jennen DG, Cavill R, Kleinjans JC (2013) Time series analysis of oxidative stress response patterns in HepG2: a toxicogenomics approach. Toxicology 306: 24–34

del Sol A, Balling R, Hood L, Galas D (2010) Diseases as network perturbations. Curr Opin Biotechnol 21: 566–571

DeNicola GM, Karreth FA, Humpton TJ, Gopinathan A, Wei C, Frese K, Mangal D, Yu KH, Yeo CJ, Calhoun ES, Scrimieri F, Winter JM, Hruban RH, Iacobuzio-Donahue C, Kern SE, Blair IA, Tuveson DA (2011) Oncogene-induced Nrf2 transcription promotes ROS detoxification and tumorigenesis. Nature 475: 106–109

Dhandapani KM, Wade FM, Wakade C, Mahesh VB, Brann DW (2005) Neuroprotection by stem cell factor in rat cortical neurons involves AKT and NFkappaB. Journal of neurochemistry 95: 9–19

Edgar R, Domrachev M, Lash AE (2002) Gene Expression Omnibus: NCBI gene expression and hybridization array data repository. Nucleic Acids Res 30: 207–210

Fedorowicz MA, de Vries-Schneider RL, Rub C, Becker D, Huang Y, Zhou C, Alessi Wolken DM, Voos W, Liu Y, Przedborski S (2014) Cytosolic cleaved PINK1 represses Parkin translocation to mitochondria and mitophagy. EMBO Rep 15: 86–93

Finnerty NJ, O’Riordan SL, Lowry JP, Cloutier M, Wellstead P (2013) Continuous real-time in vivo measurement of cerebral nitric oxide supports theoretical predictions of an irreversible switching in cerebral ROS after sufficient exposure to external toxins. Journal of Parkinson’s disease 3: 351–362

Fujita KA, Ostaszewski M, Matsuoka Y, Ghosh S, Glaab E, Trefois C, Crespo I, Perumal TM, Jurkowski W, Antony PM, Diederich N, Buttini M, Kodama A, Satagopam VP, Eifes S, Del Sol A, Schneider R, Kitano H, Balling R (2014) Integrating pathways of Parkinson’s disease in a molecular interaction map. Molecular neurobiology 49: 88–102

Guo L, Lobenhofer EK, Wang C, Shippy R, Harris SC, Zhang L, Mei N, Chen T, Herman D, Goodsaid FM, Hurban P, Phillips KL, Xu J, Deng X, Sun YA, Tong W, Dragan YP, Shi L (2006) Rat toxicogenomic study reveals analytical consistency across microarray platforms. Nature biotechnology 24: 1162–1169

Hall CJ, Boyle RH, Astin JW, Flores MV, Oehlers SH, Sanderson LE, Ellett F, Lieschke GJ, Crosier KE, Crosier PS (2013) Immunoresponsive gene 1 augments bactericidal activity of macrophage-lineage cells by regulating beta-oxidation-dependent mitochondrial ROS production. Cell Metab 18: 265–278

He F, Murabito E, Westerhoff HV (2016) Synthetic biology and regulatory networks: where metabolic systems biology meets control engineering. Journal of the Royal Society, Interface / the Royal Society 13

Ivatt RM, Whitworth AJ (2014) The many faces of mitophagy. EMBO Rep 15: 5–6

Janssen-Heininger YM, Poynter ME, Baeuerle PA (2000) Recent advances towards understanding redox mechanisms in the activation of nuclear factor kappaB. Free radical biology & medicine 28: 1317–1327

Jennen DG, Magkoufopoulou C, Ketelslegers HB, van Herwijnen MH, Kleinjans JC, van Delft JH (2010) Comparison of HepG2 and HepaRG by whole-genome gene expression analysis for the purpose of chemical hazard identification. Toxicological sciences: an official journal of the Society of Toxicology 115: 66–79

Joselin AP, Hewitt SJ, Callaghan SM, Kim RH, Chung YH, Mak TW, Shen J, Slack RS, Park DS (2012) ROS-dependent regulation of Parkin and DJ-1 localization during oxidative stress in neurons. Human molecular genetics 21: 4888–4903

Juntilla MM, Patil VD, Calamito M, Joshi RP, Birnbaum MJ, Koretzky GA (2010) AKT1 and AKT2 maintain hematopoietic stem cell function by regulating reactive oxygen species. Blood 115: 4030–4038

Kanda Y, Hinata T, Kang SW, Watanabe Y (2011) Reactive oxygen species mediate adipocyte differentiation in mesenchymal stem cells. Life Sci 89: 250–258

Kane LA, Lazarou M, Fogel AI, Li Y, Yamano K, Sarraf SA, Banerjee S, Youle RJ (2014) PINK1 phosphorylates ubiquitin to activate Parkin E3 ubiquitin ligase activity. The Journal of cell biology 205: 143–153

Kell DB (2010) Towards a unifying, systems biology understanding of large-scale cellular death and destruction caused by poorly liganded iron: Parkinson’s, Huntington’s, Alzheimer’s, prions, bactericides, chemical toxicology and others as examples. Archives of toxicology 84: 825–889

Kinder M, Wei C, Shelat SG, Kundu M, Zhao L, Blair IA, Pure E (2010) Hematopoietic stem cell function requires 12/15-lipoxygenase-dependent fatty acid metabolism. Blood 115: 5012–5022

Kitano H, Funahashi A, Matsuoka Y, Oda K (2005) Using process diagrams for the graphical representation of biological networks. Nature biotechnology 23: 961–966

Kodama T, Takehara T, Hikita H, Shimizu S, Shigekawa M, Li W, Miyagi T, Hosui A, Tatsumi T, Ishida H, Kanto T, Hiramatsu N, Yin XM, Hayashi N (2011) BH3-only activator proteins Bid and Bim are dispensable for Bak/Bax-dependent thrombocyte apoptosis induced by Bcl-xL deficiency: molecular requisites for the mitochondrial pathway to apoptosis in platelets. The Journal of biological chemistry 286: 13905–13913

Kolodkin AN, Boogerd FC, Bruggeman FJ, Westerhoff HV (2011) Modeling Approaches in Systems Biology, Including Silicon Cell Models. In Systems Biology and Livestock Science, Pas tMFW, Woelders H, Bannink A (eds), 2. Oxford, UK: Wiley-Blackwell

Kolodkin AN, Bruggeman FJ, Plant N, Mone MJ, Bakker BM, Campbell MJ, van Leeuwen JP, Carlberg C, Snoep JL, Westerhoff HV (2010) Design principles of nuclear receptor signaling: how complex networking improves signal transduction. Mol Syst Biol 6: 446

Koyano F, Okatsu K, Kosako H, Tamura Y, Go E, Kimura M, Kimura Y, Tsuchiya H, Yoshihara H, Hirokawa T, Endo T, Fon EA, Trempe JF, Saeki Y, Tanaka K, Matsuda N (2014) Ubiquitin is phosphorylated by PINK1 to activate parkin. Nature

Kruger R, Hilker R, Winkler C, Lorrain M, Hahne M, Redecker C, Lingor P, Jost WH (2015) Advanced stages of PD: interventional therapies and related patient-centered care. Journal of neural transmission

Le Belle JE, Orozco NM, Paucar AA, Saxe JP, Mottahedeh J, Pyle AD, Wu H, Kornblum HI (2011) Proliferative neural stem cells have high endogenous ROS levels that regulate self-renewal and neurogenesis in a PI3K/Akt-dependant manner. Cell stem cell 8: 59–71

Lee SJ, Lim HS, Masliah E, Lee HJ (2011) Protein aggregate spreading in neurodegenerative diseases: Problems and perspectives. Neuroscience research 70: 339–348

Lewandowski D, Barroca V, Duconge F, Bayer J, Van Nhieu JT, Pestourie C, Fouchet P, Tavitian B, Romeo PH (2010) In vivo cellular imaging pinpoints the role of reactive oxygen species in the early steps of adult hematopoietic reconstitution. Blood 115: 443–452

Lewis KN, Wason E, Edrey YH, Kristan DM, Nevo E, Buffenstein R (2015) Regulation of Nrf2 signaling and longevity in naturally long-lived rodents. Proceedings of the National Academy of Sciences of the United States of America 112: 3722–3727

Li X, Fang P, Mai J, Choi ET, Wang H, Yang XF (2013a) Targeting mitochondrial reactive oxygen species as novel therapy for inflammatory diseases and cancers. Journal of hematology & oncology 6: 19

Li Y, Cui J, Zhang CH, Yang DJ, Chen JH, Zan WH, Li B, Li Z, He YL (2013b) High-expression of DJ-1 and loss of PTEN associated with tumor metastasis and correlated with poor prognosis of gastric carcinoma. International journal of medical sciences 10: 1689–1697

Limoli CL, Rola R, Giedzinski E, Mantha S, Huang TT, Fike JR (2004) Cell-density-dependent regulation of neural precursor cell function. Proceedings of the National Academy of Sciences of the United States of America 101: 16052–16057

Lingappan K (2018) NF-kappaB in Oxidative Stress. Current opinion in toxicology 7: 81–86

Loscalzo J, Barabasi AL (2011) Systems biology and the future of medicine. Wiley Interdiscip Rev Syst Biol Med 3: 619–627

Martorana F, Gaglio D, Bianco MR, Aprea F, Virtuoso A, Bonanomi M, Alberghina L, Papa M, Colangelo AM (2018) Differentiation by nerve growth factor (NGF) involves mechanisms of crosstalk between energy homeostasis and mitochondrial remodeling. Cell death & disease 9: 391

Matsuda S, Nakanishi A, Minami A, Wada Y, Kitagishi Y (2015) Functions and characteristics of PINK1 and Parkin in cancer. Front Biosci (Landmark Ed) 20: 491–501

Morais VA, Haddad D, Craessaerts K, De Bock PJ, Swerts J, Vilain S, Aerts L, Overbergh L, Grunewald A, Seibler P, Klein C, Gevaert K, Verstreken P, De Strooper B (2014) PINK1 loss-of-function mutations affect mitochondrial complex I activity via NdufA10 ubiquinone uncoupling. Science 344: 203–207

Muller-Rischart AK, Pilsl A, Beaudette P, Patra M, Hadian K, Funke M, Peis R, Deinlein A, Schweimer C, Kuhn PH, Lichtenthaler SF, Motori E, Hrelia S, Wurst W, Trumbach D, Langer T, Krappmann D, Dittmar G, Tatzelt J, Winklhofer KF (2013) The E3 Ligase Parkin Maintains Mitochondrial Integrity by Increasing Linear Ubiquitination of NEMO. Mol Cell 49: 908–921

Nalls MA, Pankratz N, Lill CM, Do CB, Hernandez DG, Saad M, DeStefano AL, Kara E, Bras J, Sharma M, Schulte C, Keller MF, Arepalli S, Letson C, Edsall C, Stefansson H, Liu X, Pliner H, Lee JH, Cheng R et al (2014) Large-scale meta-analysis of genome-wide association data identifies six new risk loci for Parkinson’s disease. Nat Genet

Oyewole AO, Birch-Machin MA (2015) Mitochondrial-targeted antioxidants. FASEB J

Palacios N, Gao X, McCullough ML, Schwarzschild MA, Shah R, Gapstur S, Ascherio A (2012) Caffeine and risk of Parkinson’s disease in a large cohort of men and women. Movement disorders: official journal of the Movement Disorder Society 27: 1276–1282

Pfeiffer R, Wszolek ZK, Ebadi MS. (2012) Parkinson’s disease. CRC Press, Boca Raton, FL, pp. 1 online resource (xxix, 1278 p., 1224 p. of plates).

Polyzos AA, Lee DY, Datta R, Hauser M, Budworth H, Holt A, Mihalik S, Goldschmidt P, Frankel K, Trego K, Bennett MJ, Vockley J, Xu K, Gratton E, McMurray CT (2019) Metabolic Reprogramming in Astrocytes Distinguishes Region-Specific Neuronal Susceptibility in Huntington Mice. Cell Metab

Postuma RB, Lang AE, Munhoz RP, Charland K, Pelletier A, Moscovich M, Filla L, Zanatta D, Rios Romenets S, Altman R, Chuang R, Shah B (2012) Caffeine for treatment of Parkinson disease: a randomized controlled trial. Neurology 79: 651–658

Rakovic A, Shurkewitsch K, Seibler P, Grunewald A, Zanon A, Hagenah J, Krainc D, Klein C (2013) Phosphatase and tensin homolog (PTEN)-induced putative kinase 1 (PINK1)-dependent ubiquitination of endogenous Parkin attenuates mitophagy: study in human primary fibroblasts and induced pluripotent stem cell-derived neurons. The Journal of biological chemistry 288: 2223–2237

Santiago JA, Potashkin JA (2013) Shared dysregulated pathways lead to Parkinson’s disease and diabetes. Trends in molecular medicine 19: 176–186

Sha D, Chin LS, Li L (2010) Phosphorylation of parkin by Parkinson disease-linked kinase PINK1 activates parkin E3 ligase function and NF-kappaB signaling. Human molecular genetics 19: 352–363

Shore DE, Carr CE, Ruvkun G (2012) Induction of cytoprotective pathways is central to the extension of lifespan conferred by multiple longevity pathways. PLoS genetics 8: e1002792.

Soleimanpour SA, Gupta A, Bakay M, Ferrari AM, Groff DN, Fadista J, Spruce LA, Kushner JA, Groop L, Seeholzer SH, Kaufman BA, Hakonarson H, Stoffers DA (2014) The diabetes susceptibility gene clec16a regulates mitophagy. Cell 157: 1577–1590

Tamatani M, Mitsuda N, Matsuzaki H, Okado H, Miyake S, Vitek MP, Yamaguchi A, Tohyama M (2000) A pathway of neuronal apoptosis induced by hypoxia/reoxygenation: roles of nuclear factor-kappaB and Bcl-2. Journal of neurochemistry 75: 683–693

Tran HT, Chung CH, Iba M, Zhang B, Trojanowski JQ, Luk KC, Lee VM (2014) alpha-Synuclein Immunotherapy Blocks Uptake and Templated Propagation of Misfolded alpha-Synuclein and Neurodegeneration. Cell reports

van der Mark M, Nijssen PC, Vlaanderen J, Huss A, Mulleners WM, Sas AM, van Laar T, Kromhout H, Vermeulen R (2014) A case-control study of the protective effect of alcohol, coffee, and cigarette consumption on Parkinson disease risk: time-since-cessation modifies the effect of tobacco smoking. PLoS one 9: e95297.

Varcin M, Bentea E, Michotte Y, Sarre S (2012) Oxidative stress in genetic mouse models of Parkinson’s disease. Oxidative medicine and cellular longevity 2012: 624925

Walsh DM, Selkoe DJ (2016) A critical appraisal of the pathogenic protein spread hypothesis of neurodegeneration. Nat Rev Neurosci 17: 251–260

Wang X, Winter D, Ashrafi G, Schlehe J, Wong YL, Selkoe D, Rice S, Steen J, Lavoie MJ, Schwarz TL (2011) PINK1 and Parkin Target Miro for Phosphorylation and Degradation to Arrest Mitochondrial Motility. Cell 147: 893–906

Wang Y, Liu W, He X, Zhou F (2013) Parkinson’s disease-associated DJ-1 mutations increase abnormal phosphorylation of tau protein through Akt/GSK-3beta pathways. Journal of molecular neuroscience: MN 51: 911–918

Williamson TP, Johnson DA, Johnson JA (2012) Activation of the Nrf2-ARE pathway by siRNA knockdown of Keap1 reduces oxidative stress and provides partial protection from MPTP-mediated neurotoxicity. Neurotoxicology 33: 272–279

Xue M, Momiji H, Rabbani N, Barker G, Bretschneider T, Shmygol A, Rand DA, Thornalley PJ (2014) Frequency Modulated Translocational Oscillations of Nrf2 Mediate the Antioxidant Response Element Cytoprotective Transcriptional Response. Antioxidants & redox signaling

Yun J, Finkel T (2014) Mitohormesis. Cell Metab 19: 757–766

