## Supplementary material for "Design principles of ROS dynamic networks relevant to precision therapies for age-related diseases": Supplemenatary Materials

### Supplementary Materials

SM.H. The effect of α-synuclein on ATP and ROS concentrations in the detailed model . 63

#### SM.A. The simplest "no-design" module. Design A. Model A.

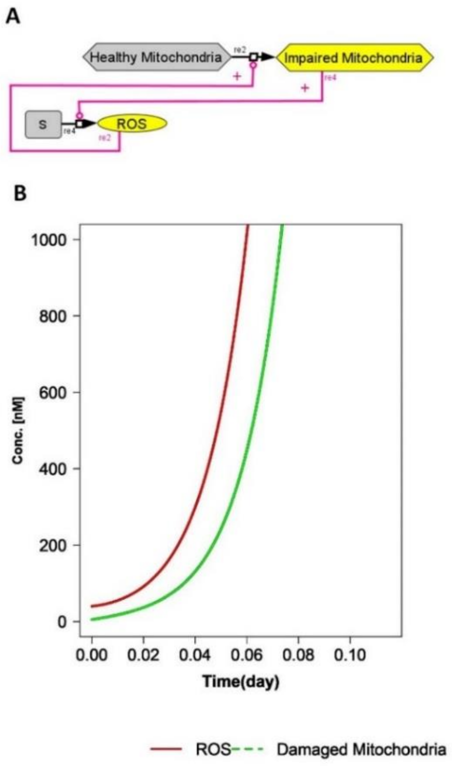

**Figure SM.A.1. The 'No design' Model A.** Network diagram showing the positive feedback loop formed by ROS damaging the mitochondria (re2) and damaged mitochondria producing ROS (re4). **B.** Simulated concentrations of ROS and damaged mitochondria (model A.cps) corresponding to the "no-design" case.

##### The following conclusions were drawn from Figure SM.A.1:

The system is unstable and is not able to reach a steady state.

##### Model building and simulations:

Model A.cps was used for generation of the figure. The model and the simulation results have been assembled into the model archive file called 'SM-A-no design'. The general idea used in the modelling is described below:
The reaction of ROS generation is catalyzed by impaired mitochondria at the rate described by the mass action equation:

Rate of ROSgeneration=ROSsynCoef*([DamagedMitochondria(t)]).

Thus, in the absence of damaged mitochondria ROS are not produced. Analogously, the mitochondrial aging is catalyzed by ROS at a rate described by the mass action equation: Rate of Mitochondrial Aging=k*([HealthyMitochondria(t)]*[ROS(t)]).

Thus, in the absence of ROS, mitochondria are not damaged. Simulations started at initial conditions where the concentration of ROS was equal to $10^{-5}$ (nM) and the concentration of damaged mitochondria was equal to $10^{-5}$ (arbitrary units). Time courses for ROS and Damaged Mitochondria were simulated.

No steady state is observed. The concentration of both ROS and Damaged Mitochondria varies exponentially with time ('exploded').

#### SM.B1. Model B1.

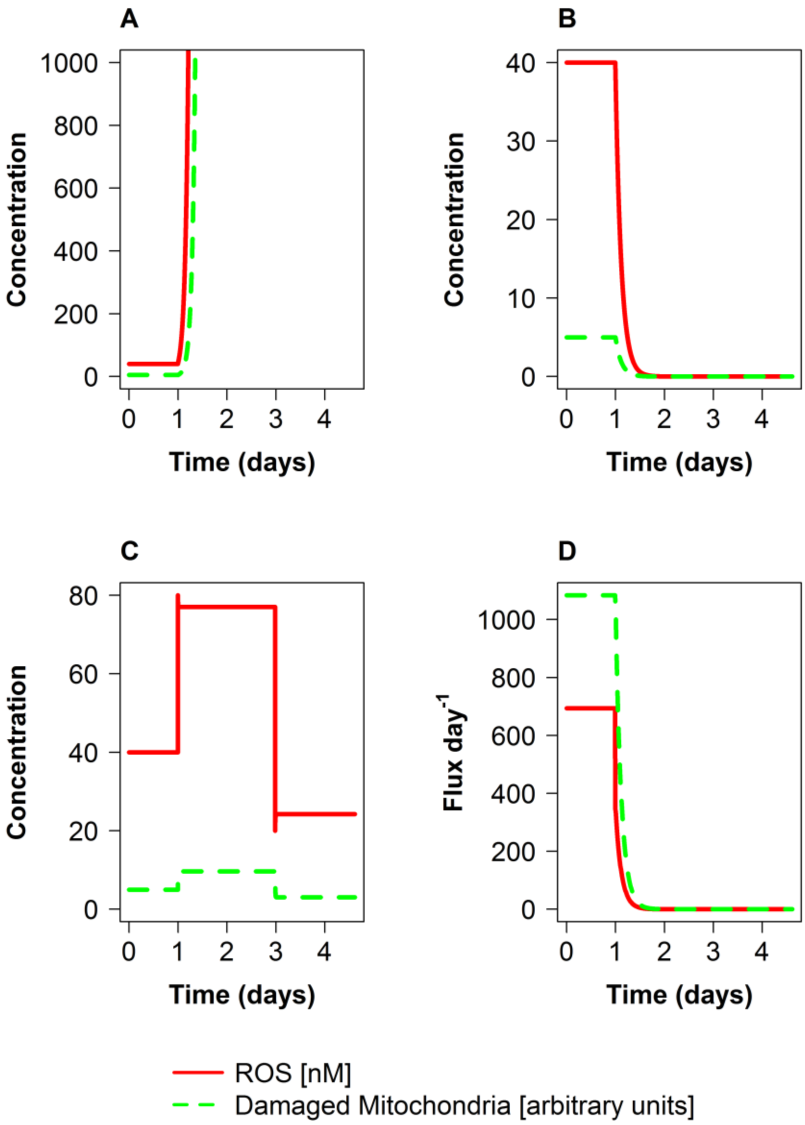

**Figure SM.B1.1. Steady state in the ROS core model. Structural and dynamic instability of the ROS core model (Model B1; design 1).**

Concentration of ROS (in nM) and damaged mitochondria (in arbitrary units) (A-C), fluxes of ROS production and generation of damaged mitochondria (D) for doubling ROS generation rate constant on day 1 (A), for halving the rate constant on day 1 followed by its restoration to its original value on day 3 (B, D), and for injection of ROS at day 1 followed by a wash out on day 3 (C).

**The following conclusions were drawn from Figure SM.B1.1:**

When simulations starts from a specific set of initial conditions, the initial state persists if a steady state is maintained. However, when the ROS generation rate constant is doubled on day 1, both the concentrations of ROS and of damaged mitochondria increase indefinitely and the system ‘explodes’: the state is structurally unstable. When the rate constant of ROS production is instead halved, the system relaxes irreversibly to the “quasi-perfect” state with the concentrations of ROS and damaged mitochondria equal to 0. The fluxes in the systems disappear accordingly. When on day 3 ROS generation is returned to its initial rate, the system stays at the same “quasi-perfect” zero-state. When at the initial steady state ROS are injected or eliminated, ROS concentration changes as expected, but does not relax back to its initial value, showing that the steady state is not dynamically stable in the sense of Lyapunov stability.

**Model building and simulations:**
Model B1 (B1.cps) used for generation of every figure, as well as simulation results have been assembled in the model archive file SM-B1.

The general idea used in the modelling is described below:

Generation of ROS is catalyzed by impaired mitochondria at a rate described by the mass action equation: Rate of ROSgeneration=ROSsynCoef*([DamagedMitochondria(t)]). Thus, in the absence of damaged mitochondria ROS are not produced.

Analogously, mitochondrial aging is catalyzed by ROS with the rate described by the mass action equation: Rate of Mitochondrial Aging=k*([HealthyMitochondria(t)]*[ROS(t)]). Thus, in the absence of ROS, mitochondria are not damaged.

Both rates of removal of ROS and removal of impaired mitochondria (mitophagy) were modelled as mass action reactions:
Rate of ROS removal = k*[ROS(t)]*[Antiox(t)]
Rate of Mitophagy = k*[Impaired mitochondria(t)]*[parkin(t)] *[p62(t)]

Antiox was included in the reaction as catalyst (i.e. without being consumed) because we considered not only antioxidant metabolites (e.g. various antioxidants which the cell would maintain at constant concentrations), but also enzymes (e.g. superoxide dismutase) catalyzing ROS removal. P62 and parkin are consumed for mitophagy as substrates. Antiox, p62 and parkin concentrations are taken constant, i.e. independent of time. Simulations started with initial conditions where the concentration of ROS was equal to 40 nM and the concentration of damaged mitochondria was equal to 5 nM. Time courses for ROS and Damaged Mitochondria were then simulated using Copasi.

#### SM.B2A. Design 2A. Models B2.A1-B2.A3

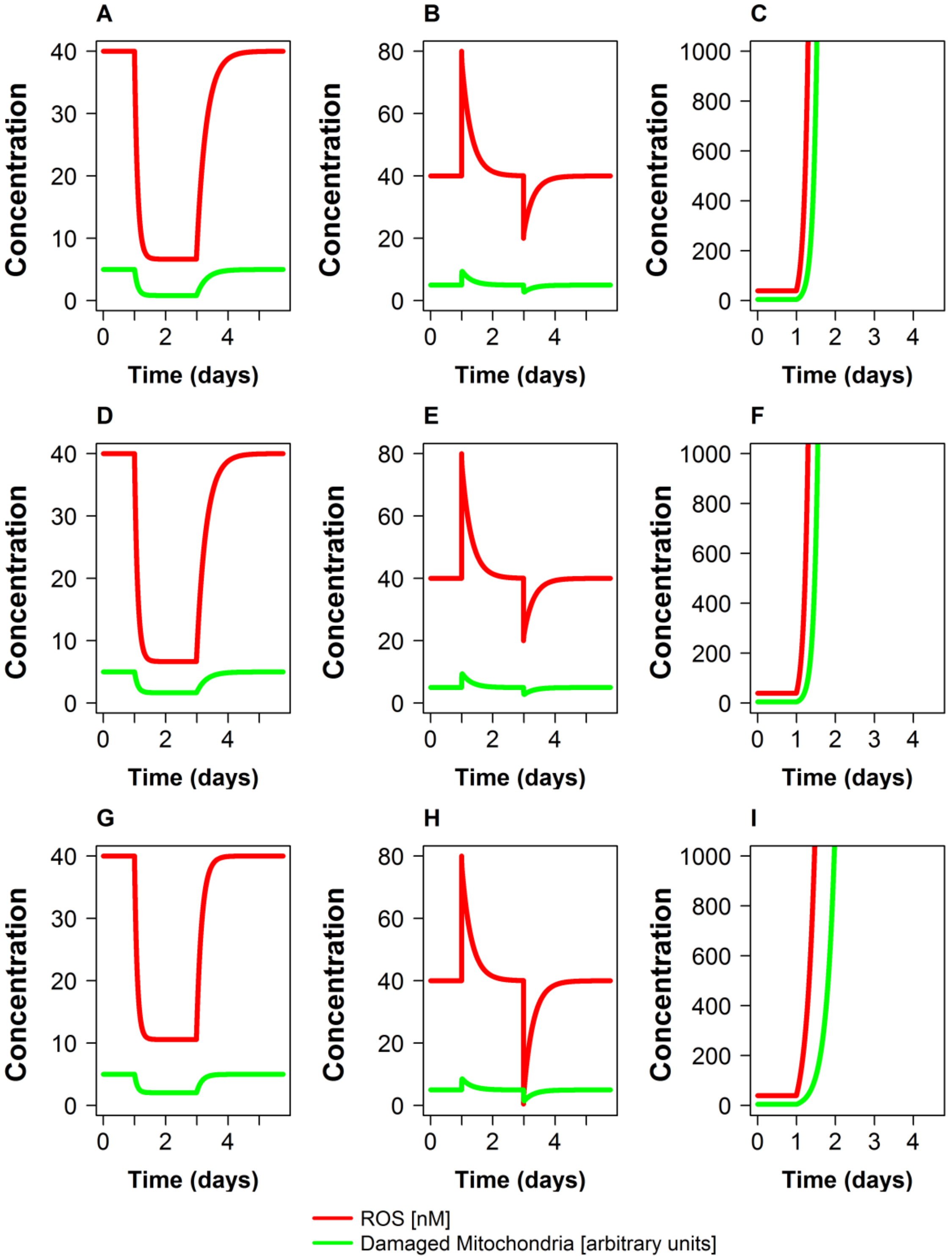

**Figure SM.B2A.1 (Models B2A.1-B2A.3). Dynamic stability obtained when basal ROS generation and/or basal mitoptosis (i.e. autophagy specific for impaired mitochondria) were added to Design 1 thereby obtaining Design 2A.** Concentrations of ROS (in nM) and damaged mitochondria (in arbitrary units) for three subtypes of Design 2A. Simulations used a model with basal ROS generation only (A-C; top row: model B2A.1-BasalROS.cps), a model with basal mitoptosis only (D-F; middle row; model B2A.2-BasalMitochondrialAging.cps) or a model with both basal ROS generation and basal mitoptosis (G-I; bottom row; model B2A.3-BasalAll.cps). In A, D, and G, ROS generation rate constant was first decreased 2 fold (day 1) and then returned back (day 3). In B, E, and H, ROS concentration was instantaneously increased two fold (day 1) and then decreased two fold (day 3), whereas in C, F, and I, ROS generation rate constant was increased 2 fold on day 1.

**The following conclusions were drawn from Figure SM.B2A.1:**

**A**) Concentrations of both ROS and damaged mitochondria follows the perturbation of ROS synthesis. Concentrations of both species decrease 6-fold to a lower steady value and then return to their initial levels. **B)** Concentrations of both ROS and damaged mitochondria transiently respond to perturbations, but both concentration changes are transient: ROS and damaged mitochondrial concentrations ultimately return to their initial levels. **C)** The system explodes. **D)** The concentration of ROS follows the perturbation, first decreasing 6-fold and then returning to initial level. The concentration of damaged mitochondria follows the perturbation, first decreasing 3-fold and then returning to initial level. **E)** Concentrations of both ROS and damaged mitochondria transiently respond to perturbations, but both concentration changes are transient: ROS and damaged mitochondrial concentrations ultimately return to their initial levels. **F**) The system explodes. **G**) ROS concentration follows the perturbation, first decreasing around 4-fold and then returning to initial level. The concentration of damaged mitochondria follows the perturbation, first decreasing around 2.5-fold and then returning to initial level. **H**) Concentrations of both ROS and damaged mitochondria transiently respond to perturbations, but both concentration changes are transient: ROS and damaged mitochondrial concentrations ultimately return to their initial levels. I: The system explodes.

We conclude that for limited extents of perturbation, the system is both structurally and dynamically stable, whilst for substantial upward perturbation of ROS production, the system remains unstable.

**Model building and simulations:**

A compendium of models B2A.1-B2A.3 (B2A.1-BasalROS.cps, B2A.2-BasalMitochondrialAging.cps and B2A.3-BasalAll.cps) used for generation of every figure, as well as simulation results have been assembled in the model archive file SM-B2A.

The general idea used in the modelling is described below:

Both rates of removal of ROS and removal of impaired mitochondria (mitophagy) are modelled as mass action reactions:
Rate of ROS removal = k*[ROS(t)]*[Antiox(t)]
Rate of Mitophagy = k*[Impaired mitochondria(t)]*[parkin(t)] *[p62(t)]

Antiox was included in the reaction as catalyst (i.e. without being consumed) because we considered not only antioxidant metabolites (e.g. various antioxidants which the cell would maintain at constant concentrations), but also enzymes (e.g. superoxide dismutase) catalyzing ROS removal. P62 and parkin are consumed for mitophagy as substrates. Antiox, p62 and parkin concentrations are taken constant, i.e. independent of time. Simulations started with initial conditions where the concentration of ROS was equal to 40 nM and the concentration of damaged mitochondria was equal to 5 nM. Time courses for ROS and Damaged Mitochondria were then simulated using Copasi.

*Model with basal ROS generation (*B2A.1-BasalROS.cps*):*

The rate of mitochondrial aging was described in the same way as in model B1:
Rate of Mitochondrial Aging=k*([HealthyMitochondria(t)]*[ROS(t)])

However, the rate of ROS generation was described by the following reaction:
Rate of ROSgeneration=ROSsynCoef*([DamagedMitochondria(t)]+kbasalROS)
Thus, even in the absence of damaged mitochondria ROS are still produced to a certain extent.

In the model B2.2-BasalROS.cps, values of ROSsynCoef and kbasalROS were calculated analytically so that initial steady state values of all variables in model B2.2-BasalROS.cps were equal to steady state values of these variables in model B1.cps (design 1).

*Model where mitochondria may get damaged also in the absence of ROS (*B2A.2-BasalMitochondrialAging.cps):

The rate of ROS generation was described in the same way as in model B1:
Rate of ROSgeneration=ROSsynCoef*([DamagedMitochondria(t)])

However, the rate of mitochondrial aging was described by the following reaction:
Rate of Mitochondrial Aging =
MitochAgingCoef*([HealthyMitochondria(t)]*[ROS(t)]+kbasalMitochAging)
Thus, even in the absence of ROS, mitochondrial damage takes place to a certain extent.

Values of MitochAgingCoef and kbasalMitochAging were calculated analytically so that initial steady state values of all variables in model B2A.2-BasalMitochondrialAging.cps were equal to steady state values of these variables in model B1.cps (design 1).

*Model where mitochondria may get damaged in the absence of ROS and ROS may be produced in the absence of damaged mitochondria (Model* B2A.3-BasalAll.cps*):*

The rate of mitochondrial aging was described by the following reaction:
Rate of Mitochondrial Aging=
MitochAgingCoef * ([HealthyMitochondria(t)]*[ROS(t)]+kbasalMitochAging)

The rate of ROS generation was described by the following reaction:
Rate of ROSgeneration=ROSsynCoeficient* ([DamagedMitochondria(t)]+kbasalROS)
Thus, even in the absence of ROS, mitochondrial damage takes place to a certain extent. As well, ROS are produced in the absence of damaged mitochondria.

Values of ROSsynCoef, kbasalROS, MitochAgingCoef and kbasalMitochAging were calculated analytically so that initial steady state values of all variables in model B2A.3-BasalAll.cps were equal to steady state values of these variables in model B1.cps (design 1).

#### SM.B2B. Design 2B. Model B2B.

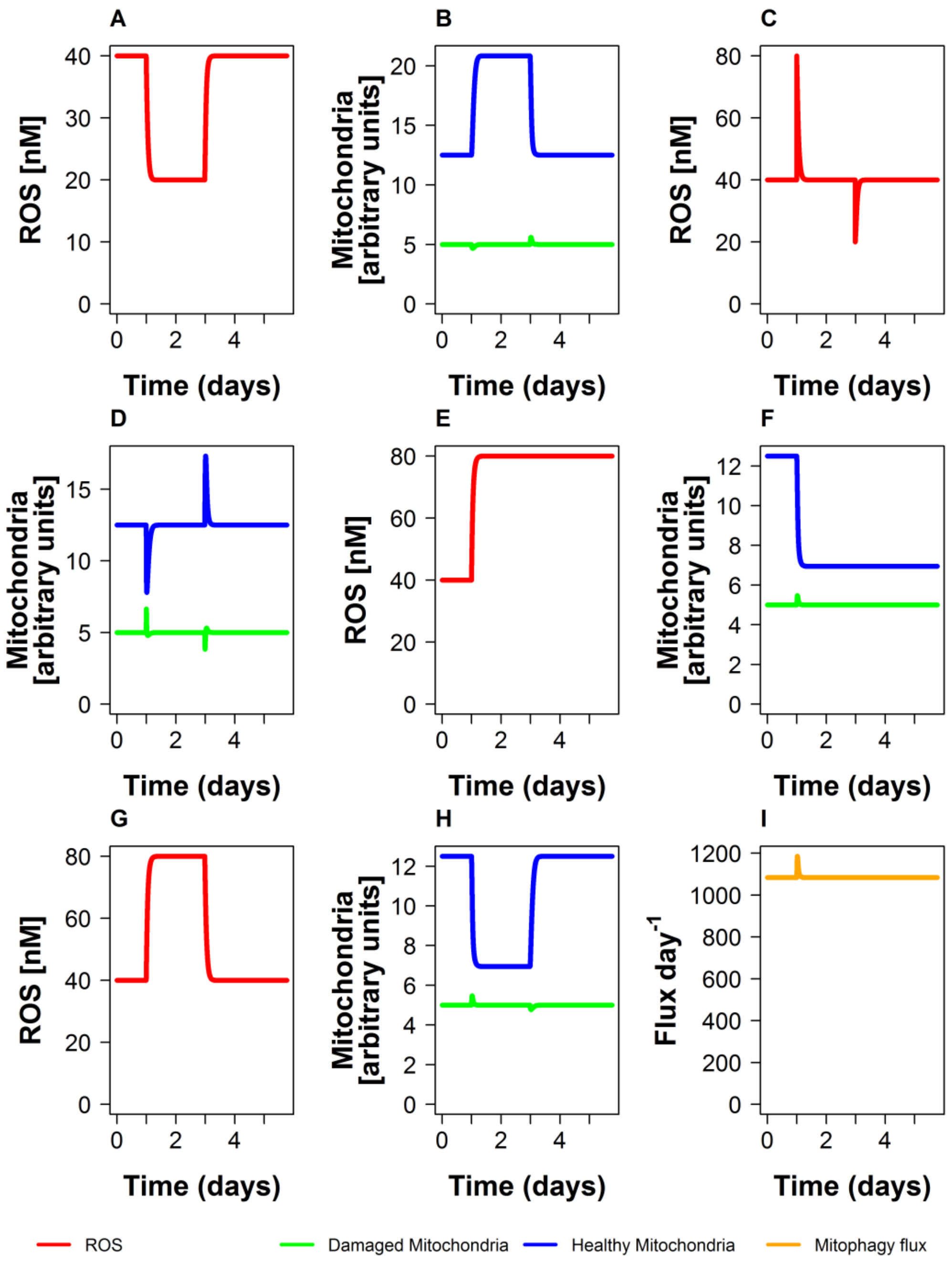

**Figure SM.B2B.1. Design 2B (Model B2B.cps). Dynamic and structural stability obtained when the reaction of mitochondrial synthesis was added to Design 1 thereby obtaining Design 2B.** Concentrations of ROS (in nM) and damaged mitochondria (in arbitrary units) for design 2B. Simulations used a model 2B.cps. In A and B, ROS generation rate constant was first decreased 2 fold (day 1) and then returned back (day 3). In C and D, ROS concentration was instantaneously increased two fold (day 1) and then decreased two fold (day 3). In E, F and I the ROS generation rate constant was increased 2 fold on day 1. In G and H, ROS generation rate constant was increased 2 fold on day 1 and then decreased to the initial level on day 3.

**The following conclusions were drawn from Figure SM.B2B.1:**

A: The concentration of ROS follows the perturbation of ROS synthesis. First, ROS concentration decrease 2 fold to a lower steady value and then return back to its initial level.

B: The concentration of healthy mitochondria follows the perturbation of ROS synthesis; first the concentration of healthy mitochondria increases to a higher steady value and then return back to its initial level. The concentration of damaged mitochondria respond to perturbations, but the concentration changes are small and transient, damaged mitochondrial concentration is ultimately returned to its initial level.

C: The concentration of ROS transiently respond to perturbations, but concentration changes are transient, ROS concentration is ultimately returned to its initial level.

D: Concentrations of both healthy and damaged mitochondria transiently respond to perturbations, but both concentration changes are transient, healthy and damaged mitochondrial concentrations are ultimately returned to their initial levels.

E: ROS concentration follows the perturbation of ROS synthesis and increases 2 fold to a new steady value.

F: The concentration of healthy mitochondria follows the perturbation of ROS synthesis and decrease to a lower steady value. The concentration of damaged mitochondria responds to perturbations, but concentration changes are small and transient, damaged mitochondrial concentration is ultimately returned to its initial level.

G: ROS concentration follows the perturbation of ROS synthesis. First, ROS concentration increases 2 fold to a higher steady value and then returns back to its initial level.

H: The concentration of healthy mitochondria follows the perturbation of ROS synthesis; first, the concentration of healthy mitochondria decreases to a lower steady value and then returns back to its initial level. The concentration of damaged mitochondria responds to perturbations, but the concentration changes are small and transient, damaged mitochondrial concentration is ultimately returned to its initial level.

I. After the fluctuation at the moment of perturbation, the flux of mitophagy returns to the initial level.

We conclude that the system is both structurally and dynamically stable, for both downward and upward perturbation of ROS production.

**Models building and simulations:**

Model B2B and all simulations for generation of every figure have been assembled in the model archive file (SM.B2B).

The general idea used in the modelling is described below:

(re1) Mitochondria are synthesised with the constant rate. The rate constant is set in such a way that the concentration of healthy mitochondria is equal to the concentration of healthy mitochondria in models B1 and B2A.

(re2) The reaction of mitochondrial aging is catalyzed by ROS with the rate described by mass action kinetics: Rate of Mitochondrial Aging=k*([HealthyMitochondria(t)]*[ROS(t)]). Thus, in the absence of ROS, mitochondria were not damaged.

(re3) The rate of removal of impaired mitochondria (mitophagy) is modelled with irreversible mass action kinetics:
Rate of Mitophagy = k*[Impaired mitochondria(t)]*[parkin(t)] *[p62(t)]

(re4) The reaction of ROS generation is catalyzed by impaired mitochondria with the rate described by mass action kinetics:

Rate of ROSgeneration=ROSsynCoef*([DamagedMitochondria(t)]). Thus, in the absence of damaged mitochondria ROS are not produced.

(re5) ROS removal is modelled as irreversible mass action reaction:
Rate of ROS removal = k*[ROS(t)]*[Antiox(t)]

Antiox was included in the reaction as catalyst (i.e. without being consumed) because we considered not only antioxidant metabolites (e.g. various antioxidants which the cell would maintain at constant concentrations), but also enzymes (e.g. superoxide dismutase) catalyzing ROS removal. P62 and parkin are consumed for mitophagy as substrates. Antiox, p62 and parkin concentrations are taken constant, i.e. independent of time. Simulations started with initial conditions where the concentration of ROS was equal to 40 nM and the concentration of damaged mitochondria was equal to 5 nM. Time courses for ROS and Damaged Mitochondria were then simulated using Copasi.

#### SM.B3. Design 3. Model B3.

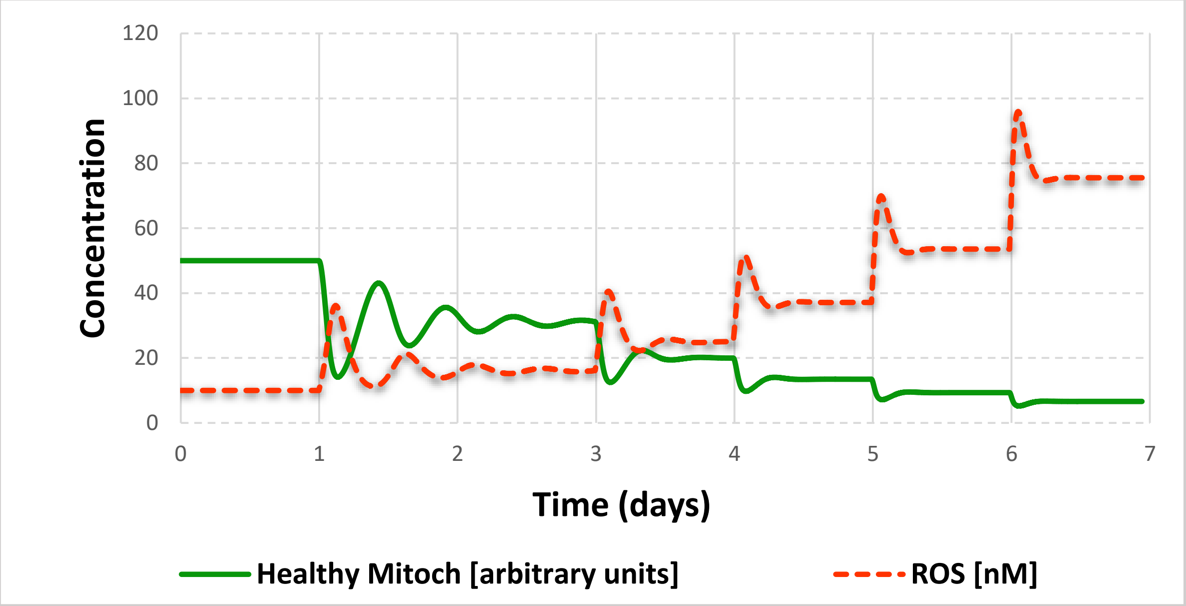

**Figure SM.B3.1. Design 3 (Model 3.cps). Homeostasis obtained when the negative feedback loop was added to design 2B thereby obtaining design 3.** Concentrations of ROS (in nM) and healthy mitochondria (in arbitrary units) for design 3. Simulations used a model 3.cps. ROS synthesis was doubled stepwise: day 1-ROS synthesis was doubled comparing to initial steady state (2 fold increase from initial value); day 3 – ROS synthesis was doubled comparing to day 1 (4 fold increase from initial value); day 4 – ROS synthesis was doubled comparing to day 3 (8 fold up from initial value); Day 5 – ROS synthesis was doubled comparing to day 4 (16 fold up from initial value); Day 6 – ROS synthesis was doubled comparing to day 5 (32 fold up from initial value).

**The following conclusions were drawn from Figure SMB3.1:**

In the response to each perturbation, concentrations of ROS and healthy mitochondria transiently reaches new steady state. Upon every doubling of ROS synthesis, ROS concentration is first increased twice, but then decreases back to the value a bit higher than the initial one. Upon every perturbation, the concentration of healthy mitochondria first decreases 2 fold, but then increases back to the value a bit lower than the steady state value before perturbation. We conclude that the system actively counteracts the increase of ROS.

**A**

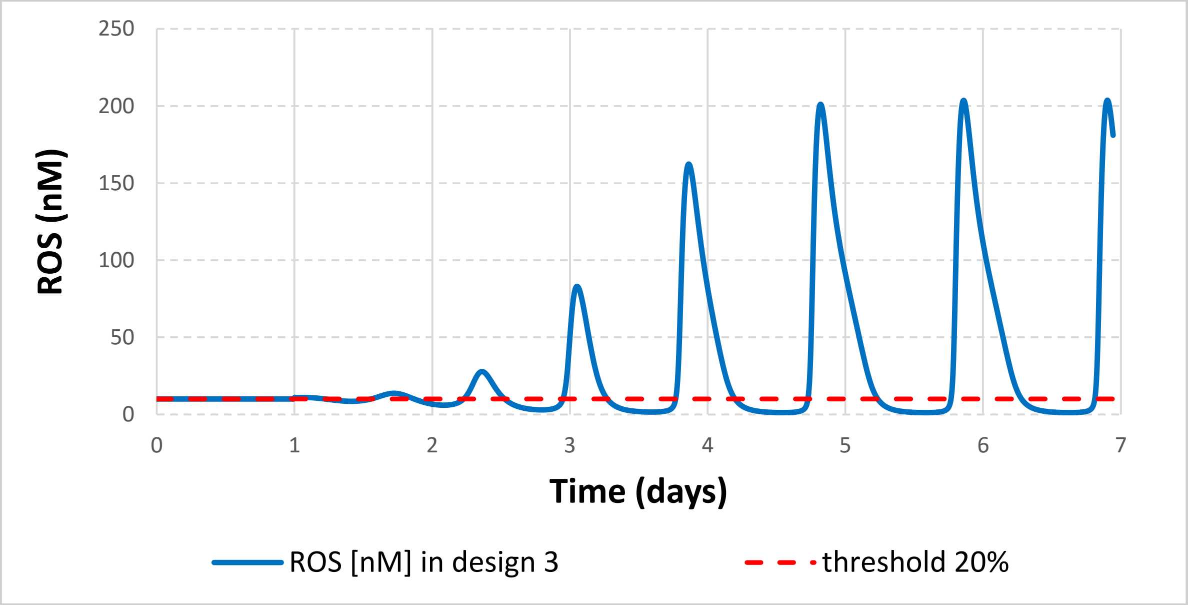

**B**

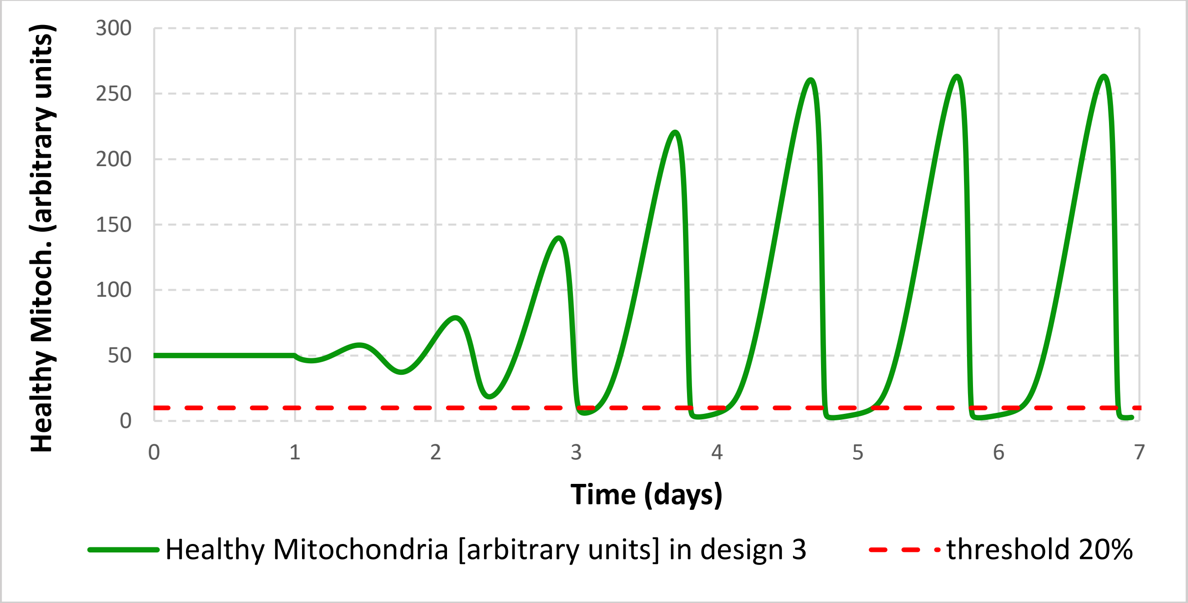

**Figure SM.B3.2. Design 3 (Model 3.cps). Compromised dynamic stability in homeostatic design 3.** Concentrations of (A) ROS (in nM) and (B) of healthy mitochondria (in arbitrary units) for design 3. Simulations used a model B3.cps. The initial ROS concentration was perturbed (transient increase from 10 nM to 11 nM) at day 1.

**The following conclusions were drawn from Figure SM.B3.2:**

A: The increase of ROS concentration from 10 nM t 11 nM at day 1 triggers an oscillatory behaviour of the ROS concentration.

B: The increase of ROS concentration from 10 nM to 11 nM at day 1 triggers an oscillatory behaviour of the ROS concentration. During oscillations, the concentration of healthy mitochondria sweeps below a hypothetical viability that corresponds to the threshold line (marked as a dotted red line) dissecting 20% (10 a.u) of the initial concentration of healthy mitochondria (50 a.u).

We conclude that the system is dynamically unstable against the perturbation of ROS concentration.

**Model building and simulations**

Model B3 and all simulations for generation of every figure have been assembled in the model archive file (SM.B3).

The general idea used in the modelling is described below:

The reactions of synthesis and degradation of Antiox (re8 and re9), p62 (re6 and re7) and parkin (re10 and re11) were added to design 2B (see Supplemental Material SM.B2B) and set in the way to reach steady state concentrations of Antiox, p62 and parkin equal to their steady state concentrations in model B2B. A negative ROS-regulated feedback loop was added. ROS catalyze reaction 12 and shift the equilibrium towards a higher rate of keap1 oxidation (deactivation). Keap1 catalyzes reaction 14 and shifts the equilibrium towards the higher fraction of inactive Nrf2. Thus, ROS deactivate keap1 and activate Nrf2. When active, Nrf2 activates the expression of p62 (reaction 6) and the antioxidant response (reaction 8). The increase of ROS concentration activates ROS removal in antioxidant response, as well as p62-mediated removal of damaged mitochondria.

Computations were performed in ds. Comparing with previous models (B1, B2A and B2B) the ROS generation rate constant was decreased 4 fold to start simulations from the steady state with higher concentration of healthy mitochondria and lower concentration of damaged mitochondria and ROS (because in the following experiments the ROS generation rate constant increased several folds). Perturbations were performed using "time event" function in COPASI. The "factor time" was accumulating in time. At the moment "factor time" reached a certain level, the following events were triggered for Figure SMB3.1:
day 1 (factor time =10)-ROS synthesis is doubled (2 fold up from initial value);
day 3 (factor time =30)-ROS synthesis is doubled (4 fold up from initial value);
day 4 (factor time =40)-ROS synthesis is doubled (8 fold up from initial value);
day 5 (factor time =50)-ROS synthesis is doubled (16 fold up from initial value);
day 6 (factor time =60)- ROS synthesis is doubled (32 fold up from initial value).

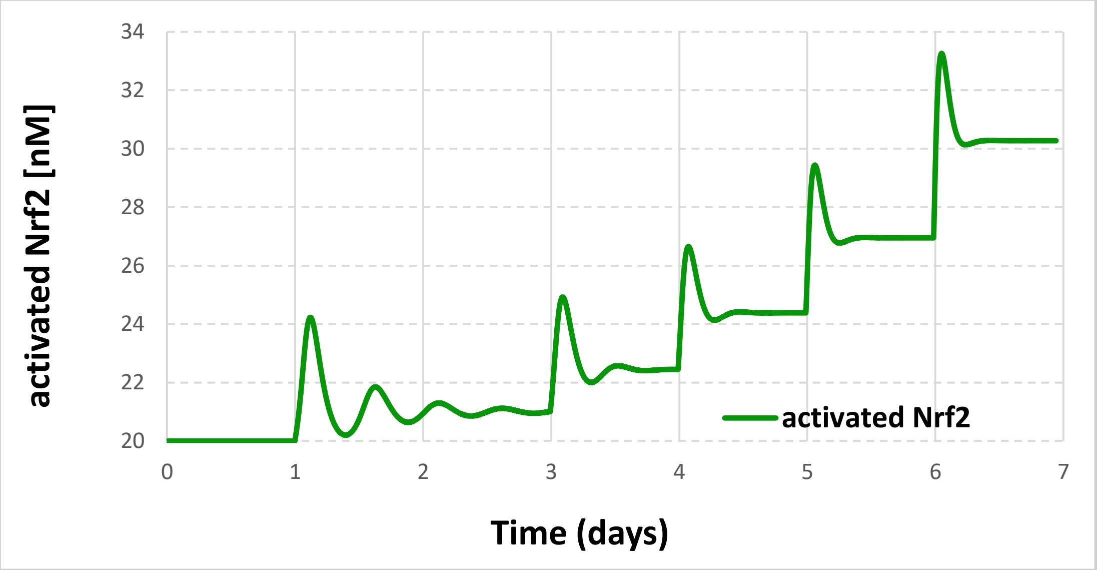

**Figure SM.B3.3. Design 3 (Model 3.cps). Oscillations of activated Nrf2. The** concentration of Nrf2 (in nM) was simulated. ROS synthesis was doubled stepwise: day 1-ROS synthesis was doubled comparing to initial steady state (2 fold increase from initial value); day 3 – ROS synthesis was doubled comparing to day 1 (4 fold increase from initial value); day 4 – ROS synthesis was doubled comparing to day 3 (8 fold up from initial value); Day 5 – ROS synthesis was doubled comparing to day 4 (16 fold up from initial value); Day 6 – ROS synthesis was doubled comparing to day 5 (32 fold up from initial value).

**The following conclusions were drawn from Figure SMB3.3:**

In the response to each perturbation, concentrations of activated Nrf2 transiently reaches a new steady state. At some range of the ROS generation rate constant, Nrf2 transiently oscillates.

#### SM.B4. Design 4. Model B4.

**A**

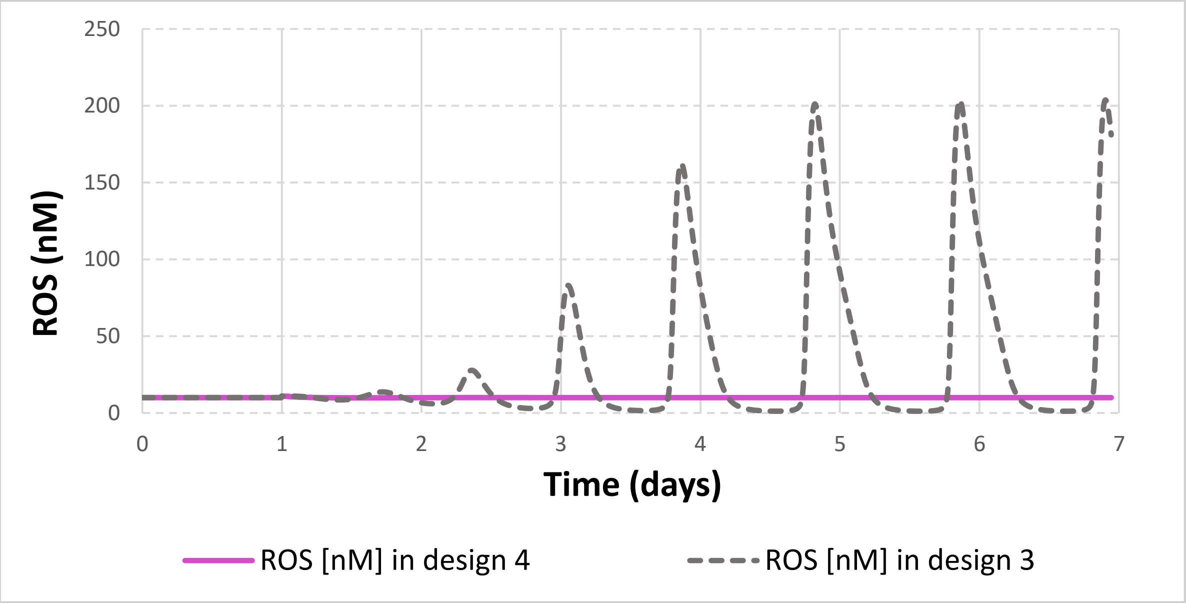

**B**

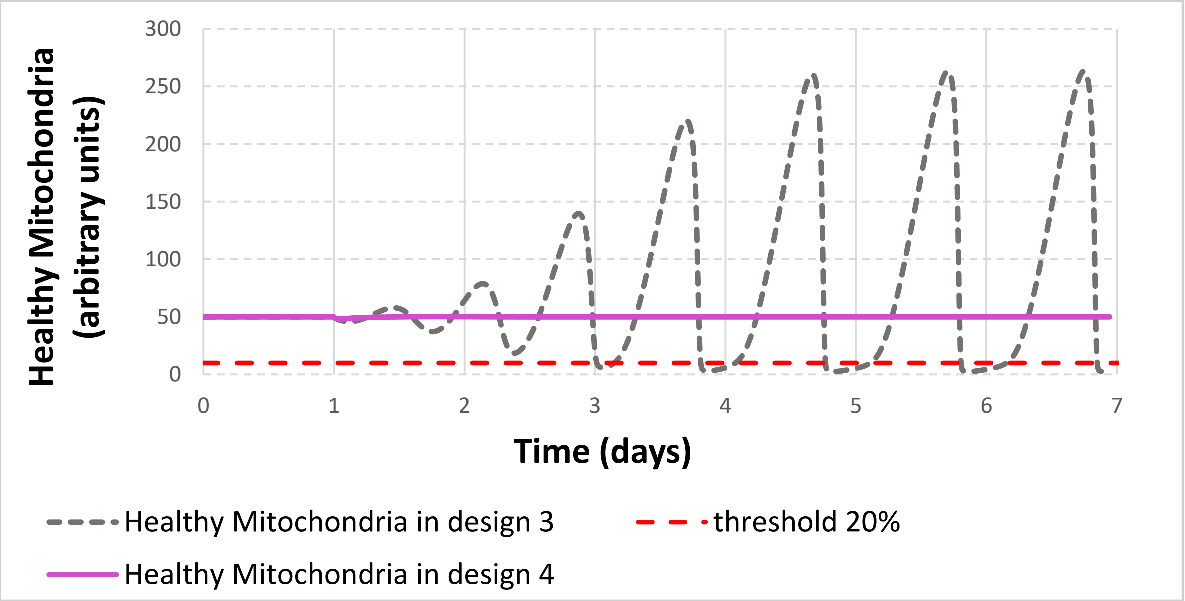

**Figure SM.B4.1. Design 4 (Model B4.cps). Dynamic stability obtained when NFκB signalling was added to design 3 thereby obtaining design 4.** Concentrations of (A) ROS (in nM) and (B) of healthy mitochondria (in arbitrary units) for design 4 (solid purple line) comparing with design 3 (dashed grey line). Simulations used models 3.cps and 4.cps. The Initial ROS concentration was perturbed (transient increase from 10 nM to 11 nM) at day 1.

**The following conclusions were drawn from Figure SM.B4.1:**

A: In design 3, the increase of ROS concentration from 10 nM to 11 nM at day 1 triggers an oscillatory behaviour of the ROS concentration. On the contrary, in design 4, ROS concentration increases transiently, but then quickly returns to the initial value.

B. In design 3, the increase of ROS concentration from 10 nM to 11 nM at day 1 triggers an oscillatory behaviour of ROS concentration. During oscillations, the concentration of healthy mitochondria sweeps below a hypothetical viability that corresponds to the threshold line (marked as a dotted red line) dissecting 20% (10 nM) of the initial concentration of healthy mitochondria (50 nM). On the contrary, in design 4, immediately upon perturbation, the concentration of healthy mitochondria drops from 50 nM to 48 nM, but then quickly returns to the initial value (50 nM) and does not sweep below a viability line (threshold line of 10 nM, shown as a dotted red line that dissects 20% of the initial concentration of healthy mitochondria).

We conclude that design 4 (comparing with design 3) gains dynamical stability against the perturbation of ROS concentration.

**A**

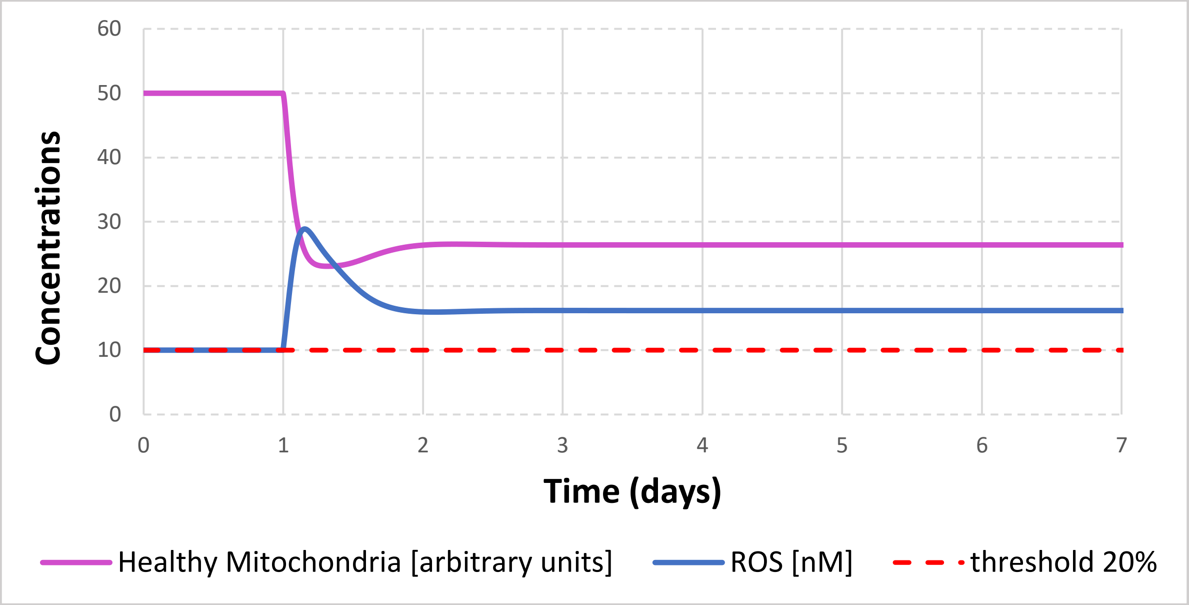

60
50
40
30
20
10
0
Concentrations
0
1
2
3
4
5
6
7
Time (days)
Healthy Mitochondria [arbitrary units]
ROS [nM]
threshold 20%

**B**

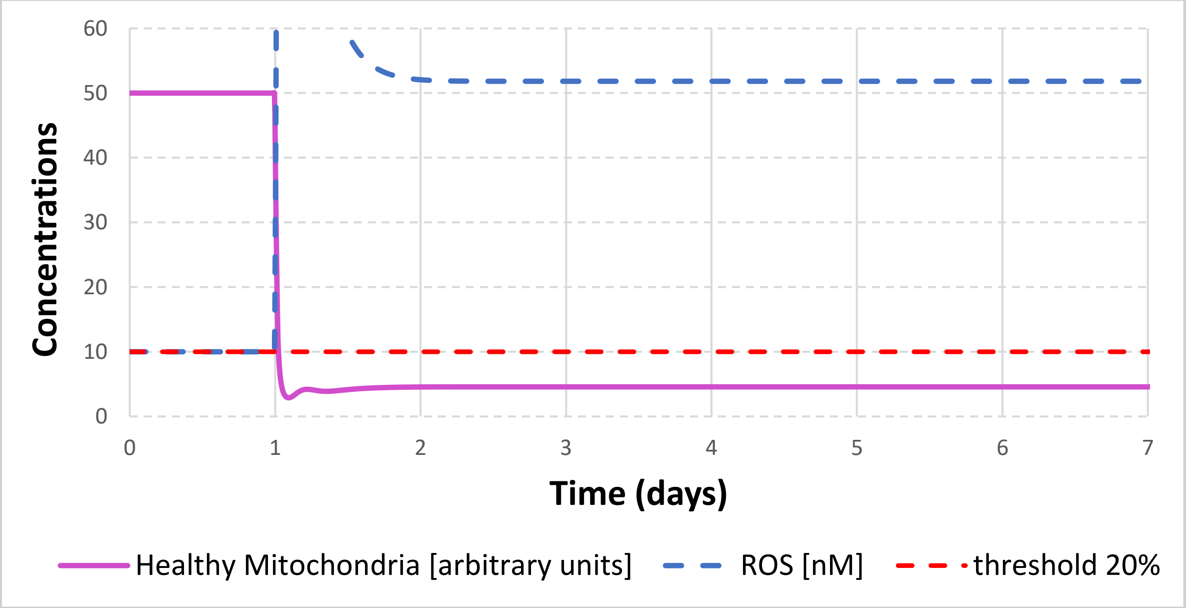

60
50
40
30
20
10
0
Concentrations
0
1
2
3
4
5
6
7
Time (days)
Healthy Mitochondria [arbitrary units]
ROS [nM]
threshold 20%

**C**

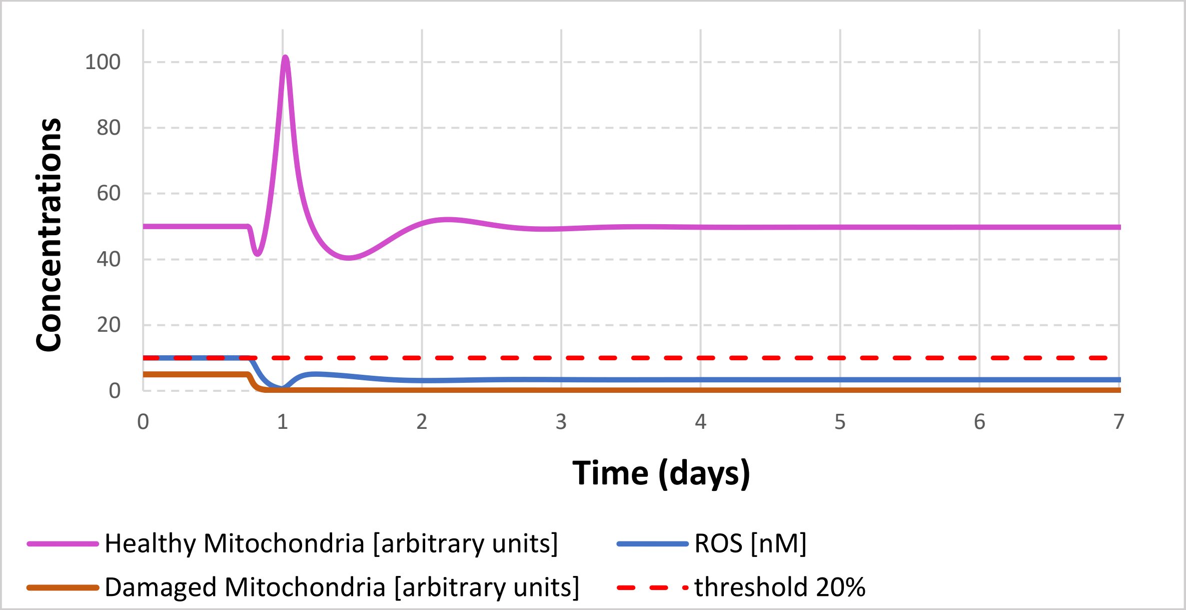
Concentrations
100
80
60
40
20
0
0
1
2
3
4
5
6
7
Time (days)
Healthy Mitochondria [arbitrary units]
ROS [nM]
Damaged Mitochondria [arbitrary units]
threshold 20%

**D**

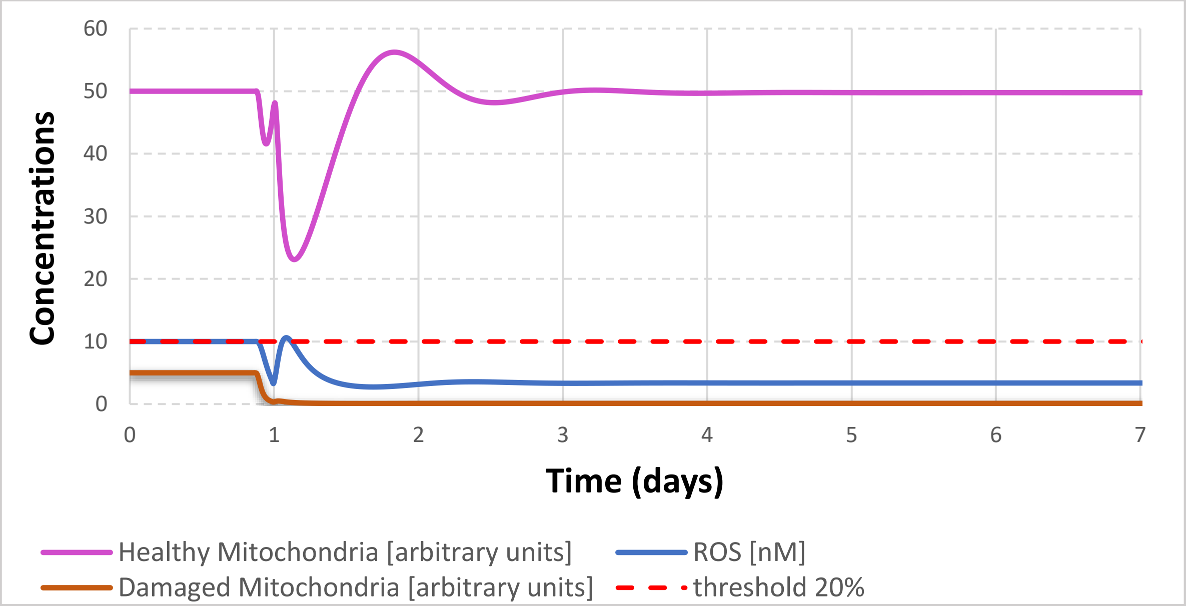
Concentrations
60
50
40
30
20
10
0
0
1
2
3
4
5
6
7
Time (days)
Healthy Mitochondria [arbitrary units]
ROS [nM]
Damaged Mitochondria [arbitrary units]
threshold 20%

**E**

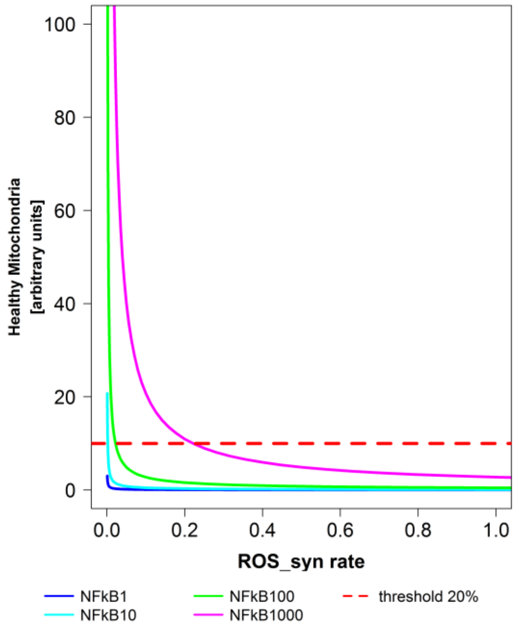

**Figure SM.B4.2. Design 4 (Model B4.cps). NFκB signalling protects from the increase of ROS generation in design 4.** Concentrations of ROS (in nM) and of healthy and damaged mitochondria (in arbitrary units) for design 4. Simulations used a model 4.cps. In A the ROS generation rate constant was increased 2 fold on day 1. In B the ROS generation rate constant was increased 15 fold on day 1. In C the ROS generation rate constant was increased 15 fold on day 1, but, 6h before the increase of ROS generation, the NFκB signalling was increased 15 fold. In D the ROS generation rate constant was increased 15 fold on day 1, but, 3h before the increase of ROS generation, the NFκB signalling was increased 15 fold. In E the steady state concentrations of healthy mitochondria (the ordinate) are shown for different rates of ROS generation (the abscissa) for 4 levels of NFκB signalling: (i) activated 1 fold (dark blue line); (ii) activated 10 fold (light blue line); (iii) activated 100 fold (green line); (iv) activated 1000 fold (purple line).

**The following conclusions were drawn from Figure SMB4.2:**

A: The concentration of ROS first increases, but then decreases and reaches the new steady state, with the level around 30% higher than the initial one. The concentration of healthy mitochondria decreases around 2 fold and persists at the new steady state level.

B: The concentration of healthy mitochondria drops below a viability line (the threshold line of 10 nM shown as a dotted red line that dissects 20% of the initial concentration of healthy mitochondria).

C: When NFκB signalling is increased, the damaged mitochondria (brown line) are converted into healthy mitochondria (purple line). The concentration of damaged mitochondria decreases. Thus, ROS concentration decreases as well. The increase of ROS synthesis stops this growth. ROS concentration increases again and the concentration of healthy mitochondria drops again. However, the concentration of healthy mitochondria does not drop below a viability line (the threshold line of 10 nM shown as a dotted red line that dissects 20% of the initial concentration of healthy mitochondria).

D: When NFκB signalling is increased, the damaged mitochondria (brown line) are converted into healthy mitochondria (purple line). The concentration of damaged mitochondria decreases. Thus, ROS concentration decreases as well. The increase of ROS synthesis stops this growth. ROS concentration increases again and the concentration of healthy mitochondria drops again. This dynamics is similar to the one described in C. However, the reduced time gap between the activation of NFκB signalling and the increase of ROS generation does not allow to achieve high peak of the concentration of healthy mitochondria. Nevertheless, at the end all concentrations reaches the same steady state values as in C. The concentration of healthy mitochondria does not drop below a viability line (the threshold line of 10 nM shown as a dotted red line that dissects 20% of the initial concentration of healthy mitochondria).

E. The higher is ROS generation, the lower is the steady state concentration of healthy mitochondria. The higher is NFκB signalling, the higher is the concentration of healthy mitochondria.

We conclude that the NFκB signalling helps to protect against the increase of ROS generation.

**A**

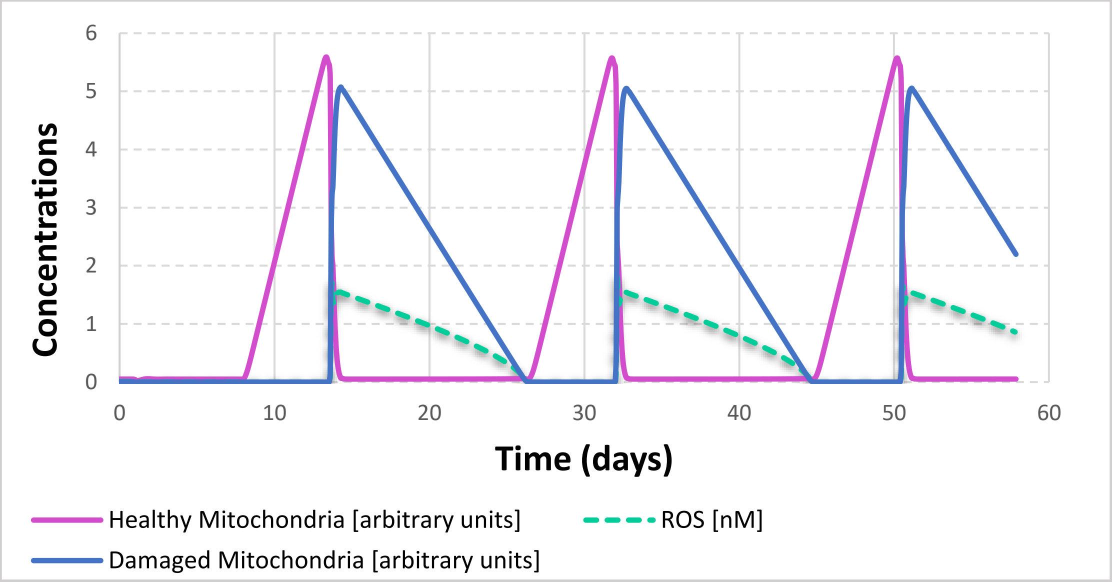
Concentrations
Time (days)
Healthy Mitochondria [arbitrary units]
ROS [nM]
Damaged Mitochondria [arbitrary units]

**B**

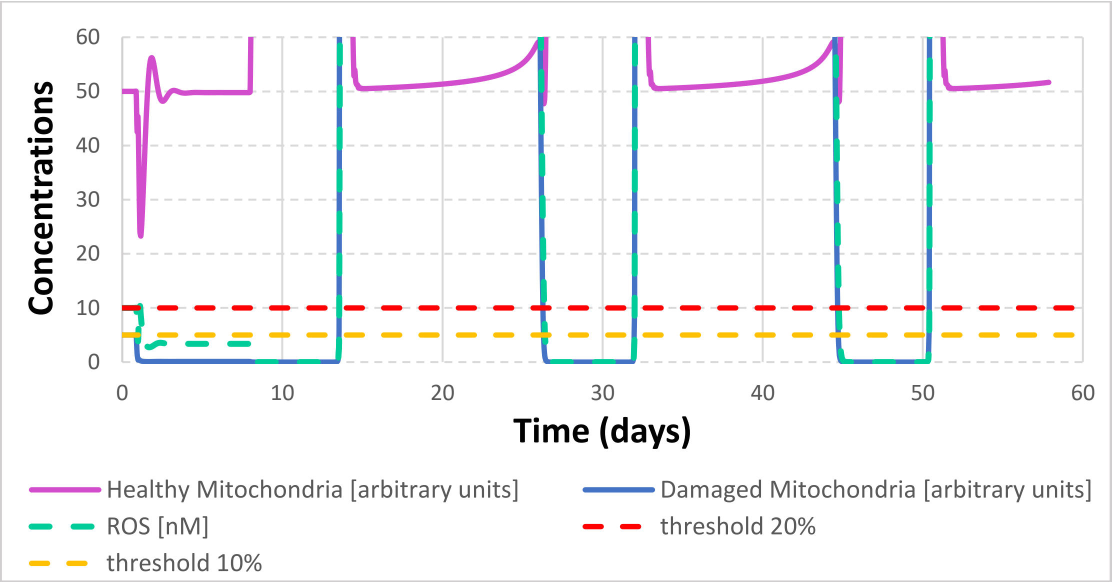
Concentrations
Time (days)
Healthy Mitochondria [arbitrary units]
Damaged Mitochondria [arbitrary units]
ROS [nM]
threshold 20%
threshold 10%

**C**

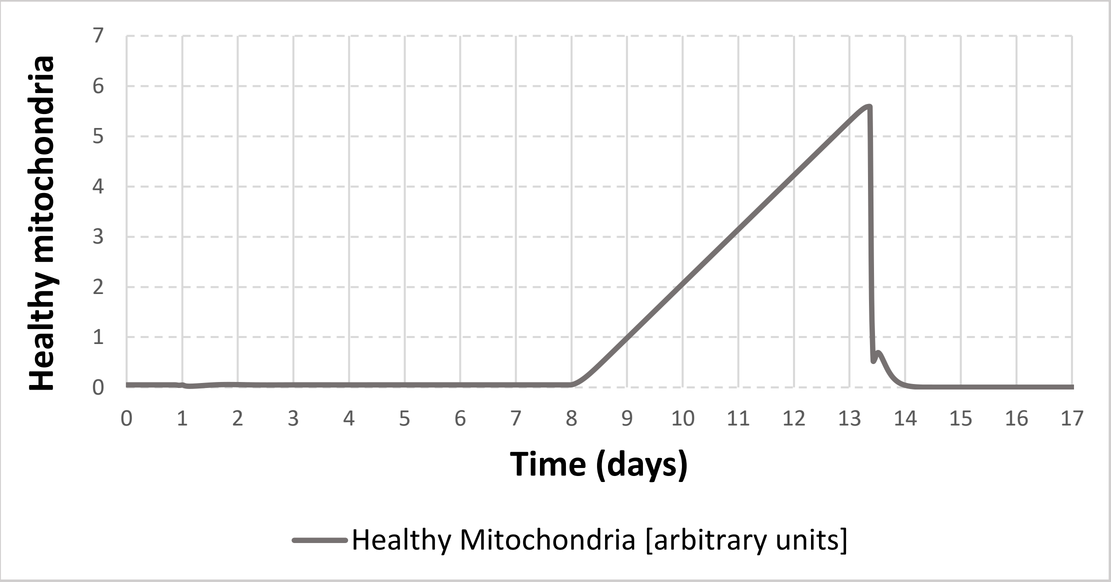

**D**

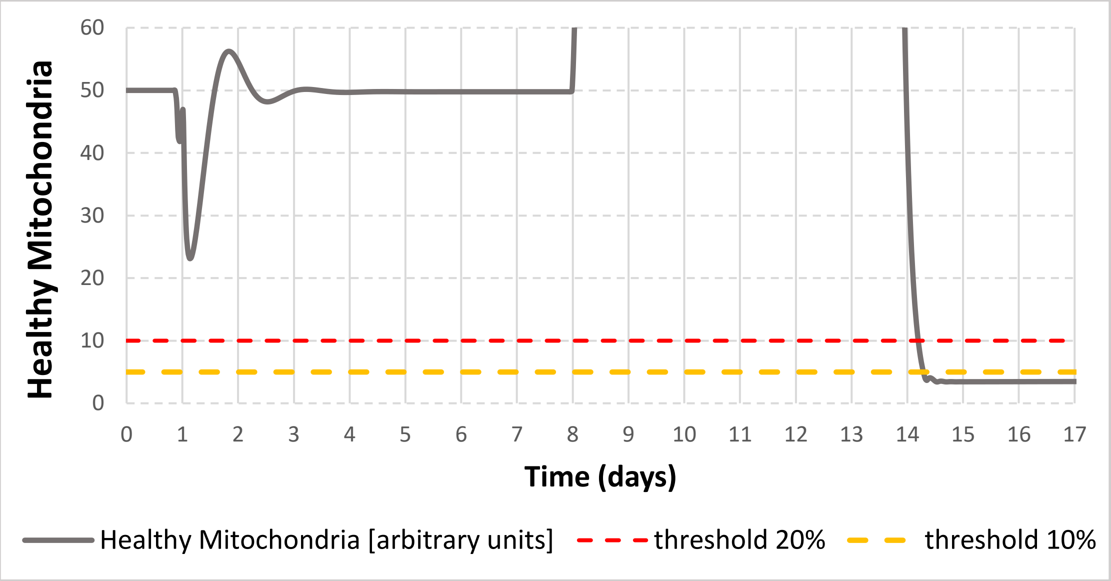

**Figure SM.B4.3. Design 4 (Model B4.cps). A potential danger of the NFκB-mediated accumulation of healthy mitochondria.** Concentrations of ROS (in nM) and of healthy and damaged mitochondria (in arbitrary units) for design 4. Simulations used a model 4.cps.

In A (bird-eye view) and B (high resolution view) the ROS generation rate constant was increased 15 fold on day 1, but, 3h before the increase of ROS generation, the NFκB signalling was increased 15 fold (the same perturbation as on Figure SM.B4.2D). When the system reached a steady state, the ROS generation rate constant was decreased 15 fold (on day 8) and oscillations started. In C (bird-eye view) and D (high resolution view): when a system was in an oscillatory mode (shown in A and B), the ROS generation rate constant was increased for the second time (on day 13.3). This was a time point when the concentration of healthy mitochondria was near its peak value.

**The following conclusions were drawn from Figure SM.B4.3:**

A: Upon the decrease of the ROS generation rate constant the system enters the oscillatory mode.

B: During the oscillation, the concentration of healthy mitochondria goes up and down the initial steady state value and does not sweep below a viability line (a threshold line of 10 nM shown as dotted red line that dissects 20% of the initial concentration of healthy mitochondria).

C: When the ROS generation rate constant is increased for the second time, the concentration of healthy mitochondria quickly goes down and oscillations stop.

D: When the concentration of healthy mitochondria goes down, it sweeps below a viability line (a threshold line of 10 nM shown as dotted red line that dissects 20% of the initial concentration of healthy mitochondria). We can note that the initial concentration of healthy mitochondria was at 50 nM. When the ROS generation rate constant is decreased, the concentration of healthy mitochondria goes up. However, when the ROS generation rate constant is increased for the second time, the concentration of healthy mitochondria sweeps much below the initial value (50 nM).

We conclude that a very high accumulation of healthy mitochondria creates a potential danger for the collapse when the ROS generation rate constant is increased for the second time. Design 4 is unstable against the second pulse of the increased ROS generation rate constant.

**Model building and simulations:**

Model B4 and all simulations for generation of every figure have been assembled in the model archive file (SM.B4).

The general idea used in the modelling is described below:

Model B4 was built by adding 5 additional reactions (with mass action kinetics) to model B3:

(re16): parkin activates NFκB signaling via IKK

(re17): the removal of NFκB signal

(re18): NFκB activates the expression of Bclxl

(re19): the degradation of Bclxl

(re20): Bclxl activates the recovery of mitochondria via biosynthesis.

The rate law of reaction 6 was modified in such a way that NFκB signal could activate p62 transcription: $v_6$= kf * NrfAct(t)*NFκB(t)*S, kf was fitted to reach steady state concentration of p62 equal to its concentration in model B3.

The rate constant of reaction 2 was adjusted in such a way that steady state concentrations of healthy and damaged mitochondria were equal to their steady state concentrations in model B3.

Computations were performed in ds, changes are shown in the scale of days. Perturbations were performed using “time event” function in COPASI.

Computations were performed in ds. Perturbations were performed using “time event” function in COPASI. The “factor time” was accumulating in time. At the moment “factor time” reached a certain level, the events were triggered.

#### SM.B5. Design 5. Models B5.1-B5.3A.

**A**

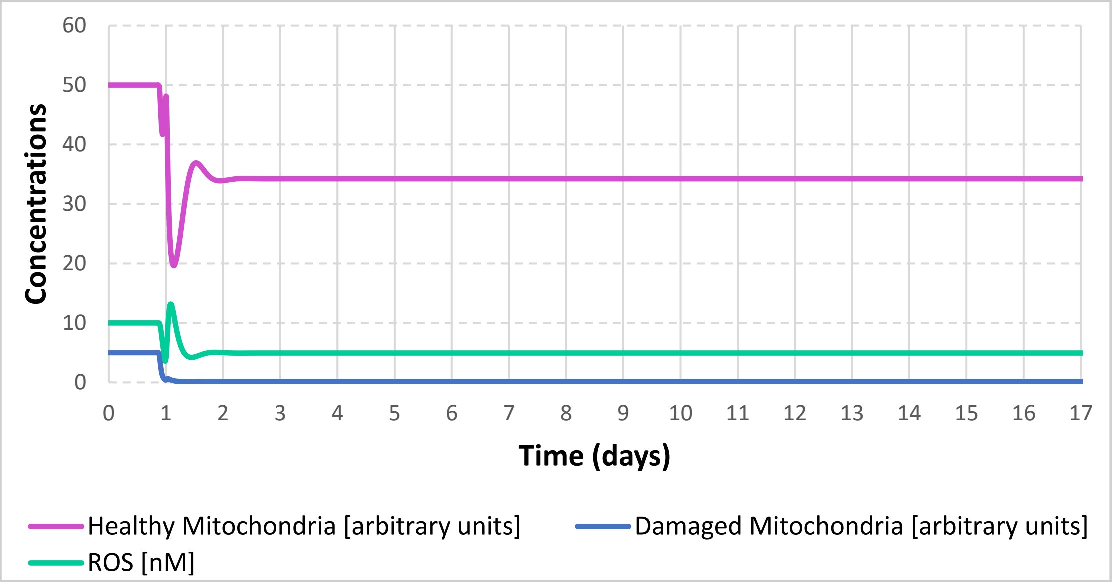

**B**

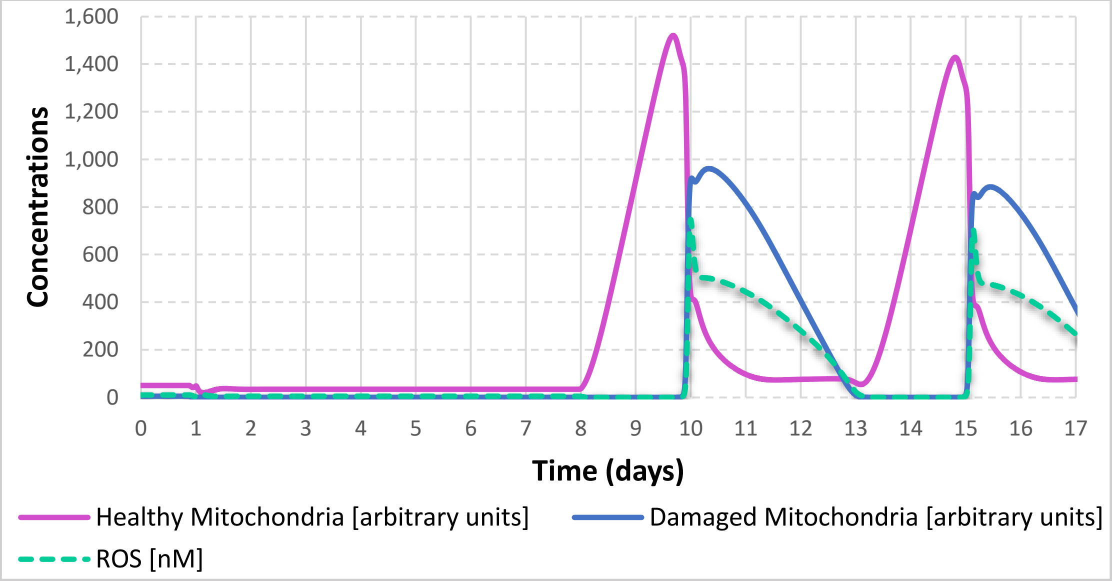

**C**

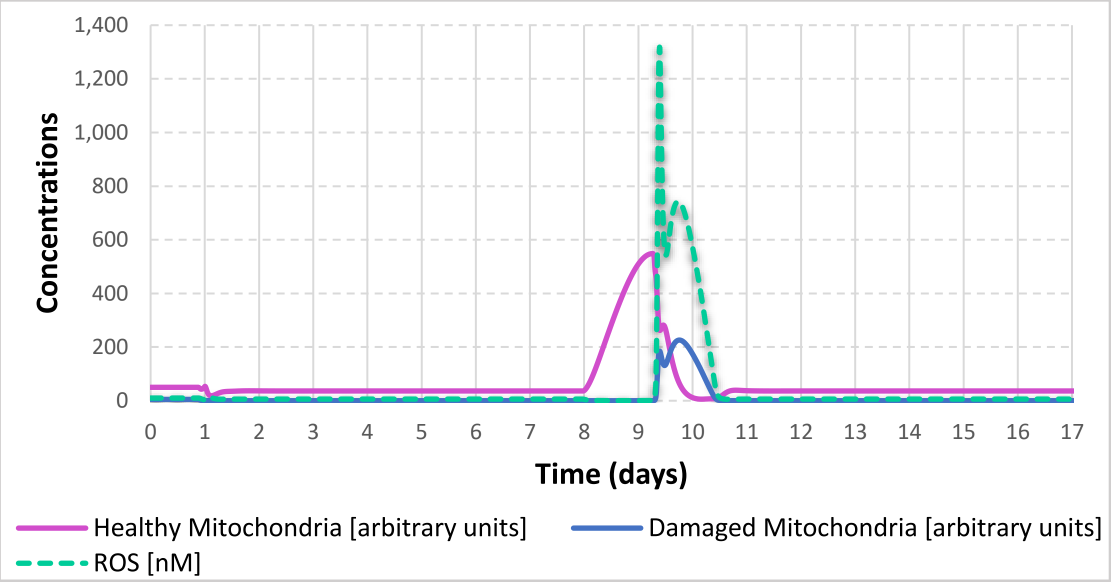

**D**

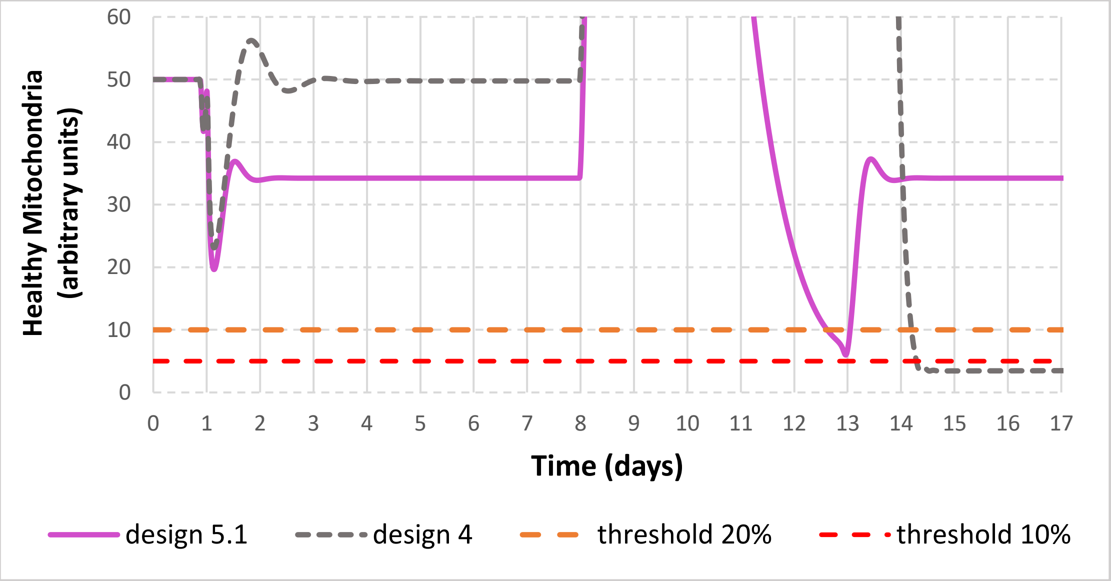

**Figure SM.B5.1. Design 5.1 (Model B5.1.cps). Robustness vis-à-vis with respect to the second pulse of ROS obtained when the regulation of NFkB signalling by ROS via DJ1 was added to design 4 thereby obtaining design 5.1.** Concentrations of ROS (in nM) and of healthy and damaged mitochondria (in arbitrary units) for design 5.1 (A-D) and design 4 (D). In A-D simulations model B5.1.cps was used. In D simulations model B5.1.cps was compared with model B4.cps.

In A, B, C and D the ROS generation rate constant was increased 15 fold on day 1, but, 3 hours before the increase of ROS generation, the NFκB signalling was increased 15 fold (the same perturbation as on Figures SM.B4.2D and SMB4.3). In A, the system reached a new steady state and no further perturbations were applied. In B, C and D the initial perturbations were followed by the decrease of ROS generation rate constant 15 fold on day 8. In C (bird-eye view) and D (high resolution view), when a system was in an oscillatory mode, the ROS generation rate constant was increased for the second time on day 9.8, at the time point when the concentration of healthy mitochondria was near its peak value.

**The following conclusions were drawn from Figure SM.B5.1:**

A: The system reaches a new steady state with the lower value of healthy mitochondria.

B: Upon the decrease of the ROS generation rate constant the system enters the oscillatory mode.

C: When the ROS generation rate constant is increased for the second time, the concentration of healthy mitochondria quickly goes down and oscillations stop.

D: In both model B4.cps (design 4) and model B5.1.cps (design 5.1) the concentration of healthy mitochondria sweeps below a viability line (a threshold line of 10 a.u. shown as a dotted red line that dissects 20% of the initial concentration of healthy mitochondria). However, in model B5.1.cps (design 5.1) the concentration of healthy mitochondria sweeps below a viability line only transiently and quickly recovers back to the initial level.

We conclude that the regulation of NFκB signalling by ROS via DJ1 provides limited robustness vis-à-vis with respect to the second pulse of ROS.

A

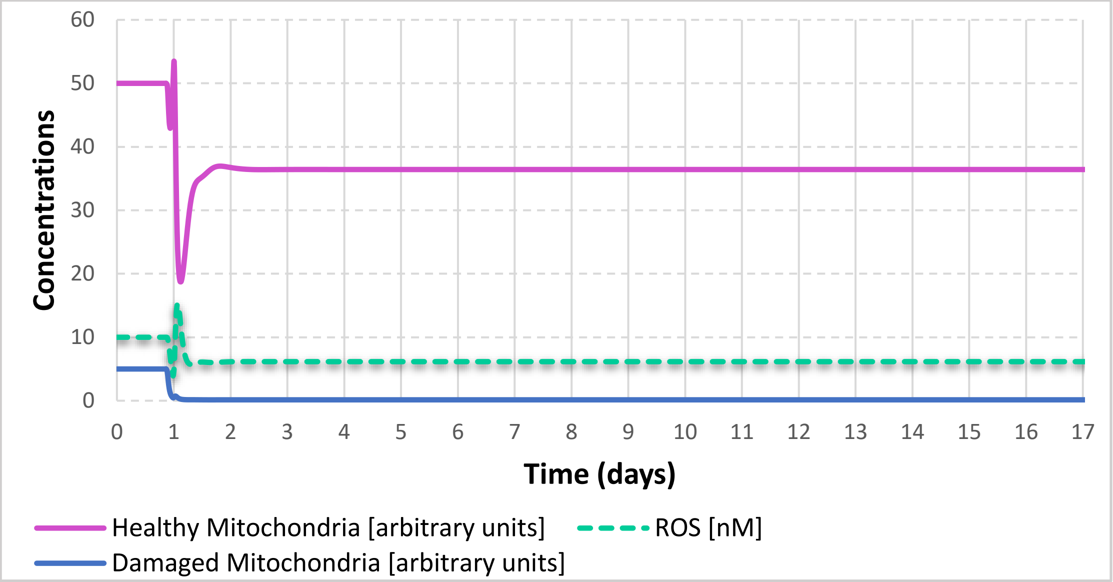

60
50
40
30
20
10
0
Concentrations
Time (days)
Healthy Mitochondria [arbitrary units]
ROS [nM]
Damaged Mitochondria [arbitrary units]

B

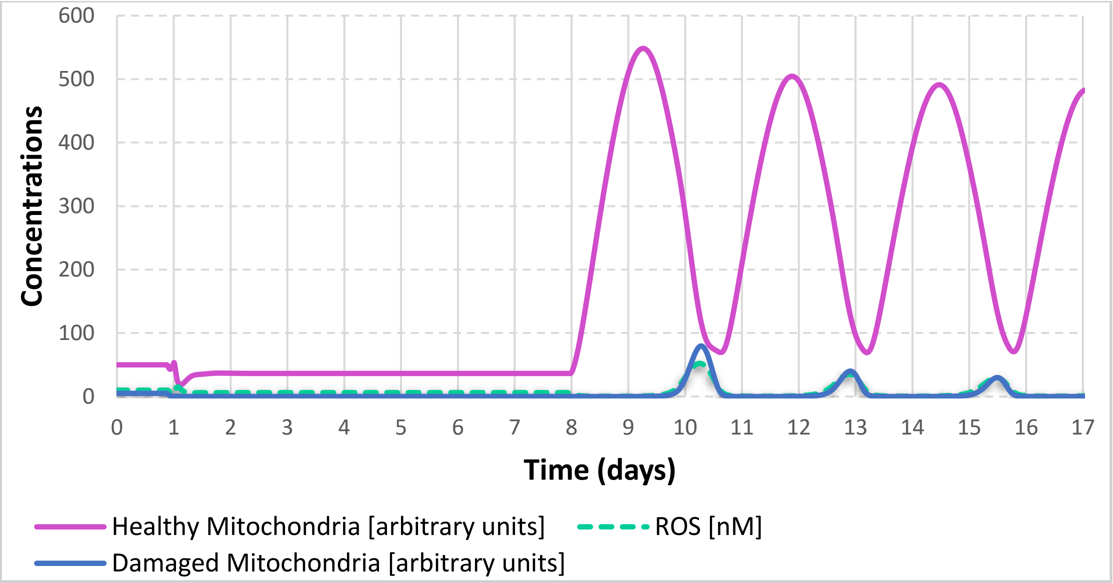

600
500
400
300
200
100
0
Concentrations
Time (days)
Healthy Mitochondria [arbitrary units]
ROS [nM]
Damaged Mitochondria [arbitrary units]

**C**

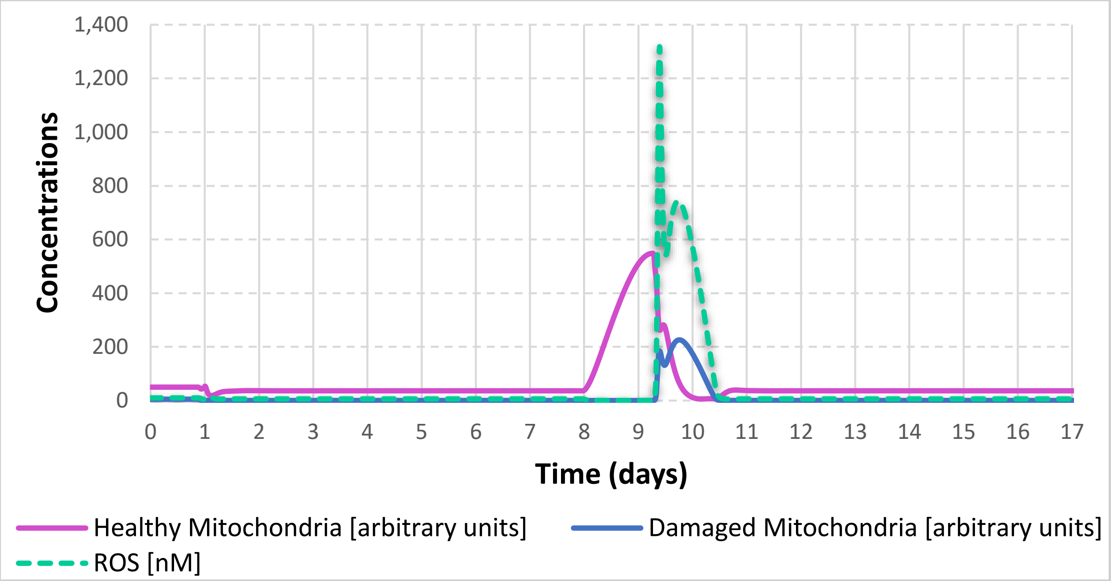

**D**

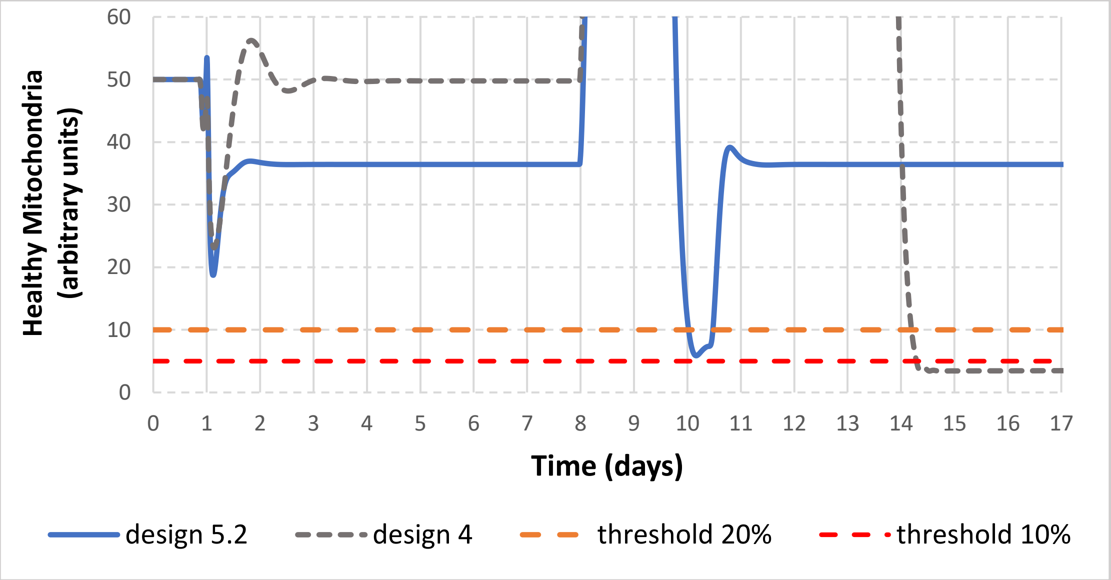

**Figure SM.B5.2. Design 5.2 (Model B5.2.cps). Robustness vis-à-vis with respect to the second pulse of ROS obtained when the regulation of Nrf2 signalling by ROS via DJ1 was added to design 4 thereby obtaining design 5.2.** Concentrations of ROS (in nM) and of healthy and damaged mitochondria (in arbitrary units) for design 5.2 (A-D) and design 4 (D). In A-D simulations model B5.2.cps was used. In D simulations model B5.2.cps was compared with model B4.cps.

In A, B, C and D the ROS generation rate constant was increased 15 fold on day 1, but, 3h before the increase of ROS generation, the NFκB signalling was increased 15 fold (the same perturbation as on Figures SM.B4.2D, SM.B4.3 and SM.B5.1). In A, the system reached a new steady state and no further perturbations have been made. In B, C and D the initial perturbations were followed by the decrease of ROS generation rate constant 15 fold on day 8. In C (bird-eye view) and D (high resolution view), when a system was in an oscillatory mode, the ROS generation rate constant was increased for the second time on day 9.3, at the time point when the concentration of healthy mitochondria was near its peak value.

**The following conclusions were drawn from Figure SM.B5.2:**

A: The system reaches a new steady state with the lower value of healthy mitochondria.

B: Upon the decrease of the ROS generation rate constant the system enters the oscillatory mode.

C: When the ROS generation rate constant is increased for the second time, the concentration of healthy mitochondria quickly goes down and oscillations stop.

D: In both models B4.cps (design 4) and model B5.2.cps (design 5.2) the concentration of healthy mitochondria sweeps below a viability line (a threshold line of 10 a.u. shown as a dotted red line that dissects 20% of the initial concentration of healthy mitochondria). However, in model B5.2.cps (design 5.2) the concentration of healthy mitochondria sweeps below a viability line only transiently and quickly recovers back to the initial level.

We conclude that the regulation of Nrf2 signalling by DJ1 provides limited robustness vis-à-vis with respect to the second pulse of ROS.

**A**

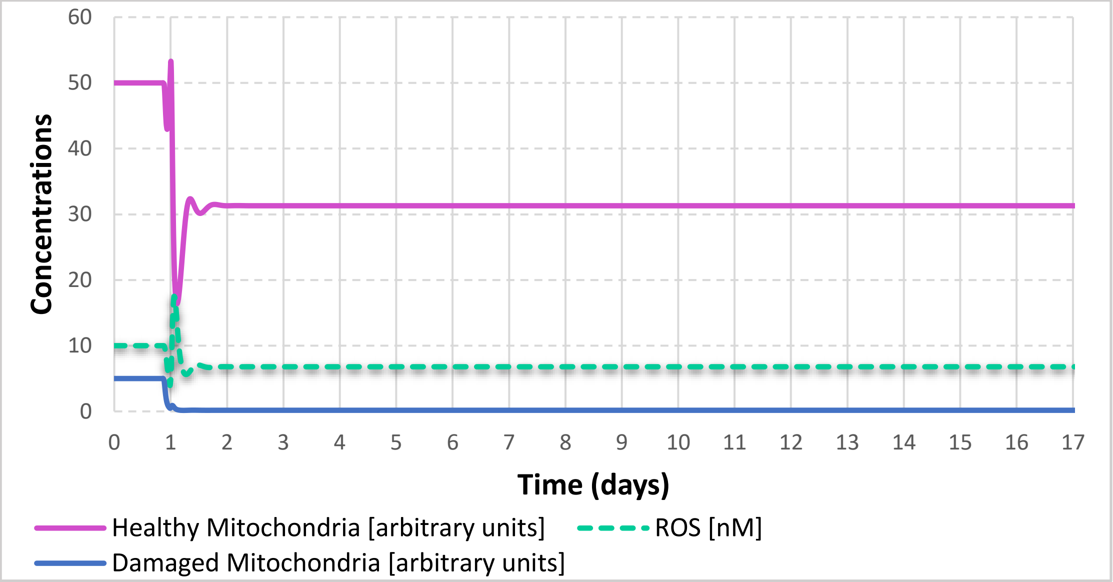

60
50
40
30
20
10
0
Concentrations
0 1 2 3 4 5 6 7 8 9 10 11 12 13 14 15 16 17
Time (days)
Healthy Mitochondria [arbitrary units]
ROS [nM]
Damaged Mitochondria [arbitrary units]

**B**

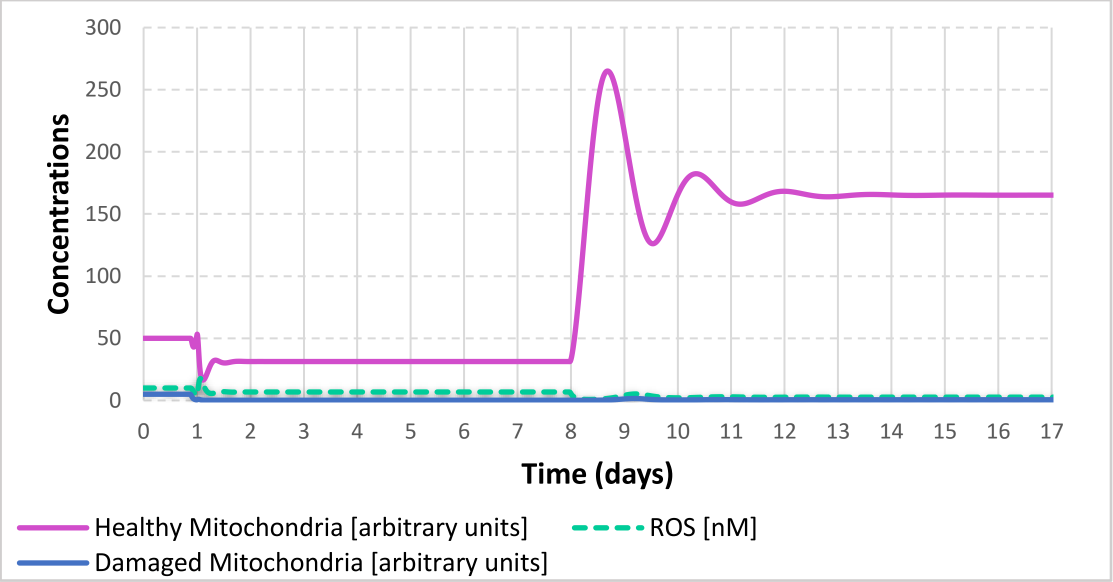

300
250
200
150
100
50
0
Concentrations
0 1 2 3 4 5 6 7 8 9 10 11 12 13 14 15 16 17
Time (days)
Healthy Mitochondria [arbitrary units]
ROS [nM]
Damaged Mitochondria [arbitrary units]

**C**

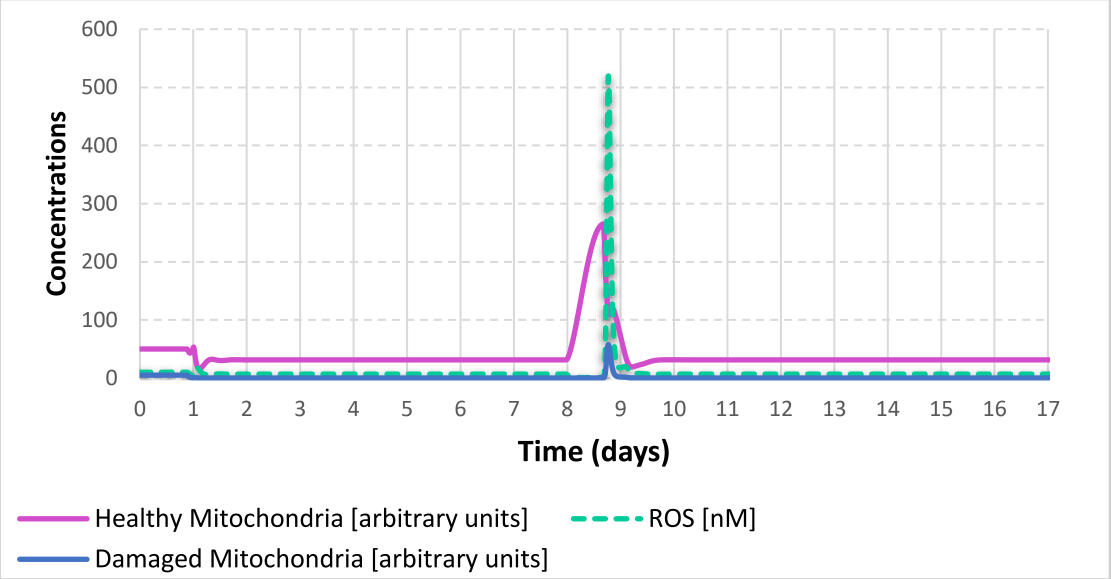

**D**

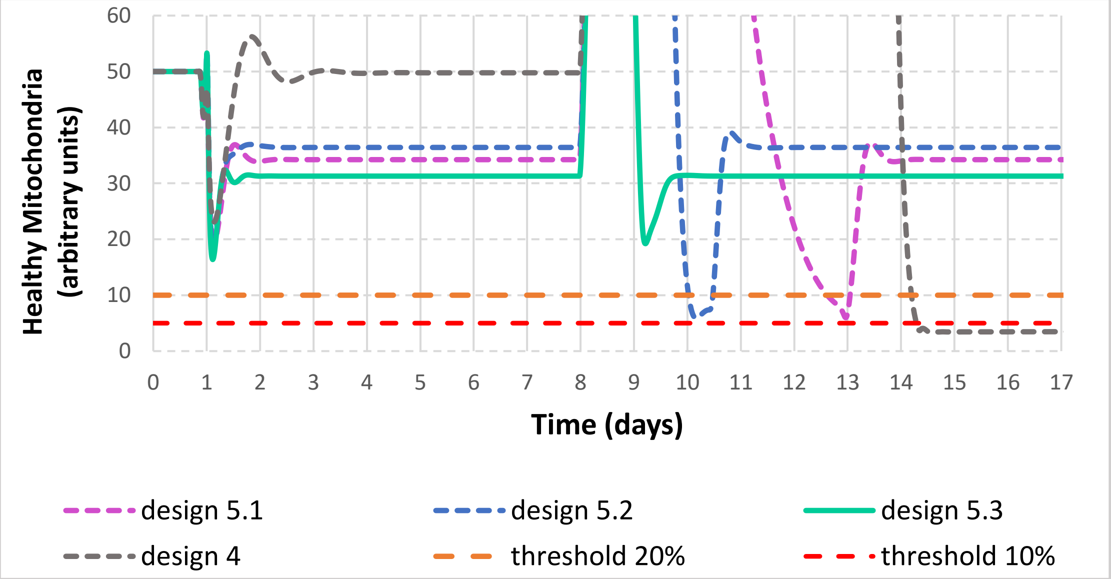

**Figure SM.B5.3. Design 5.3 (Model B5.3.cps). Robustness vis-à-vis with respect to the second pulse of ROS obtained when the regulation of both NFκB and Nrf2 signalling by DJ1 was added to design 4 thereby obtaining design 5.3.** Concentrations of ROS (in nM) and of healthy and damaged mitochondria (in arbitrary units) for design 5.3 (A-D) and design 4 (D). In A-D simulations model B5.3.cps was used. In D model B5.2.cps was compared with models B4.cps, B5.1.cps and B5.3.cps.

In A, B, C and D the ROS generation rate constant was increased 15 fold on day 1, but, 3h before the increase of ROS generation, the NFκB signalling was increased 15 fold (the same perturbation as on Figures SM.B4.2D, SM.B4.3, SM.B5.1 and SM.B5.2). In A, the system reached a new steady state and no further perturbations have been made. In B, C and D the initial perturbations were followed by the decrease of ROS generation rate constant 15 fold on day 8. In C (bird-eye view) and D (high resolution view), when a system was in an oscillatory mode, the ROS generation rate constant was increased for the second time on day 8.7, at the time point when the concentration of healthy mitochondria was near its peak value.

**The following conclusions were drawn from Figure SM.B5.3:**

A: The system reaches a new steady state with the lower value of healthy mitochondria (31 a.u.).

B: Upon the decrease of the ROS generation rate constant the system exhibites several dumping oscillations and reaches a new steady state with the higher concentration of healthy mitochondria (165 a.u.).

C: When the ROS generation rate constant is increased for the second time, the concentration of healthy mitochondria quickly goes down to the steady state value of 35 a.u.

D: In models B4.cps, B5.1.cps and B5.2.cps the concentration of healthy mitochondria sweeps below a viability line (a threshold line of 10 a.u. shown as a dotted red line that dissects 20% of the initial concentration of healthy mitochondria). However, in model B5.3.cps (design 5.3) the concentration of healthy mitochondria does not sweep below a viability line.

We conclude that the regulation of both NFκB and Nrf2 signalling by ROS via DJ1 provides robustness vis-à-vis with respect to the second pulse of ROS.

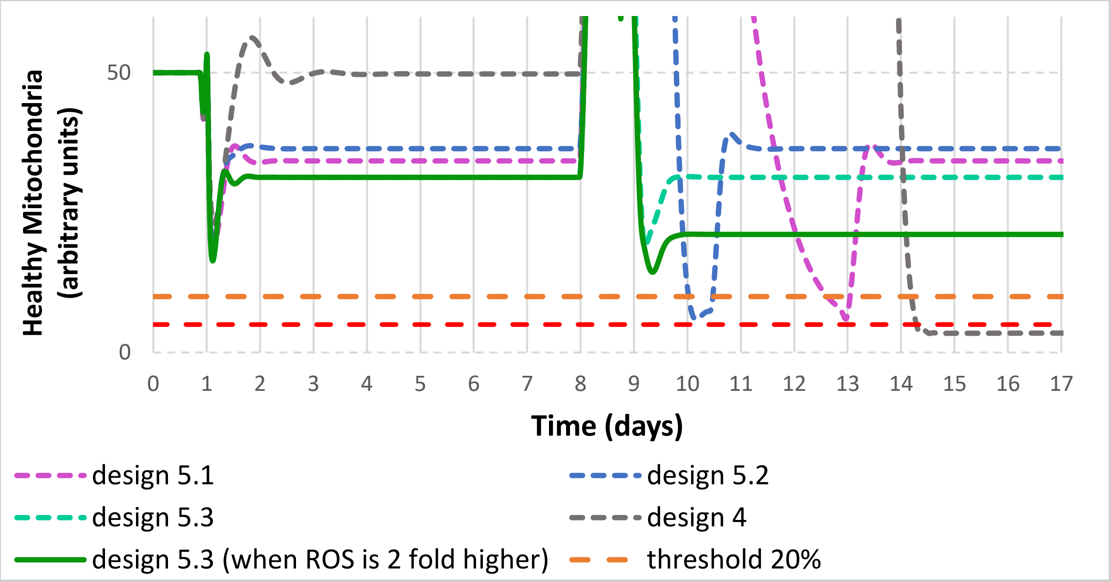

**Figure SM.B5.4. Design 5.3 (Model B5.3.cps). Robustness vis-à-vis with respect to the higher second pulse of ROS.** Concentrations of ROS (in nM) and of healthy and damaged mitochondria (in arbitrary units). Models B4, B5.1, B5.2 and B5.3.cps were compared.
The ROS generation rate constant was increased 15 fold on day 1, but, 3h before the increase of ROS generation, the NFκB signalling was increased 15 fold (the same perturbation as on Figures SM.B4.2D, SM.B4.3, SM.B5.1, SM.B5.2 and SM.B5.3). The initial perturbations were followed (on day 8) by the decrease of ROS generation rate constant 15 fold that resulted in the increase of the concentration of healthy mitochondria. At the time point when the concentration of healthy mitochondria was near its peak value, in each model, the ROS generation rate constant was increased for the second time either 15 fold (Models B4-dotted grey line; B5.1-dotted purple line; B5.2-dotted green line; B5.3-solid light green line), or 30 fold (Model B.3-dark green line).

**The following conclusions were drawn from Figure SM.B5.4:**

In models B4 (design 4), B5.1 (design 5.1), and B5.2 (design 5.2) the concentration of healthy mitochondria sweeps below a viability line (a threshold line of 10 a.u. shown as a dotted red line that dissects 20% of the initial concentration of healthy mitochondria). However, in model B5.3 (design 5.3) the concentration of healthy mitochondria does not sweep below a viability line, even if the ROS generation rate constant is increased 30 instead of 15 fold.

We conclude that the regulation of both NFκB and Nrf2 signalling by ROS via DJ1 (Model 5.3, design 5.3) provides strong robustness vis-à-vis with respect to the second pulse of ROS, even for the higher ROS challenge.

A

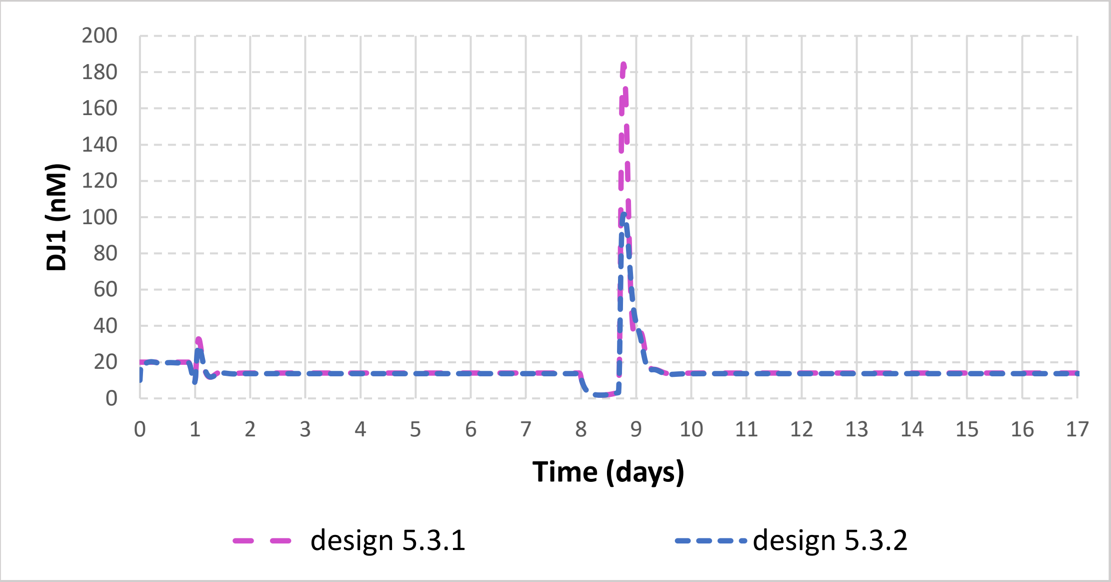

B

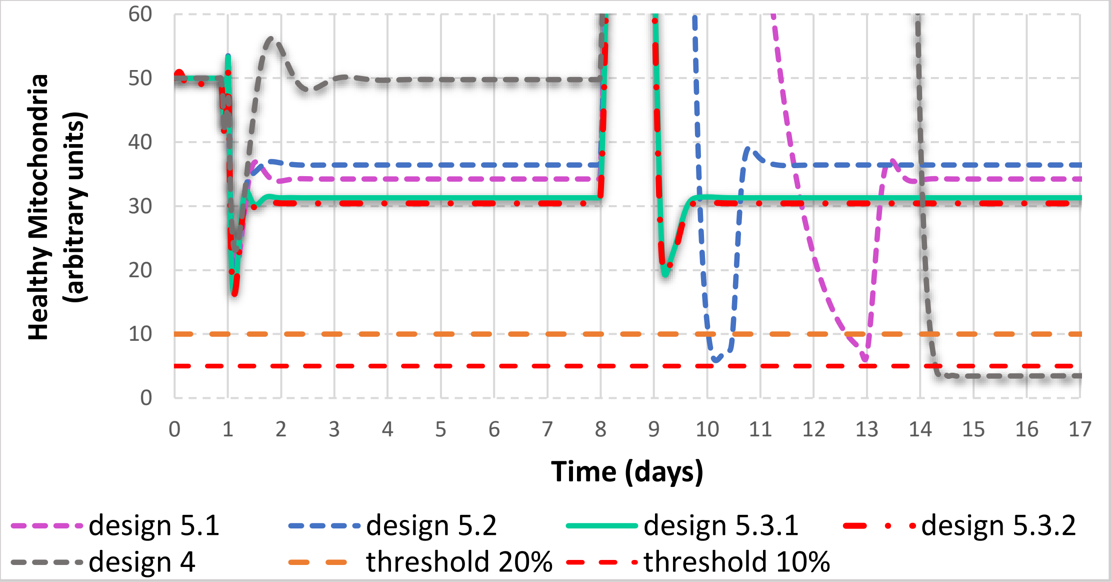

**Figure SM.B5.5. Design 5.3 (intact models B5.3.1.cps and the model with reduced DJ1 activity B5.3.2.cps) comparing with design 4 (model B4.cps), design 5.1 (model B5.1.cps) and design 5.2 (model B5.2.cps).** Concentrations of ROS (in nM) and of healthy and damaged mitochondria (in arbitrary units). In A the activity of DJ1 in the intact model (model B5.3.1) and in the model with reduced DJ1 activity (model B5.3.2.cps) is shown. In B, the concentration of healthy mitochondria for models B.4.cps, B5.1.cps, B5.2.cps, B5.3.1.cps and B5.3.2.cps (with reduced activity of DJ1) is shown. The ROS generation rate constant was increased 15 fold on day 1, but, 3h before the increase of ROS generation, the NFκB signalling was increased 15 fold (the same perturbation as on Figures SM.B4.2D, SM.B4.3, SM.B5.1, SM.B5.2, SM.B5.3 and SM.B5.4). The initial perturbations were followed (on day 8) by the decrease of ROS generation rate constant 15 fold that resulted in the increase of the concentration of healthy mitochondria. At the time point when the concentration of healthy mitochondria was near its peak value, in each model, the ROS generation rate constant was increased for the second time 15 fold.

**The following conclusions were drawn from Figure SM.B5.5:**

A: The response in DJ1 activation in model 5.3.1.cps is twice lower than in the model 5.3.2.cps.

B: In models B4 (design 4), B5.1 (design 5.1), and B5.2 (design 5.2) the concentration of healthy mitochondria sweeps below a viability line (a threshold line of 10 a.u. shown as a dotted red line that dissects 20% of the initial concentration of healthy mitochondria). However, in model B5.3 (design 5.3), for both intact DJ1 activity (model B5.3.1.cps) and for model with the reduced DJ1 activity (model B5.3.2) the concentration of healthy mitochondria does not sweep below a viability line.

We conclude that the regulation of both NFκB and Nrf2 signalling by ROS via DJ1 (design 5.3) provides strong robustness vis-à-vis with respect to the second pulse of ROS, even for lower DJ1 activity.

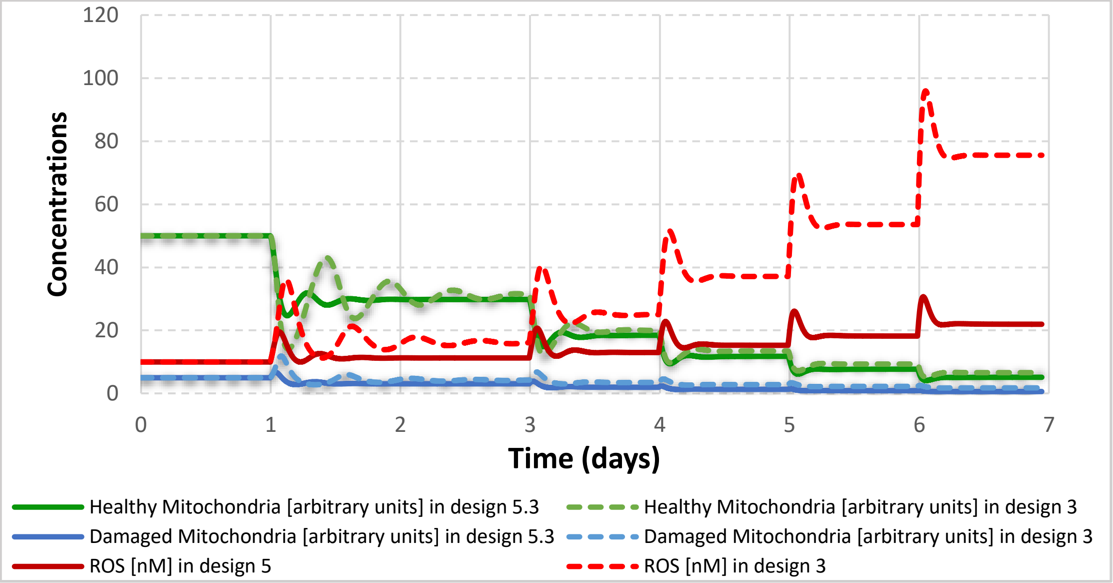

**Figure SM.B5.6. Design 5.3 comparing with design 3 (model B3.cps, Figure SM.B3.1).**

**Strong homeostasis obtained in design 5 where both NFκB and Nrf2 loops were sensitive to DJ1.** Concentrations of ROS (in nM) and healthy and damaged mitochondria (in arbitrary units) for design 3 and design 5. Simulations used a model B3.cps and B5.3.cps. ROS synthesis was doubled stepwise: day 1-ROS synthesis was doubled comparing to initial steady state (2 fold increase from initial value); day 3 – ROS synthesis was doubled comparing to day 1 (4 fold increase from initial value); day 4 – ROS synthesis was doubled comparing to day 3 (8 fold up from initial value); Day 5 – ROS synthesis was doubled comparing to day 4 (16 fold up from initial value); Day 6 – ROS synthesis was doubled comparing to day 5 (32 fold up from initial value).

**The following conclusions were drawn from Figure SMB5.6:**

In the response to each perturbation, concentrations of ROS and healthy mitochondria transiently reach a new steady state. Upon every doubling of ROS synthesis, ROS concentration is first increased twice, but then is decreased back to a value a bit higher than the initial one. Upon every perturbation, the concentration of healthy mitochondria first decrease around 2 fold, but then increase back to the value a bit lower than the steady state value before perturbation.
We conclude that the system actively counteracts the increase of ROS. The homoeostasis in design 5 (model B5.3.cps) is stronger than in design 4 (model B3.cps)

**Model building and simulations:**

Models B5.1, B5.2, B5.3.1, B5.3.2, and all simulations for generation of every figure have been assembled in the model archive file (SM.B5).

The general idea used in the modelling is described below:

Models B5.1, B5.2 and B5.3 were built by adding the two forms of DJ1 protein (DJ1active and DJ1inactive) to model B4.

DJ1 was activated (oxidized by ROS) in irreversible mass action reaction 21: v=kf*ROS(t)*DJ1inact(t)

DJ1 was inactivated in irreversible mass action reaction 22: v=kf*DJ1act(t).

In model B5.1, DJ1active inhibited the inactivation of NFκB signaling (re 17): v=S*(Vm/(Km+DJ1act(t))), Vm and Km were fit in such a way as to maintain the same steady state concentration of NFκB as in model B4.

In model B5.2, DJ1active inhibited the inactivation of Nrf2 (re14): v= S*Vm*Keap1act/(Km+ DJ1act(t)), the rate constants were adjusted in such a way as to maintain the same steady state concentrations of Nrf2act and Nrf2inactive as in model B4.

In model B5.3.1 (the complete design 5) DJ1active inhibited both inactivation of NFκB (re17) and inactivation of Nrf2 (re14).

First, concentrations of DJ1 active and inactive were fixed (DJ1 was insensitive to ROS) and model was checked to behave identically to model B4.

In model B5.3.2 derived from model B5.3.1 the capacity of DJ1 activation was reduced 2 fold. In order to obtain the reduced DJ1 activity, the total concentration of DJ1 was reduced two fold, but the rate of the reaction of DJ1 inactivation was also reduced 2 fold (to obtain the same initial steady state).

Computations were performed in ds. Perturbations were performed using "time event" function in COPASI. The "factor time" was accumulating in time. At the moment "factor time" reached a certain level, the events were triggered.

#### SM.SM.C. Design C. Model C.

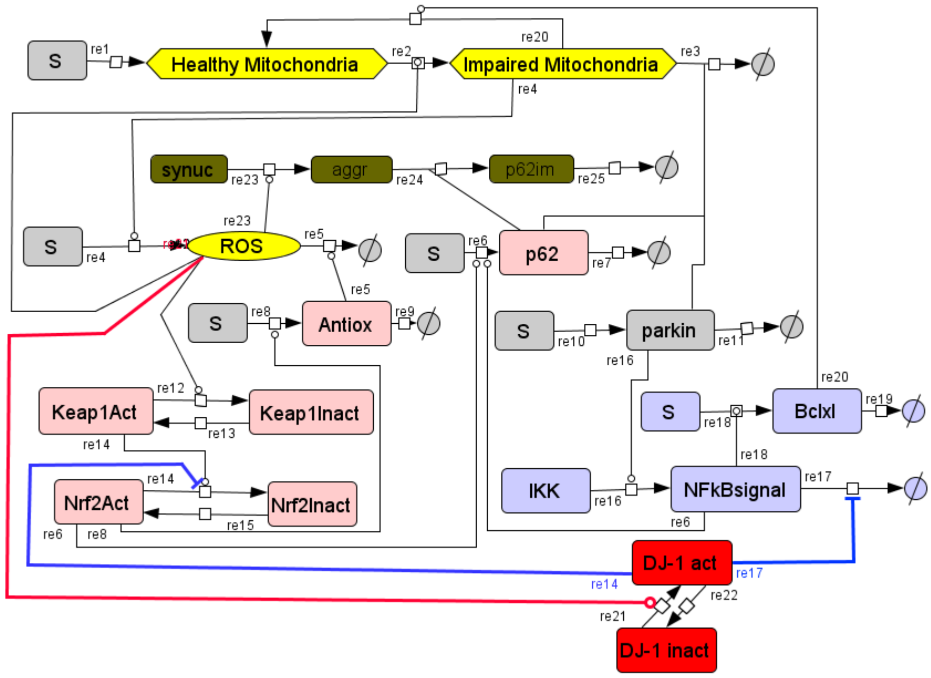

**Figure SM.C.1. Design C (Model C). Network diagram of Model C including the α-synuclein module.**

Model C was derived from Model B5 by incorporating a constant source of α-synuclein, ROS-dependent polymerisation of α-synuclein (reversible mass action reaction 23), sequestration of p62 by α-synuclein aggregates (reversible mass action reaction 24) and degradation of α-synuclein aggregates with p62 (irreversible mass action reaction 25).

**A**

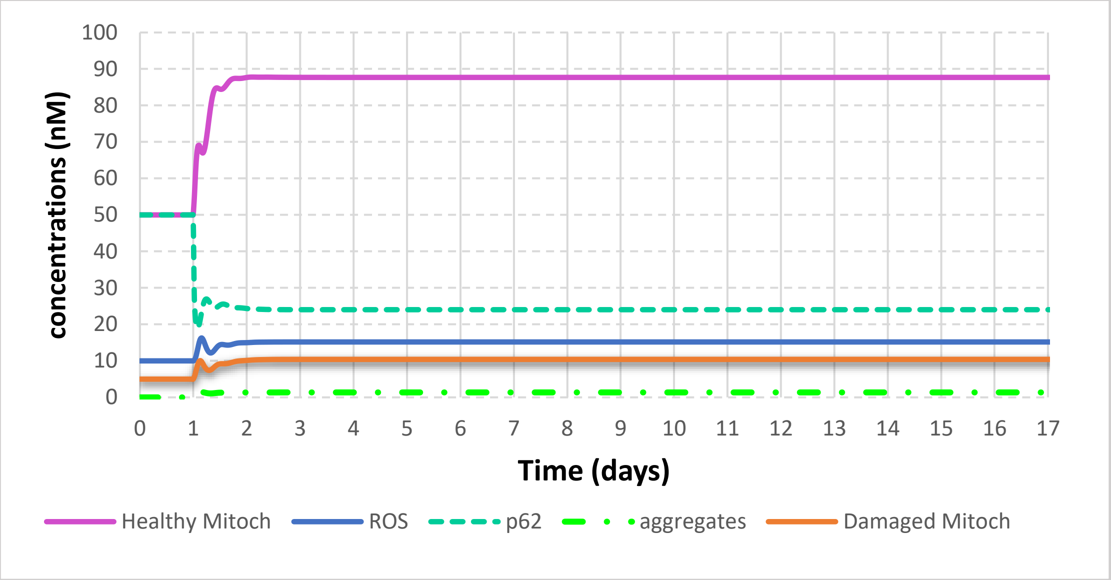

**B**

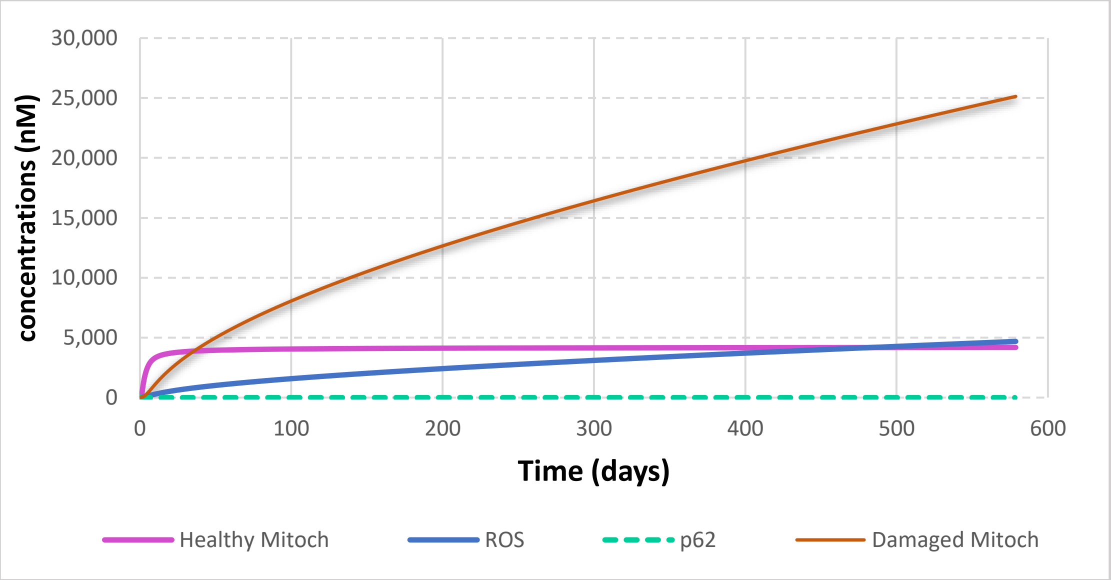

**Figure SM.C.2. Design C (Model C). The double role of α-synuclein.** Concentrations of ROS and p62 (in nM), of α-synuclein aggregates and of healthy and damaged mitochondria (in arbitrary units) for design C. First, α-synuclein was absent and system was at a steady state. On day 1, the concentration of α-synuclein was increased. Oxidation of α-synuclein by ROS caused the formation of α-synuclein aggregates. Then, α-synuclein aggregates sequestered p62. This process reduced mitophagy and helped to increase the concentration of healthy mitochondria. In A, the concentration of α-synuclein on day 1 was increased from 0 till 0.1 a.u. In B, the concentration of α-synuclein on day 1 was increased from 0 till 0.1 a.u.

**The following conclusions were drawn from Figure SM.C.2:**

A) The system reaches a new steady state with the concentration of healthy mitochondria approximately 80% higher than the initial one. The concentration of damaged mitochondria and ROS increases approximately 50%.
B) The concentration of damaged mitochondria constantly increases and the system explodes (as it is no longer able to reach a steady state).

We conclude that at low concentration α-synuclein helps to accumulate healthy mitochondria. However, high α-synuclein concentration results in accumulation of damaged mitochondria and ROS, and causes the catastrophe.

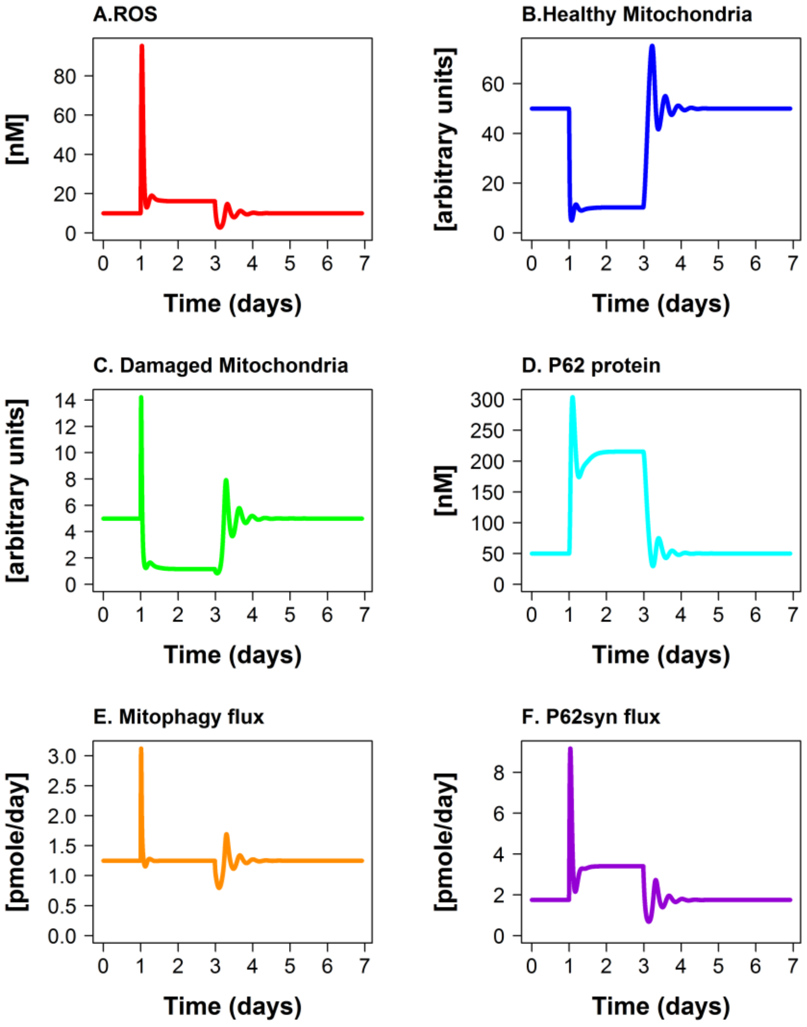

**Figure SM.C.3. Design C (Model C). Response to increase of the ROS generation rate constant (10 fold) in model C without α-synuclein.** Concentrations of ROS and p62 in nM, α-synuclein aggregates, healthy and damaged mitochondria in arbitrary units, fluxes in pmoles/day are shown for model C in response to the change in ROS generation rate constant. The ROS generation rate constant was increased 10 fold on day 1 and decreased 10 fold back to the initial level on day 3.

**The following conclusions were drawn from Figure SM.C.3:**

A) When the ROS generation rate constant is increased on day 1, ROS concentration goes up. After a sharp peak, the ROS concentration decreases again and stays at a level around 60% higher than before perturbation. The observation that ROS concentration increases just 60% while ROS generation rate constant is increased 10 fold suggests a strong ROS homeostasis. When ROS generation is decreased 10 fold, ROS concentration returns to the initial value.
B) When the ROS generation rate constant is increased 10 fold on day 1, the concentration of healthy mitochondria drops 10 fold as well. However, after two damping oscillations, the concentration of healthy mitochondria increases again and stays at a level around 5 fold below the initial level. When ROS generation is decreased 10 fold, the concentration of healthy mitochondria returns to the initial value.

C) Following the perturbation on day 1, the concentration of damaged mitochondria decreases around 4 fold. After two damping oscillations, the concentration of damaged mitochondria increases and stays at value around 5 fold below the initial level. Upon the decrease of ROS generation, the concentration of damaged mitochondria returns to the initial value.
D) The concentration of p62 protein increases substantially (around 4 fold) after the increase of ROS generation on day 1 and returns to the initial level when ROS generation is decreased on day 3.
E) Following the increase of ROS generation, the mitophagy flux increases quickly. However, after short damping oscillations the mitophagy flux returns to the initial level.
F) Levels of p62 increase upon the increase of ROS generation. After short damping oscillations p62 synthesis reaches an elevated level. Upon the decrease of ROS generation, the flux of p62 synthesis returns to the initial steady state level.

We conclude that Model C without α-synuclein exhibits strong homeostasis against 10 fold increase of the ROS generation rate constant.

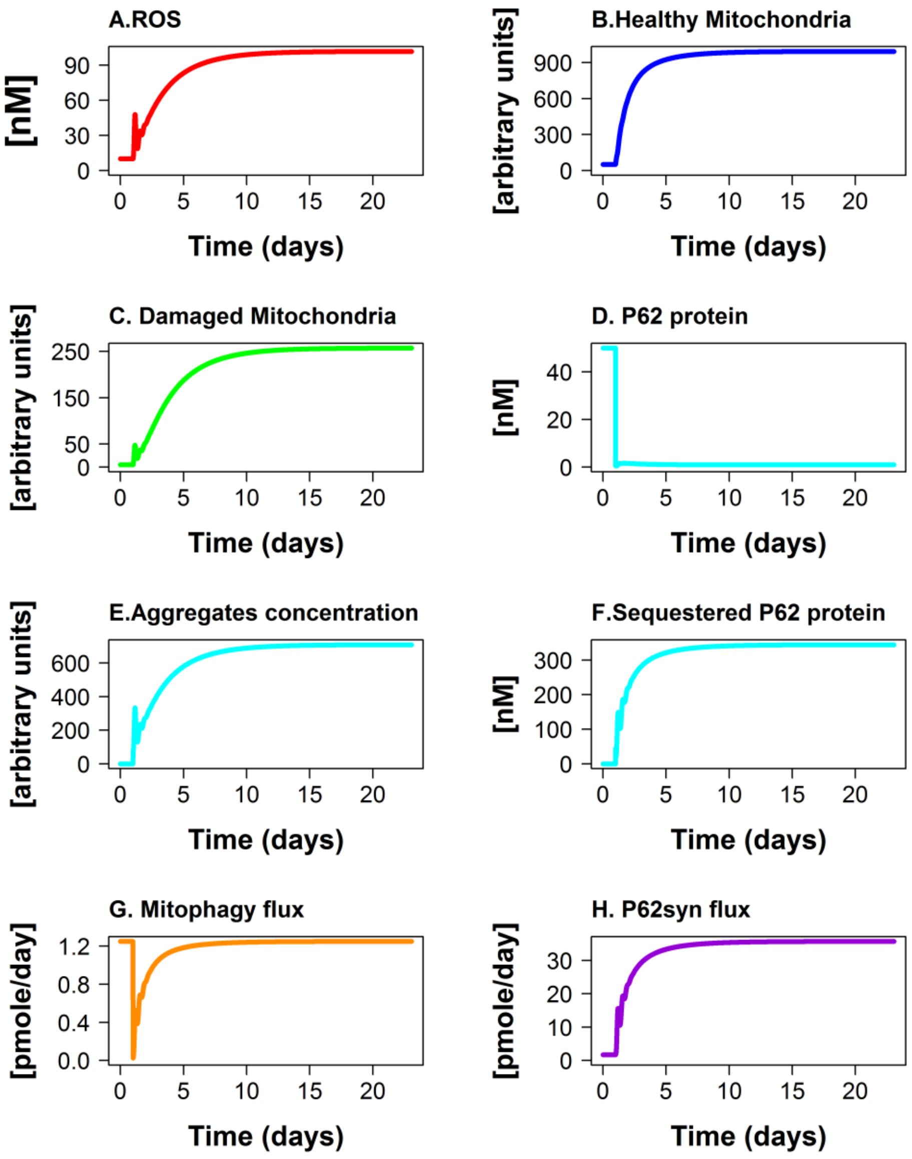

**Figure SM.C.4. Design C (Model C). Response to addition of a constant source of α-synuclein.** Concentrations of ROS (A), p62 (D) and sequestered p62 (F) in nM, healthy (B) and damaged (D) mitochondria and α-synuclein aggregates (E), in arbitrary units, and fluxes of mitophagy (G) and of p62 synthesis (H) in pmoles/day are shown for model C in response to addition of a constant source of α-synuclein on day 1.

**The following conclusions were drawn from Figure SM.C.3:**

A) ROS concentration substantially increases and reaches a new steady state.
B) The concentration of healthy mitochondria substantially increases and reaches a new steady state.
C) The concentration of damaged mitochondria substantially increases and reaches a new steady state.
D) The concentration of p62 substantially decreases and reaches a new steady state.
E) The concentration of aggregates substantially increases and reaches a new steady state.

F) The concentration of sequestered p62 substantially increases and reaches a new steady state.
G) The Mitophagy flux quickly drops, but then, with the increased synthesis of p62, mitophagy returns to the initial level.
H) P62 synthesis flux substantially increases and reaches a new steady state.

We conclude that Model C, upon addition of a constant source of α-synuclein, reaches a new steady state with higher concentration of both healthy mitochondria and ROS.

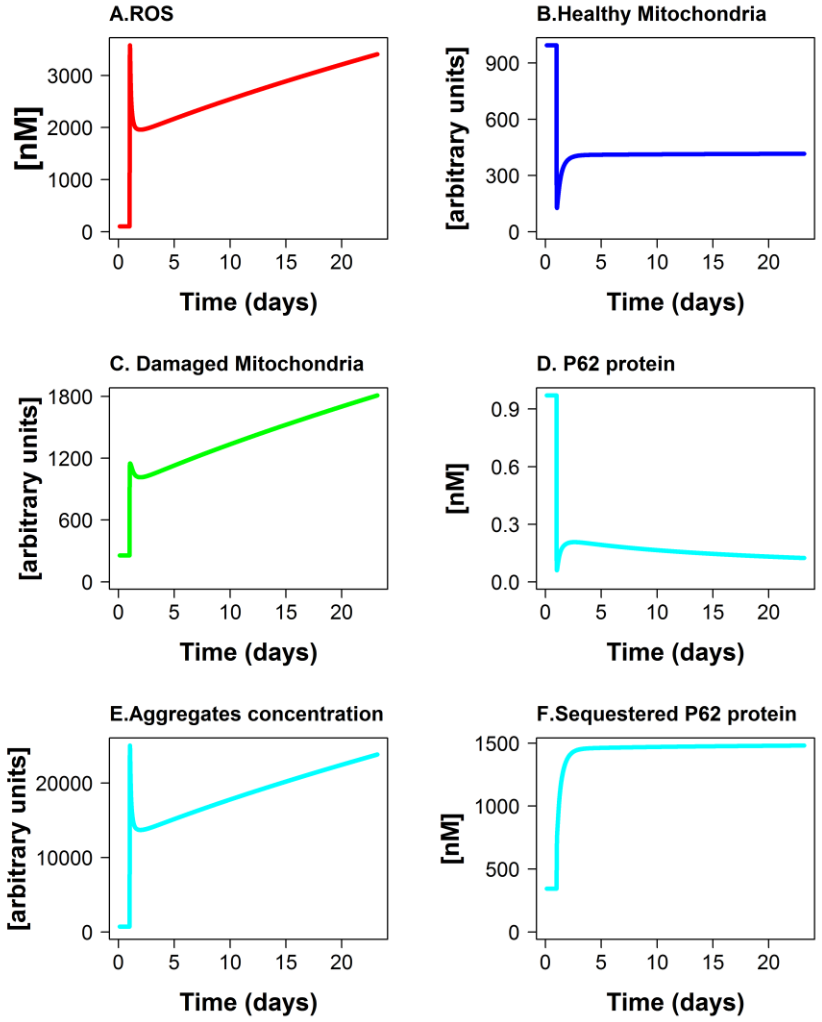

**Figure SM.C.5. Design C (Model C). Response to the increase of ROS generation rate constant in the model with α-synuclein.** Model C with the constant source of α-synuclein was in the steady state found in Figure SM.C.4. On day 1, the ROS generation rate constant was increased 10 fold (similarly to the perturbation described in Figure SM.C.3). The changes in the concentration of ROS (A), p62 (D) and sequestered p62 (F) in nM, healthy (B) and damaged (C) mitochondria and α-synuclein aggregates (E) in arbitrary units were plotted.

**The following conclusions were drawn from Figure SM.C.5:**

A) ROS concentration continuously increases and does not reach steady state. B) The concentration of healthy mitochondria first decreases around 8 fold, but then recovers 3 fold and reaches a new steady state on a value around 2 fold lower than before the increase of ROS generation. C) The concentration of damaged mitochondria continuously increases and does not reach steady state. D) p62 concentration drops 7 fold and reaches very low level (0.1 nM). E) The concentration of aggregates increases and does not reach a steady state. F) The concentration of sequestered p62 increases more than 4 fold and reaches a new steady state.

We conclude that Model C with α-synuclein does not reach a steady state when the ROS generation rate constant is increased.

#### SM.D. Model D fitting to experimental data

| | | mRNA fold changes | | | |
|---|---|---|---|---|---|
| time, min | ROS | Antiox | P62 | Bcl-xl | Nf-κB |
| 0 | 0.53 | 1 | 1 | 1 | 1 |
| 60 | 13 | 1 | 1 | 1.1 | 1.08 |
| 120 | 40 | 1.1 | 1.1 | 0.95 | 1.05 |
| 240 | 75 | 1.2 | 1.6 | 0.83 | 1.12 |
| 360 | 90 | 1.21 | 1.2 | 1 | 1.05 |
| 480 | 90 | 1.25 | 2.2 | 0.85 | 1 |
| 1440 | 20 | 2.2 | 3.2 | 1.02 | 0.63 |

**Table SM.D.T1.** Fold of change in the relative concentrations of ROS and mRNAs for p62, antioxidant response - NQO1 representative gene, Bclxl and NFκB after addition of menadione (100 μM) to HepG2 cells, starting from 1 at the steady state.

#### Model D fitting to experimental data

| $H_2O_2$ | time, min | 0 | 60 | 120 | 180 | 240 | 300 | 360 | 1440 | 2880 | 4320 | 10080 |
|---|---|---|---|---|---|---|---|---|---|---|---|---|
| | time, h | 0 | 1 | 2 | 3 | 4 | 5 | 6 | 24 | 48 | 72 | 168 |
| 50 µM | Sample Continued treat. (% of CTR) | 100 | 81 | 89 | 137 | 96 | 78 | 91 | 49 | 50 | 39 | 27 |
| | Sample Pulse treat. (% of CTR) | 100 | | 128 | | 119 | | 119 | 50 | 45 | 43 | 21 |
| | Sample Repeated treat. (% of CTR) | 100 | | 88 | 59 | 57 | 54 | 28 | | | | |
| 150 µM | Sample Continued treat. (% of CTR) | 100 | 32 | 62 | 60 | 74 | 61 | 83 | 17 | 37 | 21 | 9 |
| | Sample Pulse treat. (% of CTR) | 100 | | 48 | | 65 | | 91 | 35 | 36 | 22 | 7 |
| | Sample Repeated treat. (% of CTR) | 100 | | 19 | 20 | 12 | 10 | 3 | | | | |
| 300 µM | Sample Continued treat. (% of CTR) | 100 | 16 | 38 | 8 | 44 | 12 | 55 | 6 | 20 | 5 | 3 |
| | Sample Pulse treat. (% of CTR) | 100 | | 28 | | 52 | | 65 | 32 | 31 | 16 | 3 |
| | Sample Repeated treat. (% of CTR) | 100 | | 5 | 4 | 2 | 3 | 0 | | | | |

**Table SM.D.T1. Changes in the relative concentration of ATP.** ATP concentration (% of CTR) after either single or periodic addition of $H_2O_2$ to PC12 cells. For single treatments, $H_2O_2$ (50 µM, 150 µM, or 300 µM) was added at 0h. Single treatments were either continued (peroxide was left in the medium and allowed to go through its spontaneous degradation), or pulse (peroxide was washed out after 30 min). For periodic treatments (to simulate chronic oxidative stress) $H_2O_2$ (50 µM, 150 µM, or 300 µM) was added to cells every 1h. Results of single continued treatments were very similar to the results of pulse treatments. Thus, we did not differentiate between pulse and continued treatments during model fitting and call them both as “a single treatment”.

**A**

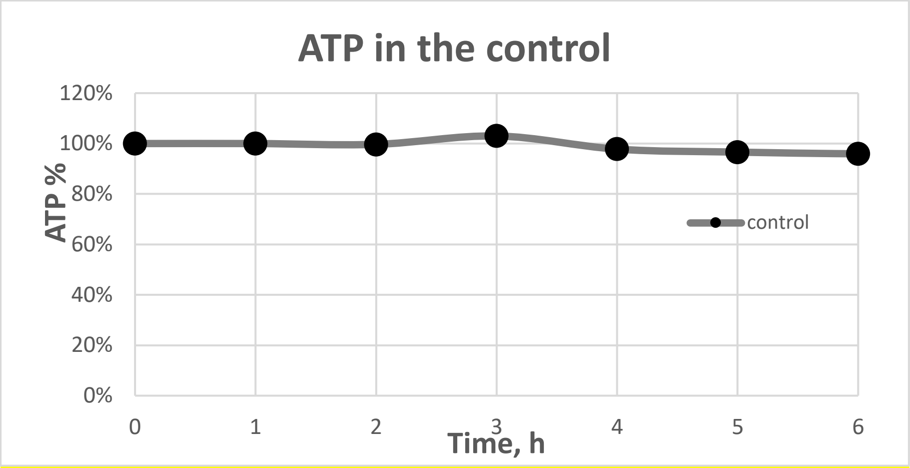

**B**

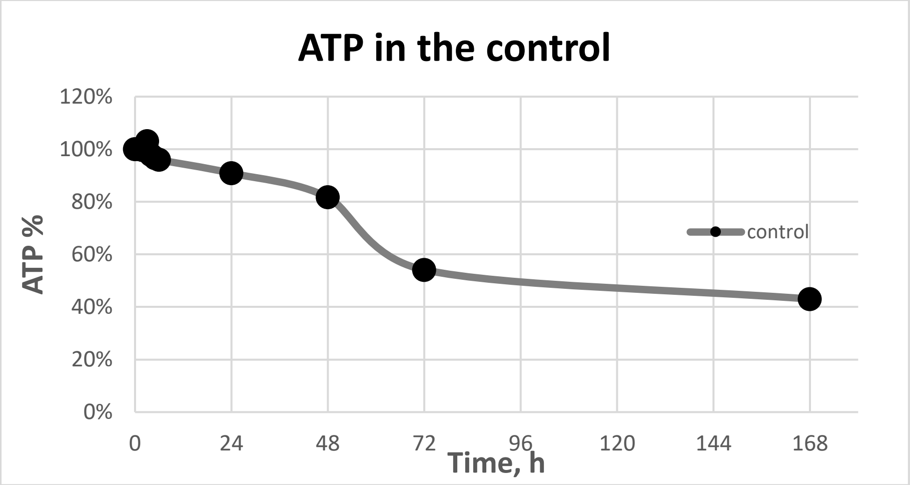

**C**

| | | | | | | | | | | | |
|---|---|---|---|---|---|---|---|---|---|---|---|
| time, min | 0 | 60 | 120 | 180 | 240 | 300 | 360 | 1440 | 2880 | 4320 | 10080 |
| time, h | 0 | 1 | 2 | 3 | 4 | 5 | 6 | 24 | 48 | 72 | 168 |
| control ATP % | 100 | **100** | **100** | **103** | **98** | **97** | **96** | **91** | **82** | **54** | **43** |

**Figure SM.D.1. Changes in control samples in time**. Changes in ATP concentration (in %) in the control plotted in the time scale of first 6 hours (A) and of 7 days (B). In C the data used are presented.

**The following conclusions were drawn from Figure SM.D.1:**

A) ATP concentration in the control in the time scale of first 6 hours does not change.

B) ATP concentration in the control declines with a severe drop between day 2 and 3.

We conclude that the control is stable within the first 6 hours, thus we only used data of the first 6 hours.

**Model D fitting to experimental data**

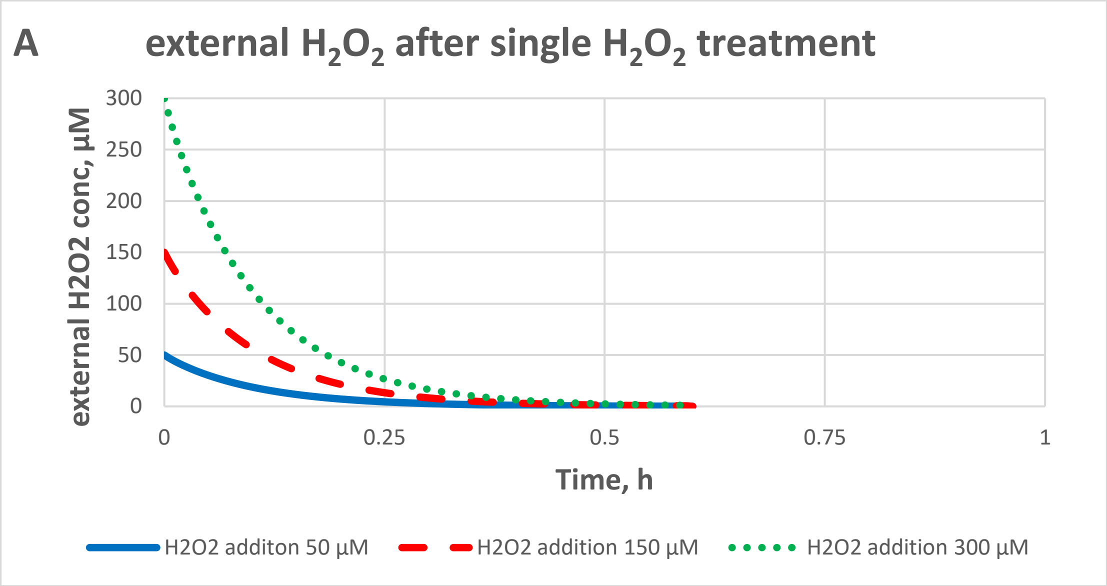

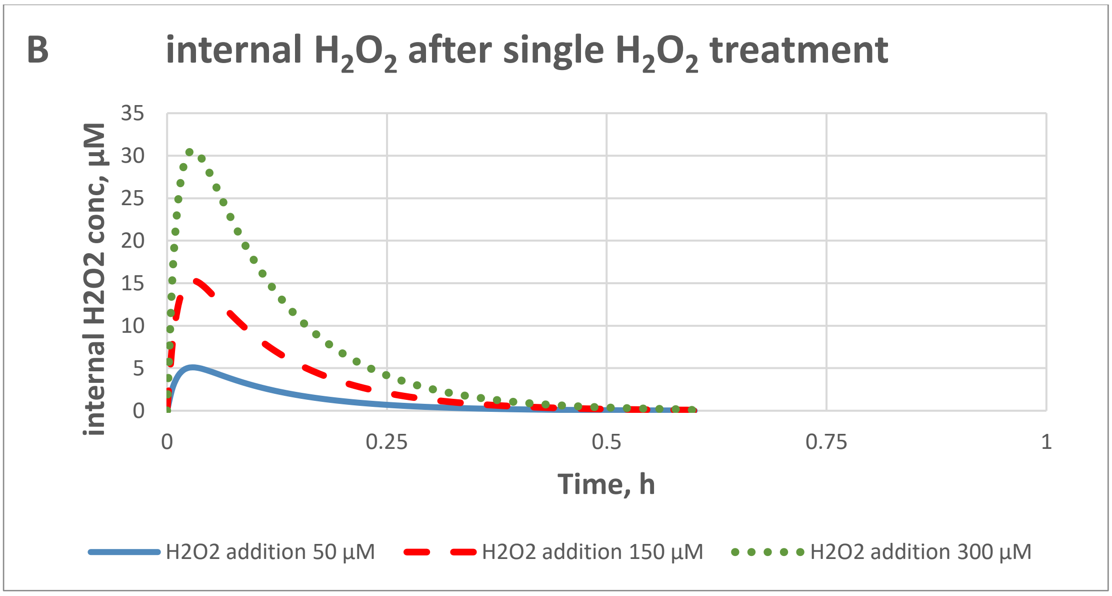

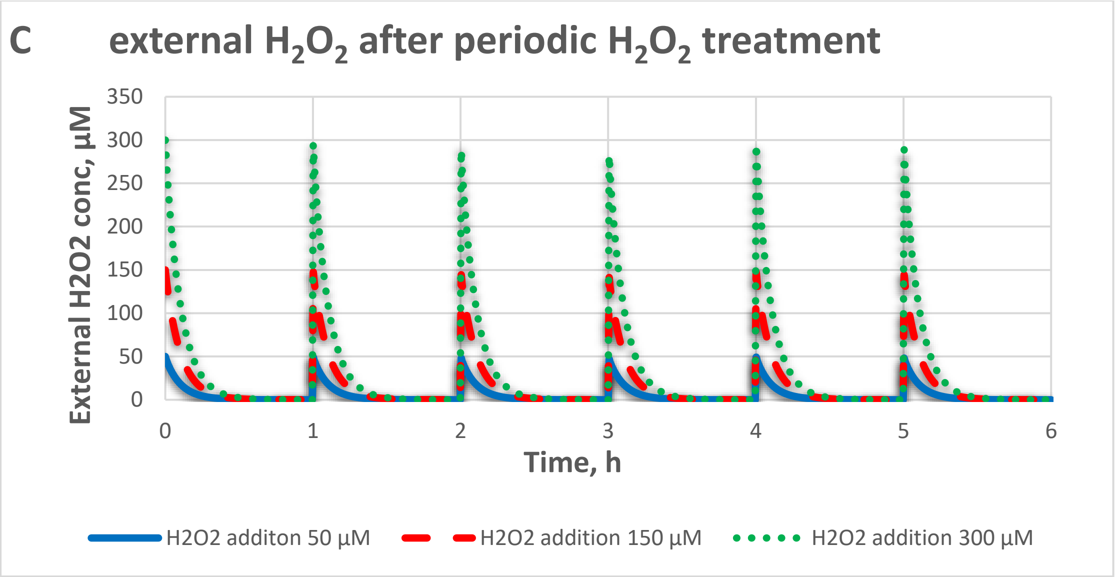
C
external $H_2O_2$ after periodic $H_2O_2$ treatment
External H2O2 conc, µM
350
300
250
200
150
100
50
0
0
1
2
3
4
5
6
Time, h
H2O2 additon 50 µM
H2O2 addition 150 µM
H2O2 addition 300 µM

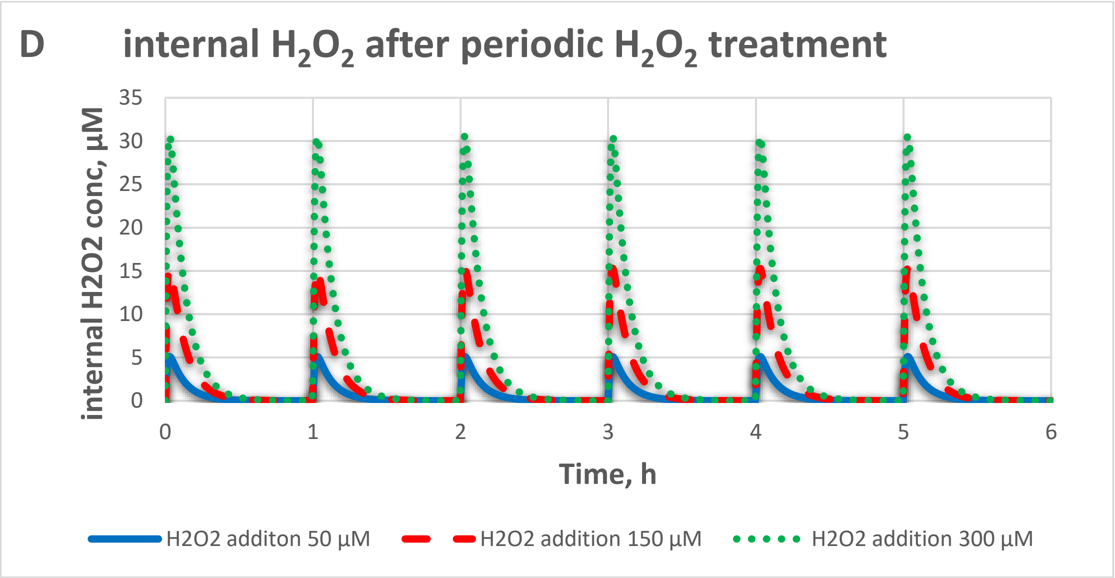
D
internal $H_2O_2$ after periodic $H_2O_2$ treatment
internal H2O2 conc, µM
35
30
25
20
15
10
5
0
0
1
2
3
4
5
6
Time, h
H2O2 additon 50 µM
H2O2 addition 150 µM
H2O2 addition 300 µM

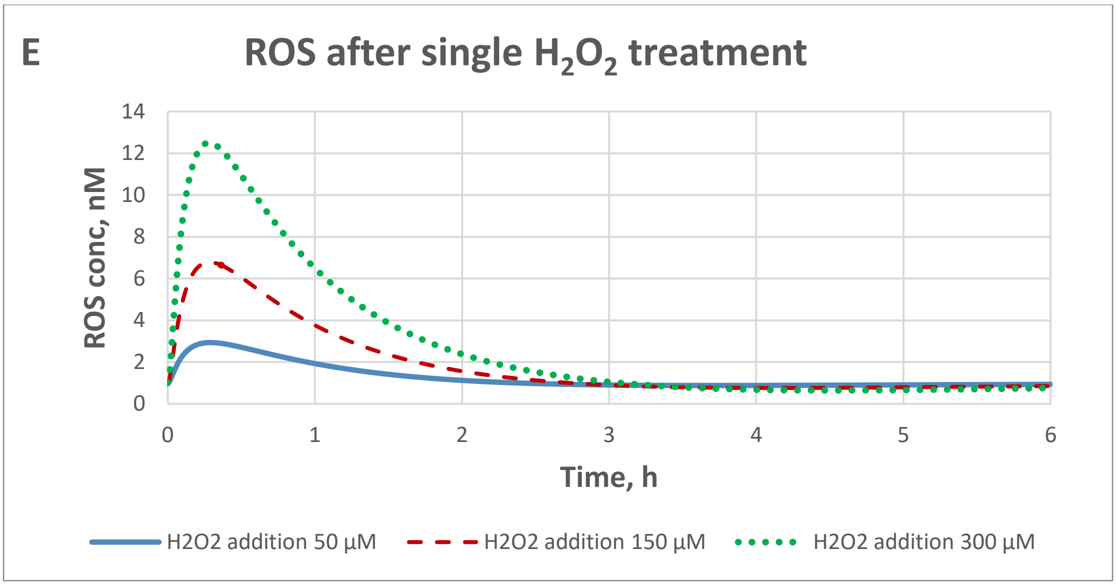

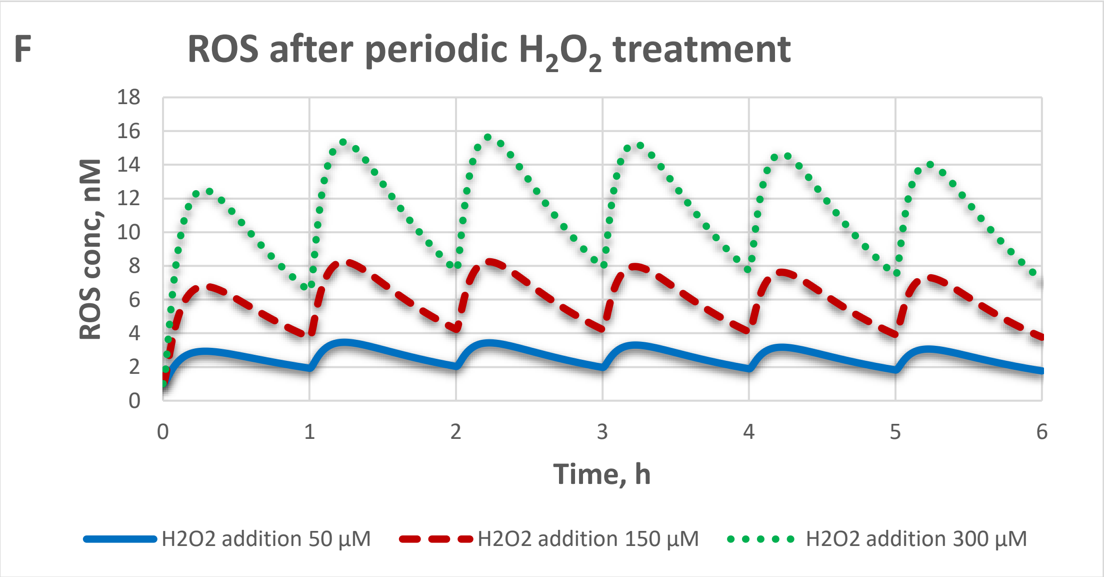

**Figure SM.D.2. Time course of $H_2O_2$ (A-D) and ROS (E-F) in detailed model D.** $H_2O_2$ (50 µM, 150 µM or 300 µM) was added either one time at the time 0h for each single treatment (A, B, E), or every 1h for periodic treatments (C, D, F). Intracellular (internal) and extracellular (external) concentrations of $H_2O_2$ (in µM) and ROS concentrations (in nM) were calculated as described in Material & Methods section.

**The following conclusions were drawn from Figure SM.D.2:**

We conclude that the dynamics of ROS concentrations followed the dynamics of $H_2O_2$ addition with a certain delay.

##### SM.E. Memory in detailed Model D

**A**

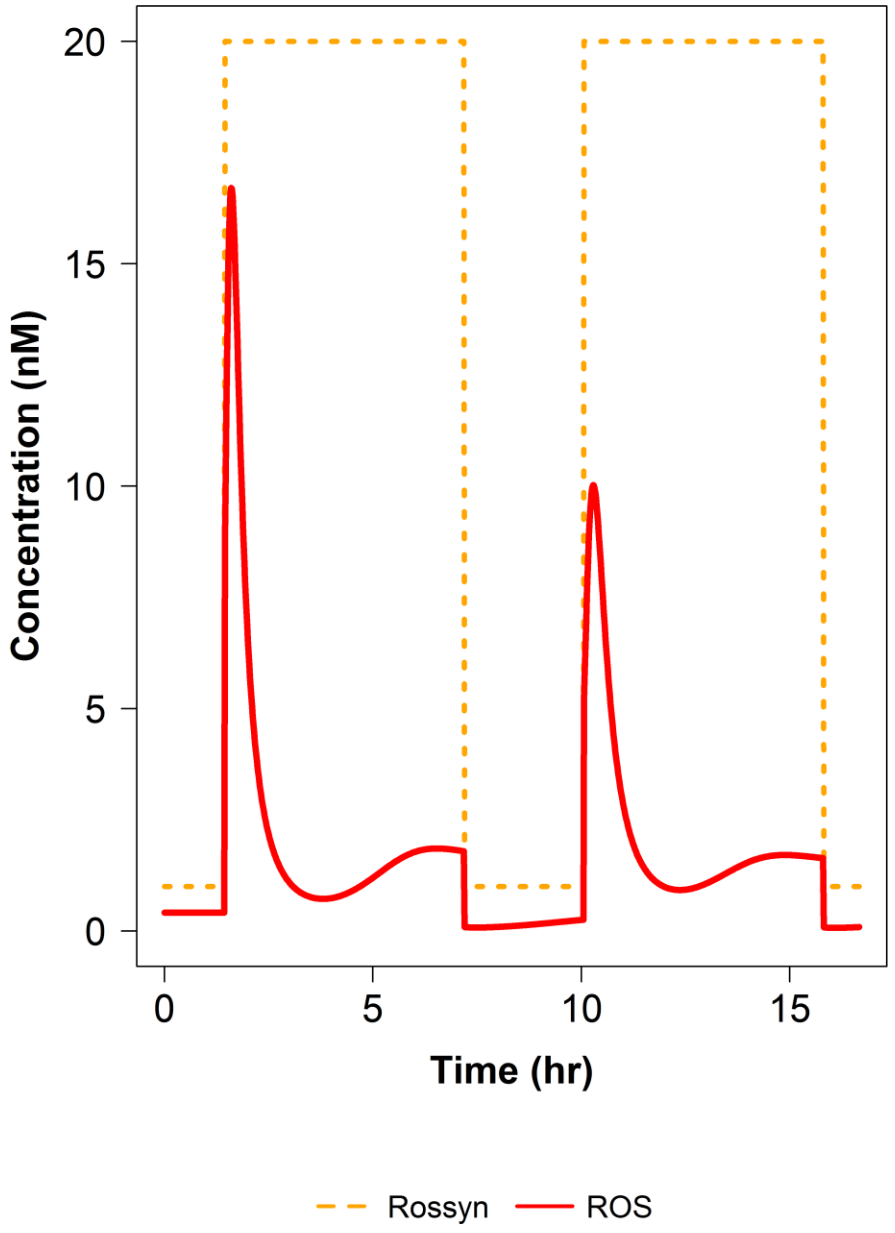

B

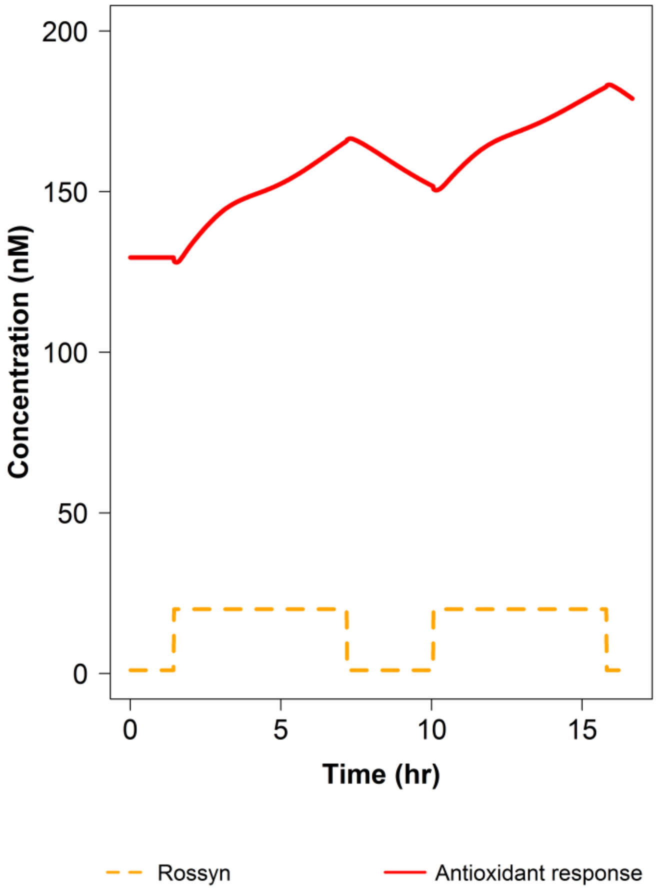

**Figure SM.E.1. Preconditioning to oxidative stress in detailed model (Model D).** The effect of the increased ROS generation rate constant on the concentration of ROS (A) and Antioxidant Response (B) was simulated in detailed model D. We started with the steady state. Then, two pulses of the increased ROS generation rate constant were applied: at 1.47h the ROS generation rate constant was increased 20 fold, at 7.2h decreased 20 fold, at 10h increased 20 fold again, and at 15.8h decreased 20 fold back (dashed orange line).

**The following conclusions were drawn from Figure SM.E.1:**

A) With every pulse of ROS generation, ROS concentration (red line) increases. However, the increase of ROS concentration was higher after the first pulse (37 fold, from 0.45 nM to 16.75 nM) than after the second pulse (22 fold, from 0.45 nM till 10 nM).

B) With every pulse of ROS generation, an antioxidant response (both antioxidant enzymes and antioxidants) changes (red line). After the first pulse of ROS generation, the antioxidant response was activated. At the time of the second pulse, the system has already an elevated level of antioxidant response.

We conclude that the preconditioning caused by the first stress stimuli may enable (train) the system to deal with the consequent stress.

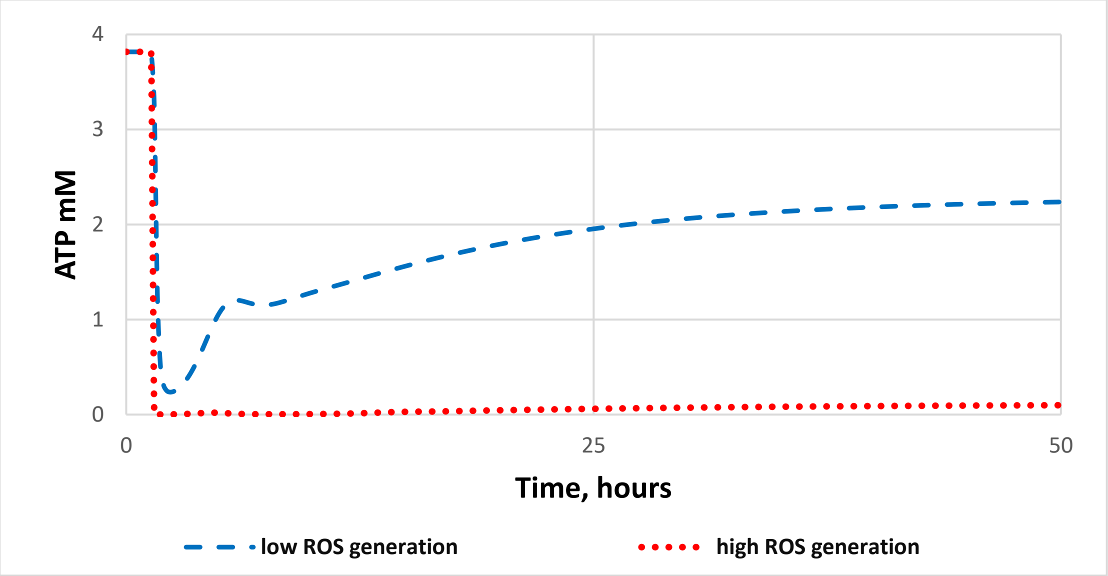

**Figure SM.E.2. Bi-stability in the detailed model D.** Changes in ATP concentration (in mM) was simulated. At the point of 20 year, the ROS generation rate constant was increased either to the smaller (e.g. 20 fold) (dashed blue line) or to the higher extend (e.g. 200 fold) (dotted red line).

**The following conclusions were drawn from Figure SM.E.2:**

At low increase of the ROS generation rate constant, the model exhibits homeostasis and ATP concentration quickly reaches a new steady state.

At higher increase of the ROS generation rate constant, ATP concentration quickly goes to 0.

We conclude that bi-stability is observed. The System is either stabilized or immediately collapsed. There is no gradual deterioration.

#### SM.F. Re-examining 5 design principles in the detailed model

All model and simulation results were assembled into the model archive file called 'SM.F'.

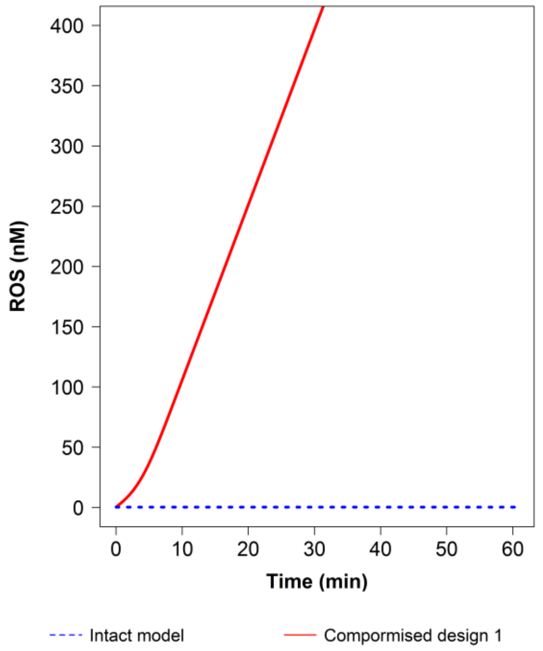

**Figure SM.F.1. The mitophagy and the removal of ROS are required for achieving a steady state (design principle 1).** The change in the concentration of ROS (in nM) was simulated in an intact detailed model D (dashed blue line) and in the model with compromised design principle 1, at which the reactions of ROS removal and mitophagy were deleted (solid red line)

**The following conclusions were drawn from Figure SM.F.1:**

The intact model maintains a steady state. In the model with compromised design principle 1, the concentration of ROS increases to infinity.

We conclude that mitophagy and the removal of ROS are required for achieving a steady state, thereby confirming that the design principle 1, identified first in a simplified model B1, works also in the detailed model D.

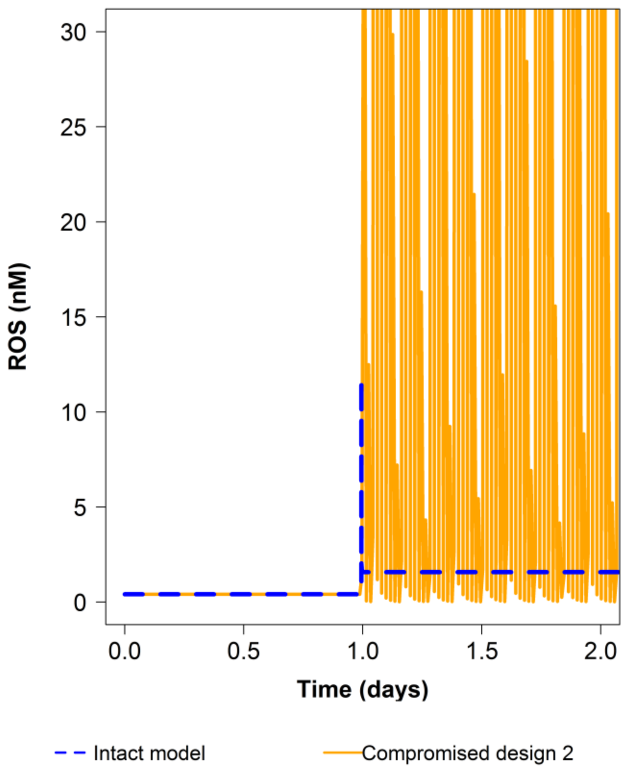

**Figure SM.F.2. Structural robustness is achieved when the concentration of healthy mitochondria is variable (design principle 2).** On day 1, the ROS generation rate constant was increased 50 fold. The change in the concentration of ROS (in nM) was simulated in an intact detailed model D (dashed blue line) and in the model with compromised design principle 2, at which the concentration of healthy mitochondria was constant (orange line).

**The following conclusions were drawn from Figure SM.F.2:**

In the intact model the concentration of ROS reaches a new steady state after a short transient peak. The model with compromised design principle 2 enters into an oscillatory mode.

We conclude that a variable concentration of healthy mitochondria is required to achieve structural robustness, thereby confirming that the design principle 2, identified first in a simplified model B2B, works also in the detailed model D.

**A**

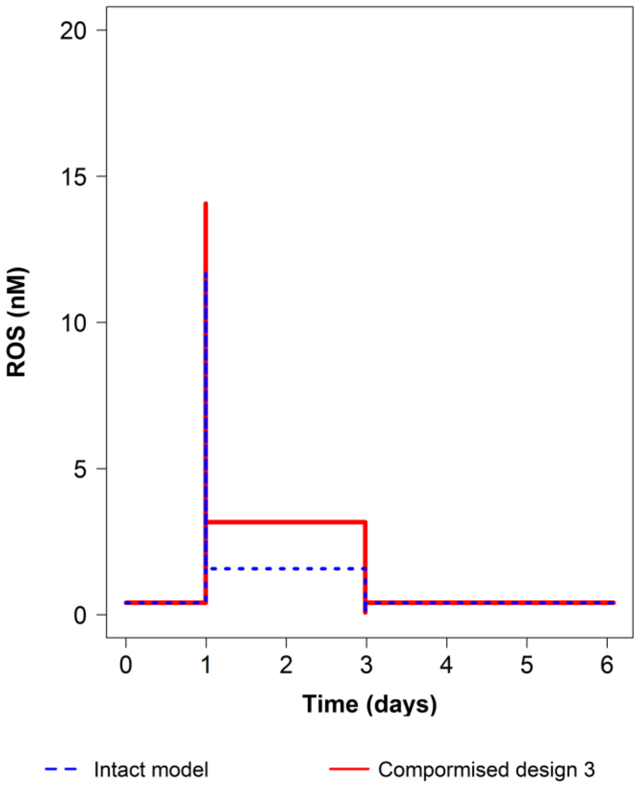

**B**

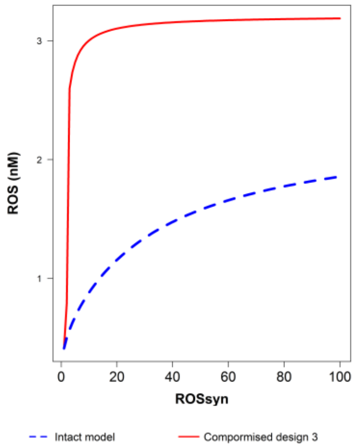

**Figure SM.F.3. Keap1-Nrf2 signaling provides homeostasis (design principle 3).** The change in the concentration of ROS (in nM) was simulated in an intact detailed model D (dashed blue line) and in the model with compromised design principle 3, at which the Keap1-Nrf2 regulation was turned off. In A, the ROS generation rate constant was increased 50 fold on day 1 and returned to the initial value on day 3. In B, the steady state concentration of ROS was plotted versus the ROS generation rate constant.

**The following conclusions were drawn from Figure SM.F.3:**

A) Changes in the ROS concentration in the intact model are less pronounced than in the model without Nrf2-Keap1 signaling.

B) For the same ROS generation rate constants, a steady state concentration of ROS is lower in the intact model comparing to the model where Nrf2-Keap1 signaling is turned off.

We conclude that Nrf2-Keap1 signaling contributes to the strength of ROS homeostasis, thereby confirming that the design principle 2, identified first in a simplified model B2B, works also in the detailed model D.

**A**

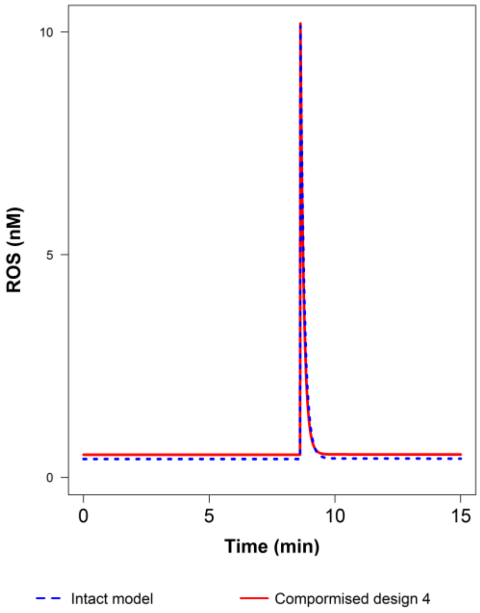

ROS (nM)
10
5
0
0
5
10
15
Time (min)
Intact model
Compormised design 4

**B**

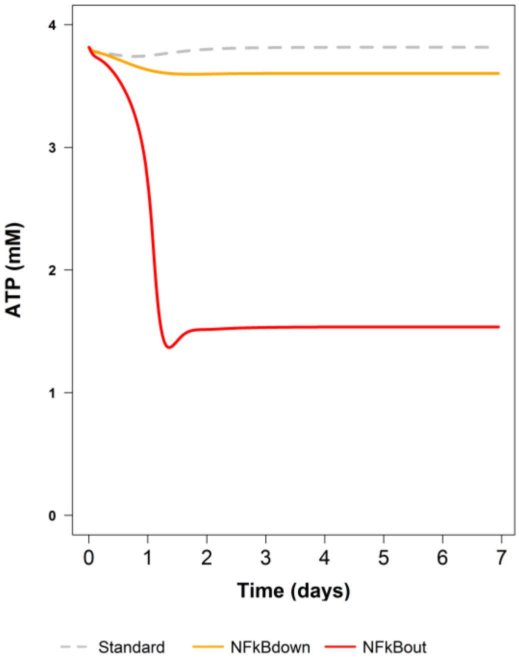

ATP (mM)
4
3
2
1
0
0
1
2
3
4
5
6
7
Time (days)
Standard
NFkBdown
NFkBout

**C**

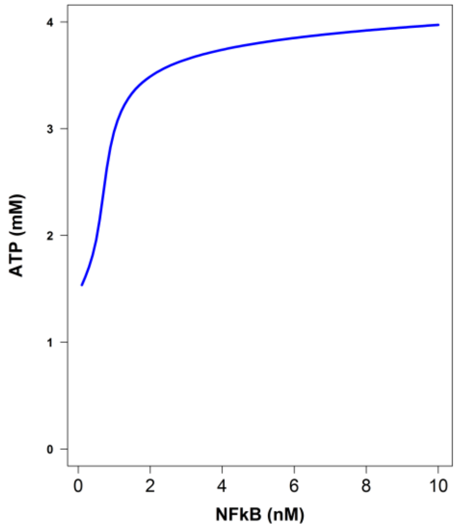

**Figure SM.F.4. The robustness against the perturbations of the initial ROS concentration in the detailed model D (design principle 4).** The changes in the concentration of ROS (in nM) and ATP (in mM) were simulated in the intact detailed model D and in the model with the reduced activity of NFkB signaling. In A, ROS (10 nM) was injected on day 8.6 in the model with (dotted blue line) and in the model without (red line) NFκB signaling. In B, NFκB signaling was knocked down (solid orange line) and knocked out (solid red line). In C, the steady state concentration of ATP was plotted versus the NFκB concentration (that is proportional to the activity of NFκB signaling. The model and the simulation results were assembled into the model archive file called 'SM.F'.

**The following conclusions were drawn from simulation of design principle Figure SM.F.4:**

A) Both in the intact model, and in the model without NFκB signaling, the concentration of ROS increases temporary from 0.4 to 10 nM (25 fold) and quickly returns back to the initial steady state value.

B) The concentration of ATP decreases upon the decrease of NFκB signaling; less pronounced in the case of knockdown (solid orange line) and much more pronounced in the case of knockout (solid red line).

C) The higher is the NFκB signaling, the higher is the steady state concentration of ATP.

We conclude that in the detailed model we are unable to reproduce the design principle 4 in terms of the role of NFκB signaling in the robustness to the ejection of ROS. Nevertheless, NFκB signaling is important to achieve higher ATP concentration. The further role of NFκB signaling in detailed model is reported in Supplemental Materials SM.G.

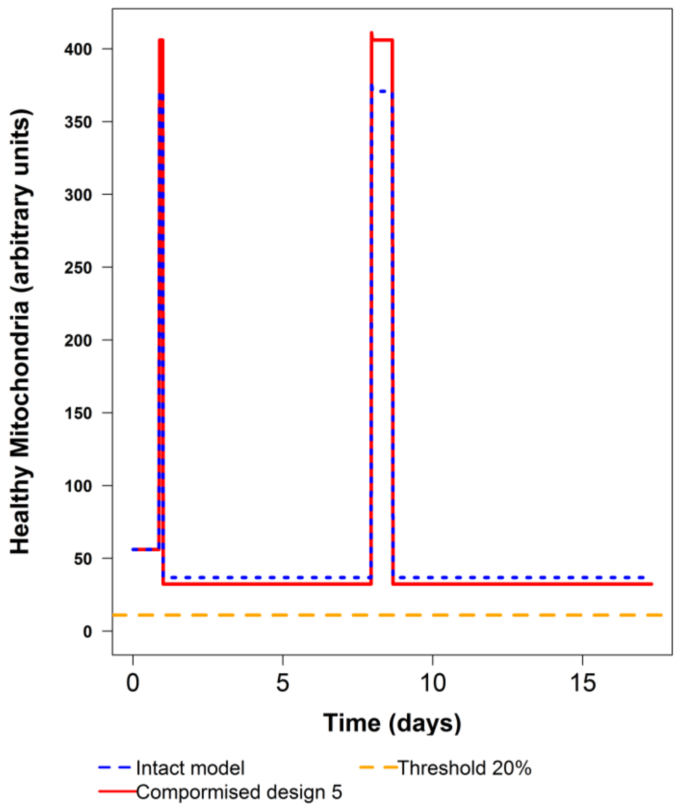

**Figure SM.F.5. Stable homeostasis capable to deal with the second peak of ROS generation in detailed model D (design principle 5).** The changes in the concentration of Healthy Mitochondria (in a.u.) were simulated in the intact detailed model D and in the model with the reduced activity of DJ-1 signaling. Initially, the model was in a steady state. At day 1, the ROS generation rate constant was increased 100 fold, but NFκB signaling was also activated 200 fold, just 3h before the increase of ROS generation. When models reached a new steady state, on day 8, the ROS generation rate constant was decreased 100 fold. When the model reached a new steady state again, on day 8.7, the ROS synthesis was increased 100 fold for the second time.

**The following conclusions were drawn from simulation of design principle Figure SM.F.5:**

Both the intact model and the model with reduced DJ-1 signaling are robust to the second peak of increased ROS generation. In both models the concentration of Healthy Mitochondria does not sweep below a viability line.

We conclude that in the detailed model, the design principle 5 is functioning in the sense that the system is robust to the second pulse of ROS. However, the role of DJ-1 in obtaining robustness is not clear, because other modules contributes to this robustness as well. The further role of DJ-1 signaling in detailed model is studied in Figure 5.

#### SM.G. Aging with compromised NFkB signaling

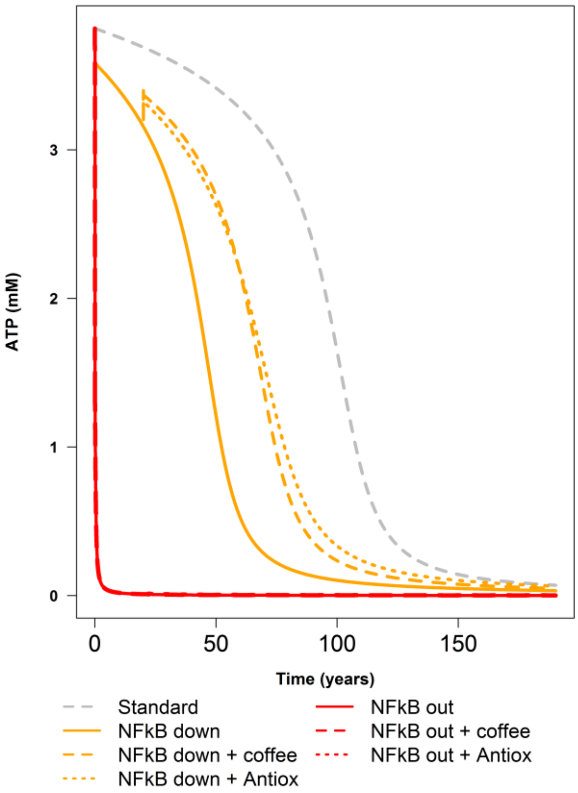

**Figure SM.G.1. ROS induced aging in the detailed model D with compromised NFκB signaling.** The change in the concentration of ATP (in mM) was simulated in the intact model (healthy aging), in the model where NFκB signaling was completely knocked out, and in the model where NFκB signaling was reduced 50% (knocked down). The effects of Nrf2-Keap1 activation (addition of coffee) or addition of antioxidants were simulated as well. The treatment by coffee started at 20 years; it was simulated by activation of Nrf2 nuclear import 1.5 fold. The treatment by antioxidant started at 20 years as well and was simulated as the activation of antioxidants proteins synthesis 1.5 fold.

**The following conclusions were drawn from Figure SM.G.1:**

In the case of healthy aging (dotted grey line), ATP first gradually decreases during the first period of around 75 years. Then the decrease of ATP accelerates. ATP drops below 50% of initial level at around 100 year, and then quickly declines in a non-linear manner.

In the model without NFκB signaling, ATP (solid red line) drops immediately. Treatment with antioxidants (dotted red line) and with caffeine (dashed red line) does not help.

In the model with reduced NFκB signaling, ATP (solid orange line) decreases much earlier than in the standard situation. However, the treatment with antioxidants (orange dotted line) and with caffeine (orange dashed line) helps.

We conclude that the compromise of NFκB signaling accelerates ROS-induced aging. The help from the treatment with both coffee and antioxidants is very limited.

#### SM.H. The effect of α-synuclein on ATP and ROS concentrations in the detailed model

All model and simulation results were assembled into the model archive file called 'SM.H'

**A**

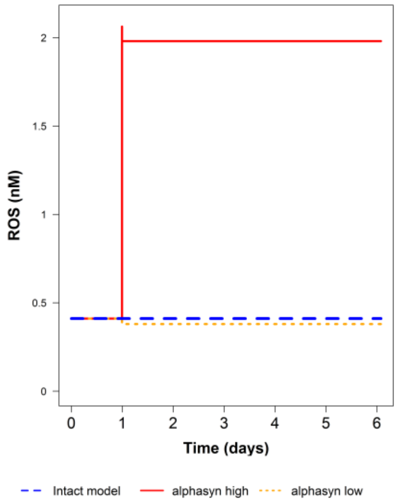

**B**

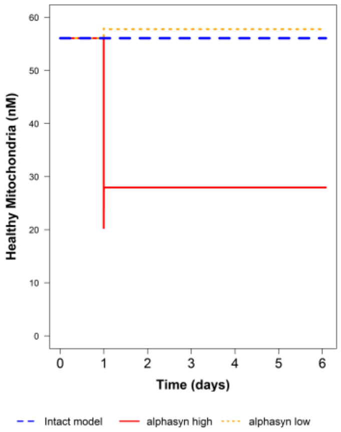

C

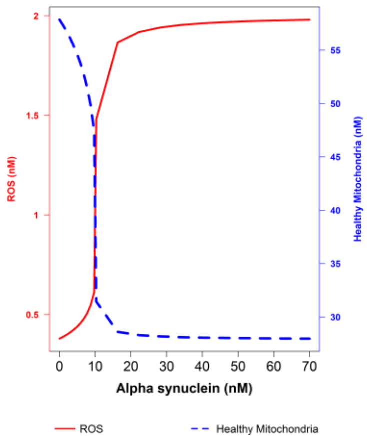

**Figure SM.H.1. The effect of α-synuclein on ATP and ROS concentrations in detailed model.** In A-B, the changes in the concentration of (A) ROS (in nM) and (B) Healthy Mitochondria (in a.u.) were simulated in the intact detailed model D (dashed blue line), in the model where the concentration of α-synuclein source was decreased on day 1 from 3 nM to 0.1 nM (dotted yellow line) and in the model where the concentration of α-synuclein source was increased on day 1 from 3 nM to 70 nM (red line). In C, the steady state concentration of ROS (dashed blue line) and Healthy Mitochondria (red line) were plotted versus the concentration of α-synuclein source.

**The following conclusions were drawn from Figure SM.H.1:**

A) The increase of α-synuclein source results in the increase of ROS concentration.

B) The increase of α-synuclein source results in the decrease of the concentration of Healthy Mitochondria.

C) The higher is the concentration of α-synuclein source, the higher is the concentration of ROS and the lower (up to zero) is the concentration of healthy mitochondria.

We conclude that in the detailed model, α-synuclein causes the increase of ROS similarly to its action in the simplified model C (SM.C). However, in contrast to what was observed in the simplified model C, α-synuclein causes the decrease in the concentration of healthy mitochondria. The further role of α-synuclein is studied in Figure 5.
